# Aggression experience and observation promote shared behavioral and neural changes

**DOI:** 10.1101/2024.12.26.630396

**Authors:** Jorge M. Iravedra-Garcia, Eartha Mae Guthman, Lenca Cuturela, Edgar J. Ocasio-Arce, Jonathan W. Pillow, Annegret L. Falkner

## Abstract

The ability to observe the social behavior of others and use observed information to bias future action is a fundamental building block of social cognition^1,2^. A foundational question is whether social observation and experience engage common circuit mechanisms that enable behavioral change. While classic studies on social learning have shown that aggressive behaviors can be learned through observation^3^, it remains unclear whether aggression observation promotes persistent neural changes that generalize to new contexts. Here, to directly compare the effects of aggression experience and observation at brain-wide scale, we develop a strategy to perform large-scale cell-type specific recordings across subcortical networks for social behavior control and learning. We record longitudinally while animals “train” through direct experience or observation, then probe shared differences in behavior and neural activity in a novel “hard” aggression context. Using supervised and unsupervised methods for behavioral quantification, we detect unique signatures of a shared behavioral strategy not present in animals with no training. During observation, we find widespread activation that mimics experience in networks for behavior generation, with critical differences in signals associated with reward and threat learning. After observation, we observe that changes persist into the novel aggression context, with increased similarity in the neural dynamics between experience and observation groups. Network-level modeling reveals persistent shared changes to a core aggression network, with widespread decoupling of inhibition from a key hypothalamic output region. This demonstrates that “experience-like” activity during observation can recruit a shared plasticity mechanism that biases behavior toward adaptive defensive strategies in new contexts.

---

Humans routinely practice aggression (for example, sparring with a safe partner or in a safe context) in the hopes that it will improve their odds of self-defense in the future. However, it is unclear whether and how aggression experience in safe contexts generates new behavioral policies that generalize to novel uncertain aggression contexts. Can similar benefits be obtained merely by observing these practice sessions? Models of social learning posit that social behaviors, like other behaviors, can be acquired or refined either as a result of either direct experience or observation^2,3^. Behavior may be modified through direct imitation or through emulation of specific behavioral strategies, and that this ability to adapt one’s own future behavior through this indirect experience is a critical building block of social cognition. Albert Bandura’s classic “Bobo doll” experiments^3^, which demonstrated that children can learn aggressive behavior through observation, highlighted that even behaviors that are considered part of an innate social repertoire can be highly influenced by social observation. In humans, exposure to violence is a risk factor that predicts increased likelihood of future violence, suggesting that these effects can be long-lasting and generalize to new contexts^4,5^, but the underlying mechanism remains unclear. In animal models, chronic exposure to aggression has been shown to predict future aggression^6^, and evidence from mice living in groups, where exposure to aggression is both frequent and behaviorally relevant, demonstrates that animals attend to the fights of others by modulating their own aggression levels^7,8^. However, whether experience and observation shape future behavior through a shared neural basis is not known.

While behavior “mirroring” neurons have been described in cortical structures across a variety of species^9^, recent evidence has shown that neurons in a key hypothalamic area for aggression^10^ (the ventromedial hypothalamus, ventrolateral area, VMHvl) may also be recruited during observation of aggression^11^. However, whether this property is widespread within larger subcortical networks for social behavior generation and whether this activity is critical for updating future behavior in novel contexts is unclear. The social behavior network (SBN) is a highly conserved network of largely subcortical brain regions (hypothalamus, amygdala, midbrain) that is conserved across vertebrates and is rich in estrogen and androgen receptors^12^. The SBN is highly interconnected, with many key regions in the “core” aggression network of the hypothalamus and amygdala sending and receiving both excitatory and inhibitory projection to and from other circuit loci ^13,14^. Additionally, this network is tightly connected with networks for appetitive and aversive learning that have been shown to promote and reinforce specific defensive strategies that lead to persistent affective states ^15–17^. These teaching signals may be recruited during observation to represent reward and threat vicariously during complex social behavior^1^, though this has not been demonstrated.

Previous work has implicated several inhibitory projections in the social behavior network as being critical for the gating and generation of aggressive motivational states and actions ^18,19^, yet how the relationship between inhibition and excitation in this circuit changes across experience is unclear. While several previous studies have looked at the synaptic or hormonal basis of aggression experience on a subset of regions within the larger aggression network^20–23^, how these changes influence large scale neural coding across time or are replicated during observational learning has not been explored. One potential hypothesis is that inhibitory networks for aggression control exert less influence after aggression training, leading to differences in aggression and other aggression preparatory behaviors.

Here, to understand whether and how direct repeated aggressive experience (“practice”) and observation can change behavioral strategy and neural dynamics, we track the evolution of neural activity in excitatory and inhibitory populations and dopaminergic release longitudinally across the brainwide social behavior network and test how these changes predict behavior and neural activity in a novel, more difficult aggression context. We find that both aggression experience and observation lead to shared widespread changes in the balance between excitation and inhibition in the social behavior network, with these networks gradually becoming more biased towards excitation with repeated experience or observation. In addition, these changes predict behavioral strategy during a “hard” fight, demonstrating that these changes can update the likelihood of future behavior to promote adaptive self-defense.

## Aggression experience and observation training promote shared behavioral strategies in a novel “hard” aggression context

To test whether aggression experience or observation similarly update aggression-control policies, we tested whether they exhibited shared behavioral strategies in a novel aggression context. We devised a train and test protocol (Fig. 1a) whereby individuals were trained through direct experience, aggression observation, or were provided no previous exposure, then tested by comparing their behavior in a novel aggression context. For the aggression experience group (EXP), males were given opportunities to practice aggression in a safe and easy-to-win context (8 days, 3×10 minute sessions per day), by giving them access to a novel, submissive mouse in their homecage. In the observation group (OBS), mice were placed on a platform above similar aggressive interactions such that they could receive multimodal cues from exposure to this fight, but not interact with the behaving animals. The nonsocial control group (NON) was given access to a mouse toy in their homecage for sessions of matched length. Following the 8-day training regime, all groups were then tested on an identical probe test, the “hard fight”, a novel aggression context with a more difficult opponent. Hard fights consisted of 3 brief (4min) interactions with a similarly sized, aggression-experienced mouse.

**Figure 1:**
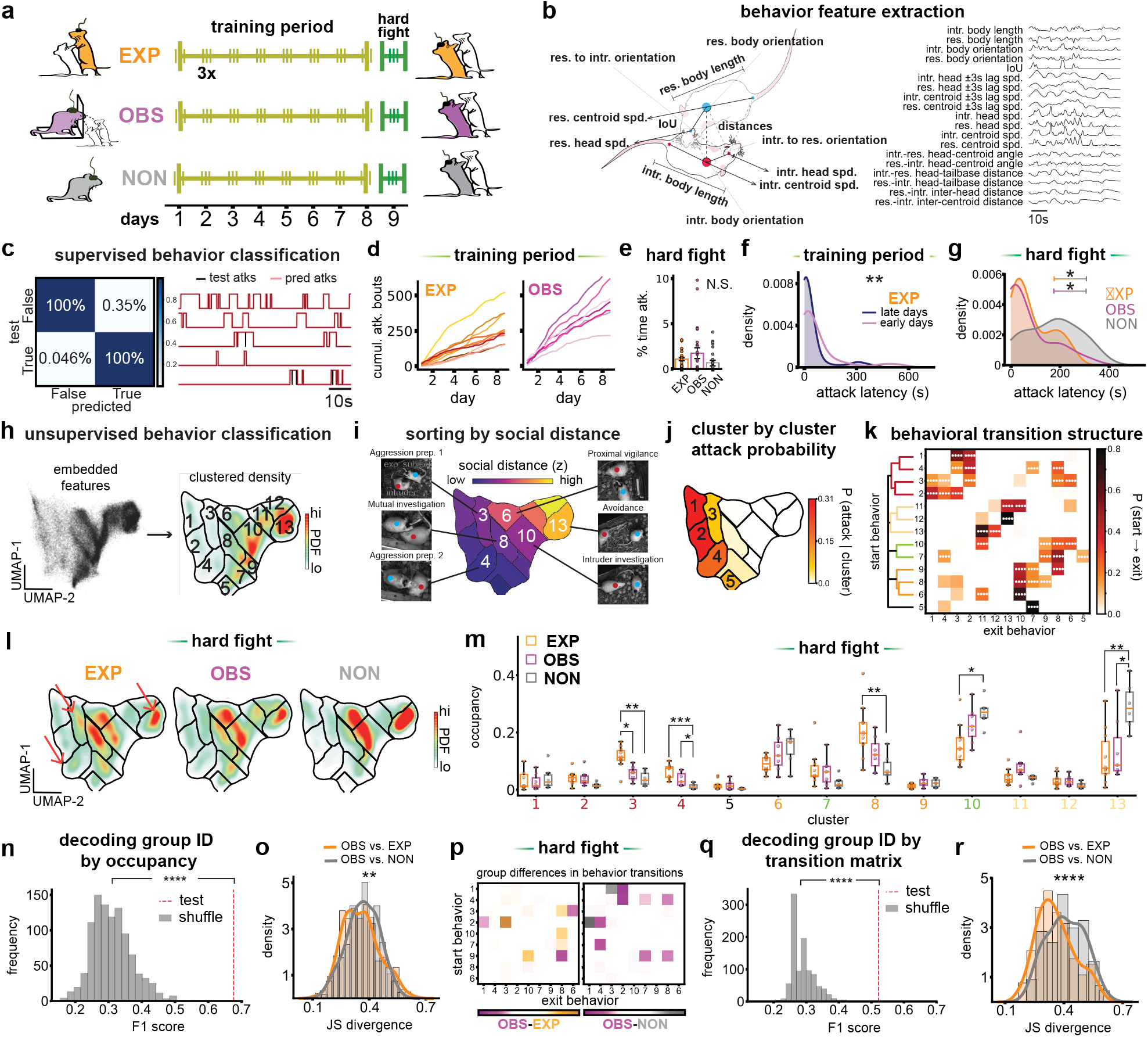
Unsupervised detection of shared changes in behavioral strategy after aggression experience and observation. **a**, Schematic of longitudinal aggression experience, observation or nonsocial experiments. **b**, *Left*: Example of behavior feature calculations from body-part tracking via SLEAP. *Right*: SLEAP-derived features represented as time series. This feature list forms a subset of the entire feature set used for supervised (**c**) behavior classification, in addition to the entire feature set used for unsupervised (**f**) classification. **c**, *Left*: Confusion matrix denoting performance of multilayer perceptron trained on feature data to predict manually (hand) annotated attack behavior. *Right*: Binary prediction of attack behaviors (red) against hand-annotated ground truth (black). **d**, Number of attacks accumulated across every day of the training period for EXP subjects (left, *N* = 10) and OBS subjects (right, *N* = 7). **e**, Average percent of the time spent attacking during the hard fight for all three behavior groups (error bars indicate mean ± SEM, EXP *N* = 30, OBS *N* = 21, NON *N* = 18). **f**, Distributions of first attack latency (s) during first two (pink) versus last two (navy) days of the training period for individuals in EXP group. (early *N* = 60, late *N* = 60, two-sample Kolmogorov Smirnov test). **g**, Distributions of first attack latency (s) during all hard fight sessions for individuals in the EXP (orange, *N* = 30), OBS (purple, *N* = 21) and NON (gray *N* = 18) groups. (two-sample Kolmogorov Smirnov tests with multiple comparisons corrections using FDR method). **h**, *Left*: UMAP embedding from all social interaction sessions. *Right*: Smoothed histogram of the left embedding with clusters numbered via watershed segmentation. **i**, UMAP embedding color-coded by z-scored distance between the experimental subject and the intruder. Video frames depicting example behaviors found within a subset of clusters (3, 4, 8, 6, 10, 13) are shown on the sides of the embedding. **j**, Probability distribution of frames containing attacks found in each UMAP cluster. **k**, Heatmap depicting pairwise transition probabilities of all 13 behavior clusters. The y-axis depicts the pre-transition cluster, and the x-axis depicts the target cluster. Clusters are sorted via unsupervised hierarchical clustering. The resulting dendrogram is visualized with a color threshold of 0.8. Each heatmap cell denotes a pairwise transition probability averaged across social interaction sessions and animal groups. (Pairwise transition probabilities are evaluated for significance via permutation testing. All 156 comparisons are Bonferroni-corrected). **l**, Averaged UMAP embedding for individuals in the EXP (left), OBS (middle) and NON (right) groups during hard fight sessions. **m**, Occupancy (fraction of time relative to entire session) of all behavior clusters across groups during the hard fight. Box plot lines represent the median occupancy, with the box extending from the first to the third quartiles of the data, and the whiskers 1.5x the inter-quartile range (EXP *N* = 10, OBS *N* = 7, NON *N* = 6. Group comparisons done via a one-way ANOVA or Kruskal-Wallis tests pending normality, post-hoc independent samples *t*-tests or Mann-Whitney U tests, and multiple comparison corrections using the FDR method). **n**, Leave-one-session-out cross-validation F1 accuracy (0.68) of group ID (EXP, OBS, NON) predictions using occupancy of UMAP clusters. (Test F1 score is evaluated against a null distribution of F1 scores generated by randomly shuffling group labels for each session). **o**, Distribution of JS divergence values computed between the occupancies of UMAP clusters of every OBS individual against every EXP (orange) or NON (gray) individual (OBS vs. EXP *N* = 630, OBS vs. NON *N* = 378, two-sample Kolmogorov-Smirnov test). **p**, Heatmaps depicting the average difference in pairwise transition probabilities for social clusters (1, 4, 3, 2, 10, 7, 9, 8, 6) between OBS and EXP individuals (left), and OBS and NON individuals (right). Orange cells reflect pairwise transition probabilities higher for the EXP group, while gray or purple cells reflect the same for the NON and OBS groups, respectively. **q**, Same as (**n**) but using pairwise transition probabilities of social UMAP clusters (F1 score of 0.52). **o**, Same as (**o**) but JS divergence values are computed using the pairwise transition probabilities of social UMAP clusters. For all comparisons, \**P*<0.05, \*\**P*<0.01, \*\*\**P*<0.001, \*\*\*\**P*<0.0001. For detailed statistics, see TableS1.

To quantify differences in behavior across training and during the hard fight, we tracked the pose of all individuals and converted pose to a series of individual and social behavioral features (Fig. 1b)^15,24^. We deliberately and exclusively used a top-down camera to quantify behavior, since this is reflective of what the observers viewed from their raised platform. We first trained a supervised behavioral classifier on tracked pose to detect frames attack across the training dataset (Fig. 1c) and evaluated with a held-out dataset. Though the number of attacks in the EXP and OBS group increased across training (Fig. 1d), this did not result in a significant difference in opponent-directed attack across training groups in the hard fight (Fig 1e, Extended Fig. 1a-b). This indicates that neither experience or exposure leads to runaway aggression phenotypes in new contexts, and instead may alter other behavioral strategy or timing. In support of this, we observed that in the experience group, attack latencies decrease from the early days to the late days of the training regime (Fig. 1f), and during the hard fight, but EXP and OBS groups had significantly lower attack latencies than the nonsocial control group (Fig. 1g).

To detect differences in defensive strategy beyond attack, we performed unsupervised clustering of behavior across training and the hard fight^15,25^. To generate a common behavioral embedding (Fig 1h), we performed non-linear dimensionality reduction of behavioral features, then performed density-based clustering of this low-dimensional behavioral embedding, and finally then projected subsets of data into this embedding by group. Across all data, clusters could be mapped to interpretable behaviors, including a series of aggression preparatory postures, investigative behaviors, as well as distal vigilant and attentive states (Fig. 1i-j, Extended Fig. 1c-d). We found that several clusters defined by their low proximity between the animals contained attack and a number of aggression preparatory behaviors, while more distal clusters represented timepoints of social disengagement. Mapping the transition structure between clusters revealed an aggression supercluster, with an increased likelihood of transition between aggression and aggression preparatory behavior (Fig. 1j-k). To directly compare behavioral differences between aggression experience, observation, and nonsocial cohorts, we compared differences in both the cluster occupancy (Fig. 1l-o) and behavior transition matrix (Fig. 1p-r) during the hard fight. We found significant group differences across a subset of behavior clusters and in the raw features that contribute to these clusters (Extended Fig. 1d) that indicated shared changes to an aggression-control policy. Both aggression experienced and observation individuals spend significantly more time in several key aggression preparatory clusters and less time in the inattentive cluster. In addition, experienced animals also show increased time in proactive intruder investigation and decreased passive response to opponent investigation, relative to the nonsocial group. Group identity can be significantly decoded from cluster occupancy (Fig. 1n) or from the transition matrix (Fig. 1q), suggesting the uniqueness of each group’s behavioral strategy. However, when we asked whether behavior in OBS individual animals was more “EXP-like” or “NON-like”, we found that overall behavioral similarity was greater in cluster occupancy, persistence (Fig. 1o, Extended Fig. 1e-g), and in the behavior transition matrix (Fig. 1p,r), particularly in the aggression supercluster (Fig. 1p). Critically, this data suggests that both aggression experience and observation can bias behavior towards similar strategies in a novel aggression context.

## Longitudinal multi-region recordings of excitatory, inhibitory, and dopaminergic signals across aggression experience and observation

Next, to understand whether changes in the activity of the social behavior network across aggression experience and observation could potentially facilitate shared behavioral changes, we developed a 2-color multifiber recording strategy inspired by previous methods for use in freely behaving animals^26,27^. This allows us to record neural activity longitudinally at a network-wide scale (Fig 2a) across training and the hard fight. We targeted an extended social behavior network that included regions in the hypothalamus (ventromedial hypothalamus ventrolateral area (VMHvl), ventral premammillary are (PMv), medial preoptic area (MPOA), and anterior hypothalamus (AH)), amygdala and extended amygdala (medial amygdala (MeA), posterior amygdala (PA), bed nucleus of the stria terminalis (BNST)), midbrain (lateral periaqueductal gray (lPAG)), and regions involved in social valence processing and learning (lateral habenular (LHb), prelimbic area of prefrontal cortex (PrL), ventral lateral septum (vLS), and dopamine release into the nucleus accumbens (NAc-DA)), by implanting custom thin fiber arrays over these targets. Since previous work has implicated inhibitory projections in a core aggression circuit as a gating mechanism for aggression preparation and action, we developed a strategy to map both putative excitatory and inhibitory signals simultaneously at each node in the network (Fig 2b). We used a vgat-ires-cre mouse to target putative inhibitory neurons (vgat+) and putative excitatory populations (vgat-) using a dual-color viral strategy, and also recorded dopamine release in the NAc using a genetically-coded DA sensor (Extended Fig. 2)^28^. We used a custom multifiber isosbestic regression strategy for motion correction in the fighting animal (Fig. 2c) and compared the processed signal across the 3 groups during the training phase (Fig. 2d-g) and during the hard fight.

**Figure 2:**
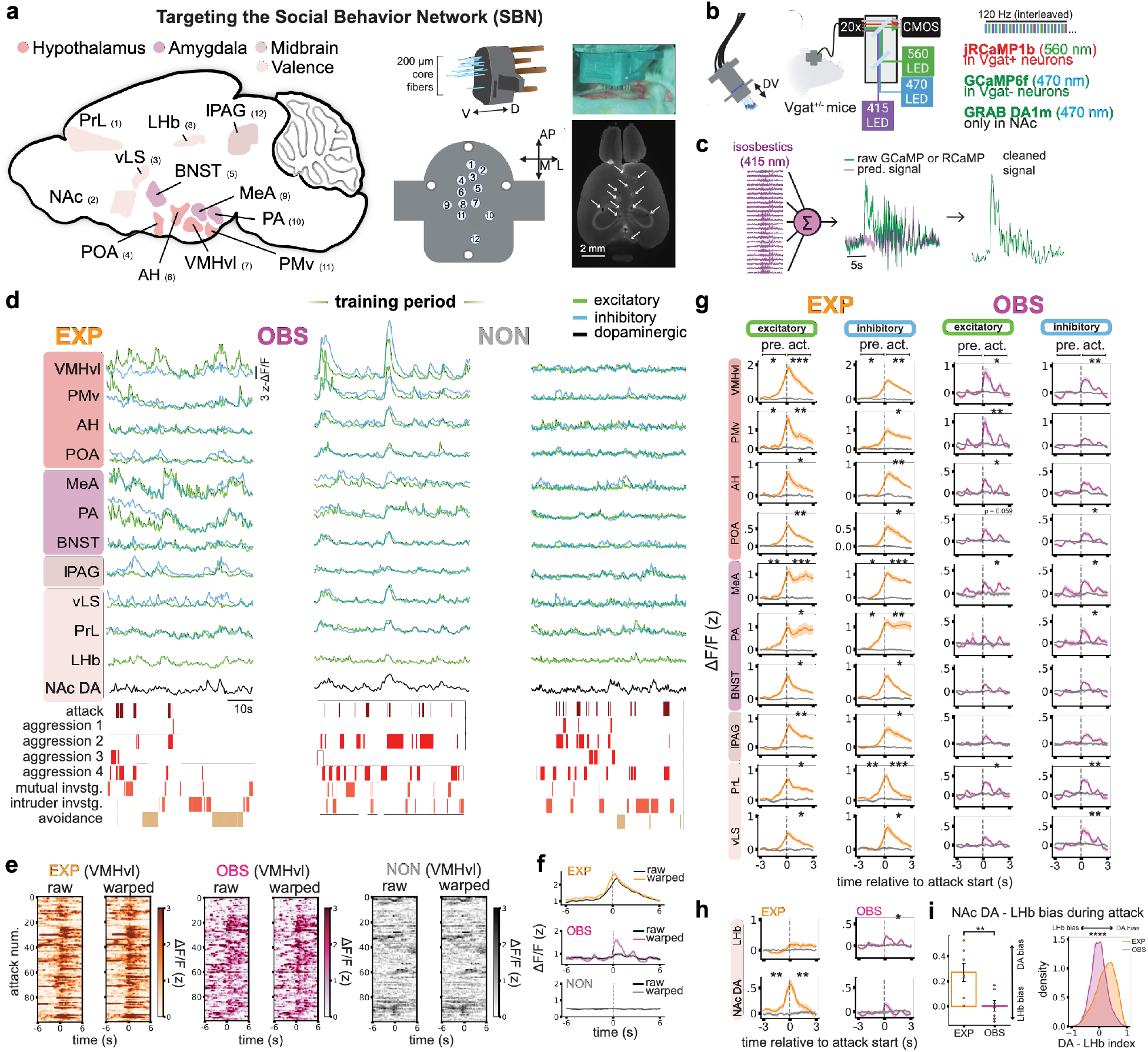
Multi-Region recording of excitatory, inhibitory and dopaminergic networks across aggression experience and observation. **a**, *Left:* Illustration showing recorded regions of the social behavior network (SBN) in the mouse hypothalamus, amygdala, midbrain and valence areas. *Middle:* Side and top views of custom-designed 12-fiber implants for targeting of the SBN. *Right:* Photo of implantation, with top-down light-sheet histology of all 12 fibers delivered into the brain of a sample mouse. **b**, *Left:* Schematic of implant alignment with branching 12-fiber patch cord affixed to custom-designed connector. *Middle:* Three-color imaging strategy in Vgat^+/−^ mice, with 560nm and 470nm LED stimulation for RCaMP- and GCaMP-infused neural populations, coupled with 415nm calcium-insensitive stimulation for motion artifact corrections. *Right:* Custom viral strategy overview. Each brain area except for the NAc receives an infusion of a cre-in jRCaMP1b sensor (expressed in Vgat+ cells) and a cre-out GCaMP6f sensor (expressed in Vgat-cells). NAc only receives an infusion of a GRAB DA1m sensor for recording of dopaminergic activity. **c**, Strategy for signal correction. For each session, isosbestic signals from every region of interest are fitted individually to each GCaMP and RCaMP signal via regression, and the resulting predicted signal is used to subtract the real signal. **d**, Example recordings of multi-site and two-color calcium imaging in EXP (orange), OBS (purple) and NON (gray) sessions during the training period. Excitatory signals are shown in green, inhibitory in blue and dopaminergic in black. Neural signals are aligned to supervised and unsupervised behavior labels (attack labels are derived from supervised classification, whereas aggression 1-4, mutual investigation, intruder investigation and avoidance labels are derived from unsupervised classification). **e**, Example heatmaps depicting unwarped versus time-warped VMHvl activity (z-ΔF/F) during attack windows in representative animals from the EXP (left), OBS (middle) and NON (right) groups. **f**, Mean unwarped or time-warped peri-event time histogram (PETH) of VMHvl activity for EXP (top), OBS (middle) or NON (bottom) groups during the training period. **g**, Training period PETHs of attack-aligned (−3s to +3s) excitatory or inhibitory SBN activity for the EXP (left, orange), OBS (right, purple), and NON (left and right, gray) groups. Activity from each population is normalized by the median activity of the same population 3-5s prior to attack start. In the EXP and OBS groups, activities from −1s to 0s of attack initiation (pre.) and from 0s to +1s of attack initiation (act.) in each population are evaluated separately for significance against the same time periods of the same population’s activities in the NON group (EXP *N* ranges from 8-12, OBS *N* = 7, *N* = 6. Independent samples *t*-tests or Mann-Whitney U tests followed by corrections to all comparisons made for each group using the FDR method). **h**, Same as (**g**) but for excitatory LHb activity (top) and dopaminergic NAc activity (bottom). **i**, DA – LHb index (NAc DA activity – mean LHb E activity, normalized by the sum of the two, for the time period of −1s to 1s centered around attack start). *Left:* Average DA-LHb index values for individuals in the EXP (*N* = 7) versus OBS (*N* =7) groups. (Data are presented as mean ± SEM. Independent samples *t*-test). *Right:* DA – LHb index distributions for all attack trials in EXP (orange, *N* = 22400) versus OBS (purple, *N* = 22400) individuals. (Two-sample Kolmogorov Smirnov test). For all comparisons, \**P*<0.05, \*\**P*<0.01, \*\*\**P*<0.001, \*\*\*\**P*<0.0001. For detailed statistics, see TableS1.

## Attack recruits network-wide activation during experience and observation

We first tested whether aggression networks were activated by specific behaviors in our three groups. Behavioral alignment for NON controls was yoked to randomly assigned EXP individuals and thus reflected random timepoints. We observed network-wide dynamics across excitatory, inhibitory and dopaminergic signals during aggression experience (EXP) with particularly active patterns in the “core” reciprocally connected hypothalamic (VMHvl, PMv) and amygdala (MeA, PA) aggression regions (Fig. 2d). Surprisingly, we also observe robust activation of similar brain regions in observer individuals, even though they are not interacting and are relatively immobile (Fig. 2d, middle).

In observers we observed extensive trial-to-trial variability in attack-aligned activity, perhaps due to actual uncertainty in the observer about the precise time of attack. To align this signal despite this variability, we performed single trial dynamic time warping across all attack trials in EXP, OBS, and NON groups^29^. Using this method, we observed significant attack-aligned signals in EXP individuals in both excitatory and inhibitory signals in nearly every region recorded, with activity preceding attack in many regions (Fig. 2g, left). One notable exception from this activation pattern in EXP individuals is the excitatory population in the LHb, a brain region critical for aversive learning. In OBS animals, after time-warping, we observe attack-aligned activation in a wide swath of regions, including in the “core” subcortical aggression network (VMHvl, PMv, AH, MeA, PA, Fig 2g, right), but also in regions associated with learning and action monitoring (LHb, vLS, PrL). In many regions, particularly in inhibitory populations, attacks late in training recruited less activity than attacks early in training in both EXP and OBS groups (Extended Fig. 3). Critically, we observed no significant attack-aligned activity in any signal in the NON animals, even after time warping (Fig. 2e-h).

Performing attack increases dopamine release in the NAc^15,30^, while *being* attacked increases the response of LHb excitatory neurons^16^. This likely indicates a difference in the perceived valence of this interaction between the two individuals: one experiences it as rewarding, while the other experiences it as aversive. During observational learning, vicarious reward and punishment signals are predicted to drive learning in the absence of direct reinforcement^3^ and vicarious dopaminergic signals have been observed in rodents observing conspecifics receive food rewards^31^. To test whether OBS animals were experiencing signals of vicarious reward and vicarious punishment that matched the experienced individuals, we compared the response of post-warped attack-aligned NAc-DA and LHb-E on OBS and EXP individuals. Here, in animals performing aggression, we observe attack aligned transient responses in the NAc-DA signal, but not in the LHb. In contrast, in observers, we observed significant attack-aligned responses in the LHb, but not in NAc-DA (Fig. 2h). At the single trial level, we observed that the populations in EXP individuals are highly biased towards NAc-DA response, while OBS are biased towards LHb-E response (Fig. 2i). Intriguingly, this may indicate that OBS animals may primarily learn vicariously through the experience of the individual being attacked.

## Neural profiles during observation and hard fight resemble animals with direct experience

To test whether neural activity during aggression observation mimics the signal during experience, we projected the activity of each signal from the training phase onto the behavior map (Fig. 3a) and then compared the similarity across groups in this signal embedding (Fig. 3b). We compared experience and observation groups and observation and behavior-yoked nonsocial controls. This process allows us to ascertain whether signals during observation are a better “match” to signal during experience better than to a random control. In individual neural-behavior maps, we observe that across nearly all signals, activity is biased in the social clusters relative to the nonsocial clusters, and this property is shared by OBS individuals in several core aggression regions including VMHvl, MeA, and PA (Fig. 3c-d). Socially biased activity in OBS is decreased in sessions later in the training regime (Extended Fig. a-b). We used cosine similarity to compare neural-behavior maps across groups, specifically looking at the similarity between EXP-to-OBS and OBS-to-NON (Fig. 3c-e). Here, we find that during observation, inhibitory signals in OBS animals are more similar to EXP, while excitatory OBS signals are more similar to NON (Fig. 3f), with one notable exception being the PA^20,23^.

**Figure 3:**
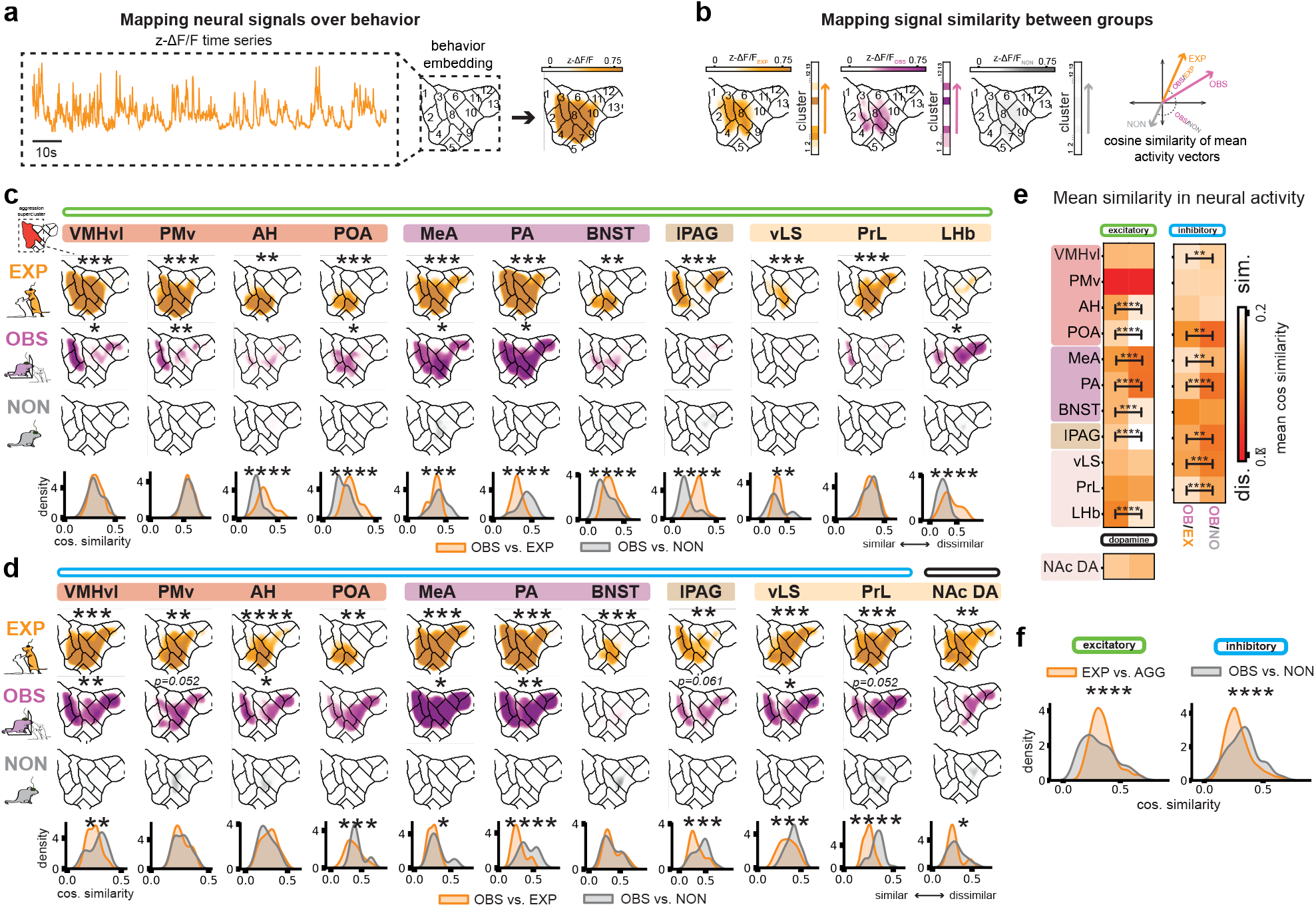
During aggression observation, neural profiles in inhibitory networks are similar to direct experience. **a**, Visualization of population activity across behavior space (activity map). **b**, Strategy for comparing neural activity between groups. For each session and neural population, neural activity is binned into a 13×1 vector where each element corresponds to the mean activity of the population for each UMAP cluster. Similarity in neural activity between groups is quantified as the cosine similarity between mean activity vectors. **c**, *Top:* Excitatory activity maps for each SBN area for EXP (orange), OBS (purple) and NON (gray) animals during the training period. Shown maps represent an average of every map taken from every session across the training period for a representative animal. To evaluate activation of neural populations during aggression clusters, mean activity within these clusters is averaged and subtracted by the mean activity within nonsocial clusters. Mean activity for each population in EXP and OBS animals is compared against that of NON animals for statistical significance. (EXP *N* ranges from 8-12, OBS *N* = 7, *N* = 6. Pending normality, independent samples *t*-tests or Mann-Whitney U tests followed by multiple comparisons corrections to all within-group comparisons using the FDR method). *Bottom:* Cosine similarity distributions between mean activity vectors made up only of social UMAP clusters for every OBS animal and every EXP (orange, *N* ranges from 63-84) or NON (gray, *N* = 42) animal. (Two-sample Kolmogorov-Smirnov test, with all group comparisons corrected using the FDR method). **d**, Same as (**c**) but for inhibitory population activity in the SBN or dopaminergic activity in NAc. **e**, Mean similarities in excitatory and dopaminergic activity (left) and inhibitory activity (right) between every OBS animal and every EXP or NON animal. (Independent samples *t*-tests for Mann-Whitney U tests followed by corrections for all comparisons using the FDR method). **f**, Aggregated distributions for all OBS/EXP (*N* = 777) or OBS/NON (*N* = 463) cosine similarities in excitatory (left) and inhibitory (right) activity. (two-sample Kolmogorov-Smirnov test). For all comparisons, \**P*<0.05, \*\**P*<0.01, \*\*\**P*<0.001, \*\*\*\**P*<0.0001. For detailed statistics, see TableS1.

Next, we asked whether aggression experience and observation lead to more similar neural profiles when animals were presented with a similar behavioral challenge. We predicted that neural similarity between OBS and EXP might be more similar than OBS and NON in the hard fight, even though OBS and NON animals were equally inexperienced. We quantified differences aligned to attack (Fig. 4a-b) and across the whole neural-behavior map (Fig. 4c-f). We find significant group differences in the amount of activity at the time of attack between EXP and NON, particularly in inhibitory populations (VMHvl, PMv, AH, MeA, BNST, PAG). A majority subset of these population differences were also present in OBS animals, particularly in the “core” hypothalamic aggression circuit (PMv, AH) and inputs from the amygdala (MeA, BNST). In addition to map similarity, we also observe group level differences in the tuning to individual behavioral features, suggesting that both experience and observation similarly flatten the encoding of specific features (Extended Figure 5). Using cosine similarity, we found that the neural maps of observer animals more closely resembled the maps of experienced animals rather than the nonsocial animals in both excitatory (Fig. 4c,e), inhibitory (Fig. 4d-e), and dopaminergic signals (Fig. 4c,e), with left-shifted for EXP-OBS comparisons on a majority of individual signals (Fig. 4f). Notably, LHb signal in OBS animals more closely resembled NON animals, suggesting that practice though experience uniquely reduces experienced threat in novel contexts. Overall, these data indicate aggression observation can generate “experience-like” neural profiles at brain-wide scale in addition to promoting “experience-like” behavior.

**Figure 4.**
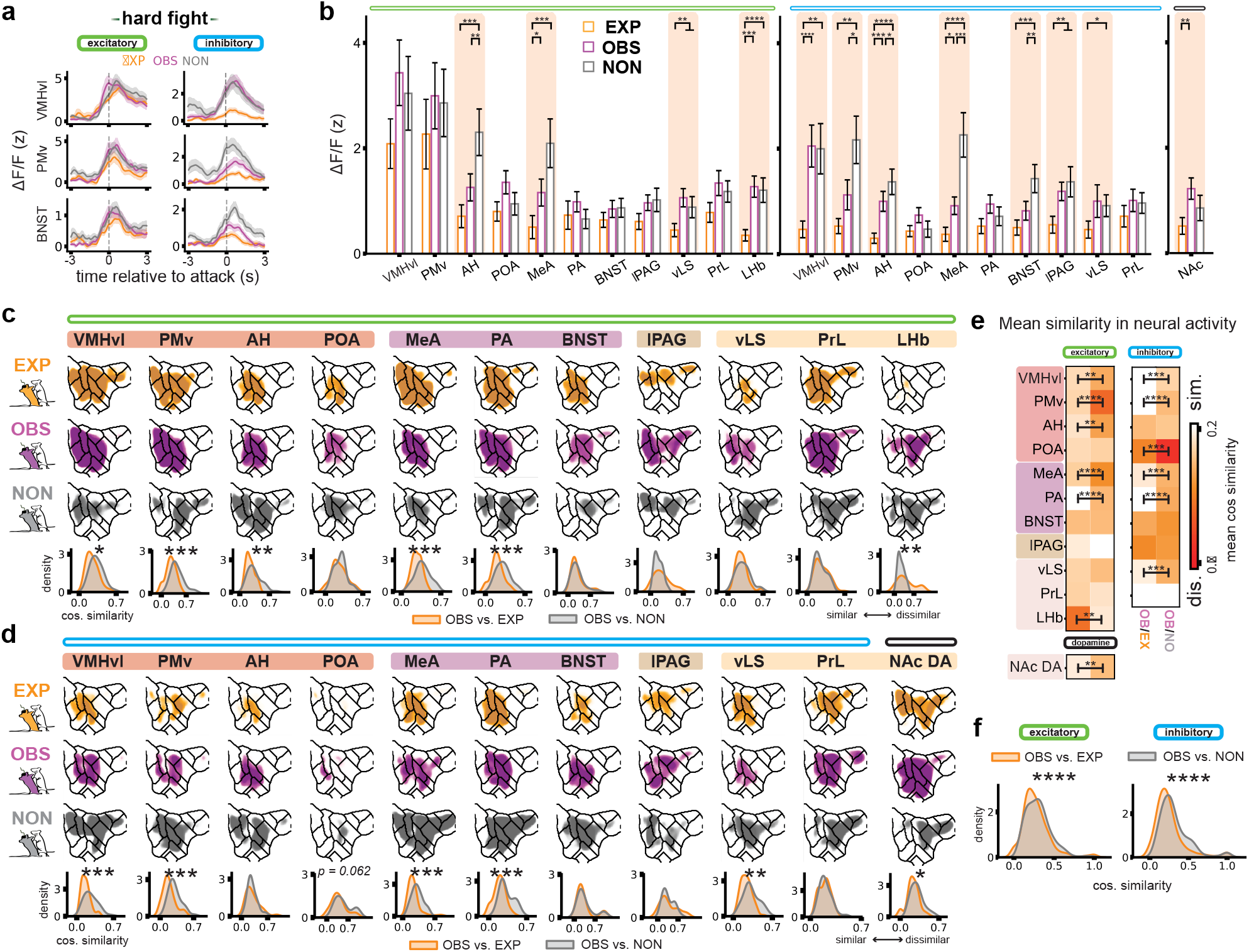
Previous aggression experience and observation promote similar neural profiles during a novel aggression context. **a**, Peri-event time histograms of VMHvl, PMv and BNST excitatory (left) and inhibitory (right) activity for all groups during hard fight. **b**, Average z-scored ΔF/F for −1s to 1s of attack initiation during hard fight for excitatory (left), inhibitory (middle) and dopaminergic (right) populations. (Error bars represent mean z-ΔF/F across all attack windows ± SEM. EXP *N* = 72, OBS *N* = 80, NON *N* = 42. One-way ANOVA or Kruskal-Wallis tests pending normality, followed by post-hoc independence *t*-tests or Mann-Whitney U tests with comparisons corrected using FDR method). **c**, *Top:* Excitatory activity maps for each SBN area for EXP (orange), OBS (purple) and NON (gray) animals during hard fights. Shown maps represent an average of every map taken from every hard fight session for a representative animal. *Bottom:* Cosine similarity distributions between mean activity vectors made up of all UMAP clusters of every OBS animal and every EXP (orange, *N* ranges from 63-84) or NON (gray, *N* = 42) animal. (Two-sample Kolmogorov-Smirnov test, with all group comparisons corrected using the FDR method). **d**, Same as (**c**) but for inhibitory population activity in the SBN or dopaminergic activity in the NAc. **e**, Mean similarities in excitatory and dopaminergic activity (left) and inhibitory activity (right) between every OBS animal and every EXP or NON animal. (Independence *t*-tests or Mann-Whitney U tests pending normality, followed by corrections for all comparisons using the FDR method). **f**, Aggregated distributions for all OBS/EXP or OBS/NON cosine similarities in excitatory (left, *N* = 812) and inhibitory (right, *N* = 462) activity. (two-sample Kolmogorov-Smirnov test). For all comparisons, \**P*<0.05, \*\**P*<0.01, \*\*\**P*<0.001, \*\*\*\**P*<0.0001. For detailed statistics, see TableS1.

## Aggressive experience and observation both result in the uncoupling of inhibition from the core aggression network during training and in the hard fight

We predicted that aggression experience and observation facilitate similar future behavioral strategies through a shared plasticity mechanism that is absent in the nonsocial group. To test this, we next examined whether inhibition in the social behavior network exerts less control over core excitatory networks for aggression by looking at its ability to predict behavior and neural activity. First, we trained a decoder to predict attack on a frame-by-frame basis during training and hard fight using network-wide neural activity. We found that while signals from all groups could robustly predict attack in the hard fight (Fig. 5a), only aggression experience could reliably predict attack during the training, likely due to the temporal jitter in attack encoding in observation animals. Additionally, the ability of models trained on data from the training period to predict attack in the hard fight increases with training (Fig. 5b). Ablating excitatory regions *or* inhibitory regions and examining the corresponding loss in decoder accuracy revealed that in EXP individuals, excitation becomes more important in predicting moment-to-moment attack (Fig. 5c), and this importance emerges over the training epoch (Fig. 5d).

**Figure 5.**
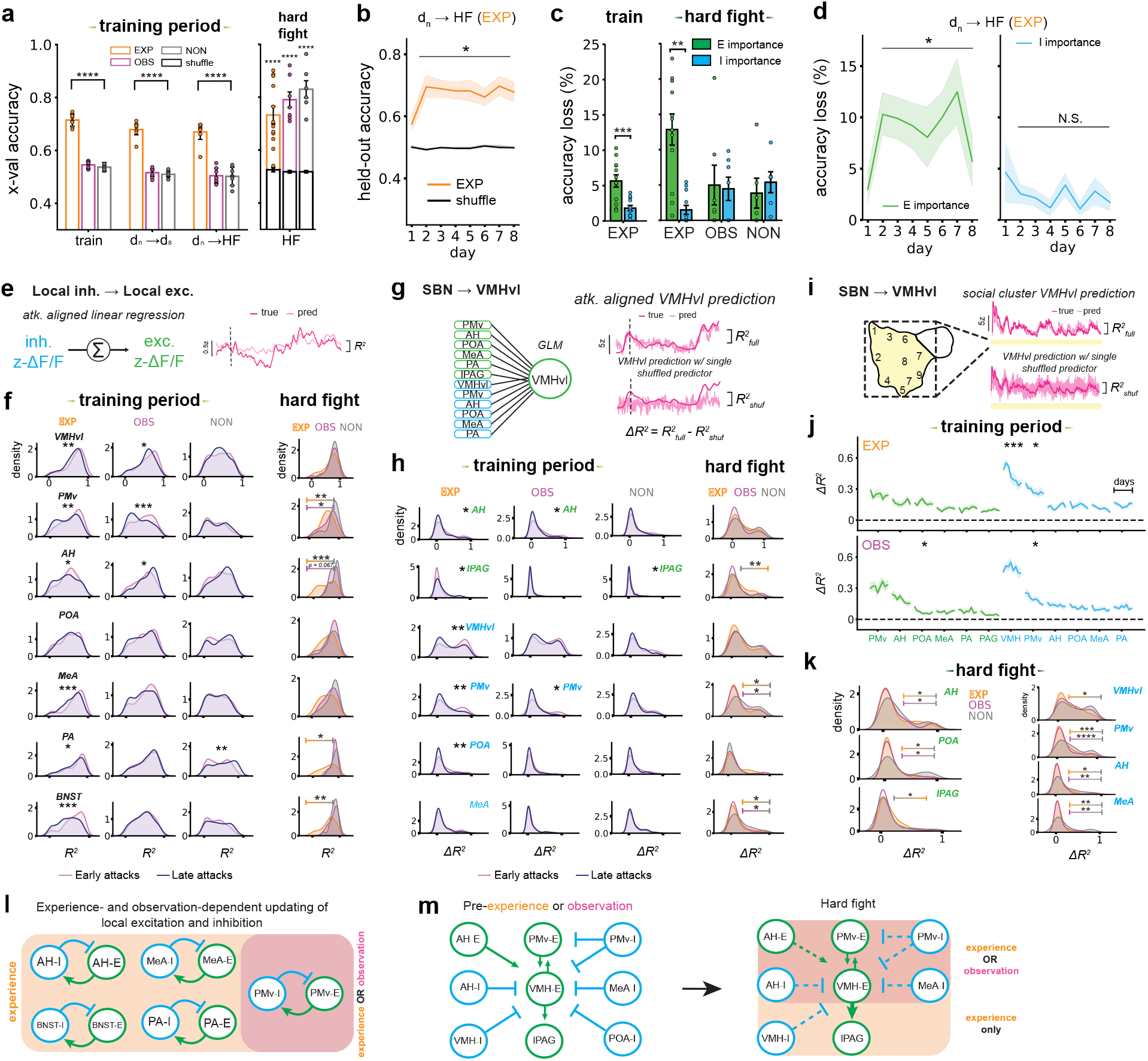
Experience and observation promote widespread decoupling of inhibitory and excitatory aggression networks. **a**, Cross-validated accuracy scores resulting from network-based decoding of attacks. *Left: k*-nearest neighbors classifiers were trained to predict frame-by-frame attack labels for each day of the training period using all SBN signals, and were then tested on 3 held-out sets of data: 1) held-out SBN signals from the same day (train), 2) held-out SBN signals from the 8th day (d_n_ d_8_), and 3) held-out SBN signals from the hard fights (d_n_ HF). (Data are reported as mean accuracy ± SEM. EXP *N* = 12, OBS *N* = 7, NON *N* = 6. Independent samples t-tests with all comparisons FDR-corrected). *Right:* Decoding performance of classifiers trained only on hard fight SBN signals and tested on held-out signals from the same sessions (HF). Null accuracy distributions were generated for validation of each group’s classifiers by training additional classifiers on shuffled SBN signals and testing on the original held-out signals. (Data are reported as mean accuracy ± SEM. EXP *N* = 12, OBS *N* = 7, NON *N* = 6. Independent samples t-tests with all comparisons FDR-corrected). **b**, Held-out accuracy scores for classifiers trained on training period signals from each day to predict hard fight labels for the EXP (*N* = 12) group. (Shaded area represents mean for each day ± SEM. p-value is extracted for the effect of time on classifier accuracy using a linear-mixed effects model). **c**, Loss of decoding accuracy following feature importance testing of excitatory or inhibitory signals. For each day and animal, two separate attack classifiers were trained: one only on excitatory SBN signals, and another on inhibitory signals. The accuracies of excitatory only (I importance, blue) or inhibitory only (E importance, green) models were then compared against the accuracy of the full model as a percentage loss of the latter’s accuracy. *Left:* Averaged E and I importances across the training period for the EXP group. (Error bar represents mean ± SEM. EXP *N* = 12, OBS *N* = 7, NON *N* = 6 Related *t-*test). *Right:* Averaged E and I importances for all groups during hard fight sessions. (Error bars represent mean ± SEM. One-way ANOVA, post-hoc related *t*-tests with corrections for group comparisons using Bonferroni method are performed). **d**, Held-out accuracy loss (%) from inhibitory-only (left) and excitatory-only (right) classifiers trained on training period signals from each day to predict hard fight labels for the EXP (*N* = 12) group. (Shaded area represents mean ± SEM. p-values are extracted for the effect of time on classifier accuracy in separate E-importance or I-importance linear-mixed effects models and are corrected via the FDR method). **e**, Linear regression strategy for mapping correlations between local excitatory and inhibitory signals during attacks. **f**, *Left:* R^2^ distributions of local inhibitory to excitatory signals for a subset of SBN areas during early (first 30, pink) versus late (last 30, navy) attacks for EXP (left), OBS (middle) and NON (right) animals. (Two-sample Kolmogorov-Smirnov tests with all comparisons within each group FDR-corrected). *Right:* R^2^ distributions of the same SBN areas during first hard fight attacks for each group (EXP: orange, OBS: purple, NON: gray). (Two-sample Kolmogorov-Smirnov tests with all group comparisons FDR-corrected). **g**, Multi-region encoding of excitatory VMHvl activity during aggression. Generalized Linear Models (GLMs) were trained on excitatory activity from PMv, AH, POA, MeA, PA and lPAG, along with inhibitory activity from VMHvl, PMv, AH, POA, MeA and PA, to predict excitatory VMHvl activity during each attack. The original fit of these models was saved as R^2full^. Then, each model iteratively predictsed on a feature set with 1/12 predictors shuffled. The resulting fit of these models (R^2shuf^) was used to subtract the original fit (ΔR^2^) which provided a measure of importance of the shuffled predictor in predicting VMHvl activity. **h**, *Left:* ΔR^2^ distributions from GLM fits during early (first 30, pink) versus late (last 30, navy) attacks for EXP (left), OBS (middle) and NON (right) animals. (Two-sample Kolmogorov-Smirnov tests with all comparisons within each group FDR-corrected). *Right:* ΔR^2^ distributions from GLM fits during attacks in hard fight sessions. (Two-sample Kolmogorov-Smirnov tests with all group comparisons FDR-corrected). **i**, Multi-region encoding of VMHvl activity during social behavior. The same strategy is employed as in (**d**) but by aligning neural activity to individual social UMAP clusters rather than discrete attacks. **j**, Average ΔR^2^ for each predictor of VMHvl activity across all 8 days of the training period for EXP (top) and OBS (bottom) animals. (Shaded area represents the mean ΔR^2^ from GLM fits across each of 10 individual social UMAP clusters ± SEM. p-values are extracted for the effect of time on each population’s averaged ΔR^2^ value in linear-mixed effects models. All comparisons within each group are corrected using the Bonferroni method). **k**, ΔR^2^ distributions for a subset of GLM fits during social behaviors in hard fight sessions. Shown distributions for each group are a pool of ΔR^2^ scores from GLM fits across each social UMAP cluster. (Two-sample Kolmogorov-Smirnov tests with all group comparisons FDR-corrected). **l**, Proposed model for updating of local excitatory and inhibitory signals in the SBN. **m**. Proposed model of experience- and observation-dependent change in core aggression network in the SBN. For all comparisons, \**P*<0.05, \*\**P*<0.01, \*\*\**P*<0.001. For detailed statistics, see TableS1.

Finally, to directly quantify changes in the real-time relationship of co-recorded signals, we used a series of network models to map the changing relationship of excitation and inhibition across experience and observation. We tested the prediction that training might weaken the local coupling between excitatory and inhibitory populations in the core aggression network across training during attack. We used a linear model to predict the excitatory attack-aligned neural signal from each site using its local inhibitory signal, and looked at the fits separately for attacks early during training and late during training (Fig. 5c). We found that fits were systematically worse for the attacks occurring later in the training the core hypothalamic and amygdala network, and that this property was shared by the OBS but not the NON group. This decrease in local coupling persisted into the hard fight in the EXP group, with the PMv emerging as a key shared site of local plasticity (Fig. 5e,f). Next, to map the changing influence of the broader network on the activity of a key aggression hub across training, we predicted the excitatory VMHvl signal with signals from the social behavior network and then tested how each fit would be impacted with each signal shuffled one at a time (ΔR^2^, Fig. 5g-k). We quantified this difference for attack-aligned activity (Fig. 5g-h) and for the full signal during social clusters (Fig. 5i-k). We observed that model fits were less impacted following inhibition ablation for late attacks during training and also during the hard fight (Fig. 5h) and with shared plasticity in the OBS individuals emerging in several key aggression-control regions (PMv, MeA, AH, POA).

Overall, these data strongly support a model by which repeated aggression experience can promote widespread decoupling of inhibition from the activity of aggression output control regions across experience training, which generalizes to new aggression contexts. A subset, but not all, of these changes, are mirrored in animals merely observing aggression, suggesting that these shared changes can similarly promote specific behavioral strategies (Fig. 5l-n).

## DISCUSSION

Here, for the first time, we show that both aggression experience and observation recruit a brain-wide network of excitatory, inhibitory, and valence signals, and that changes to this activation pattern predict a shared behavior-control policy in a new aggression context. First, both groups (experience and observation) show a decreased latency to attack in the novel context, a classic signature of an aggression winner-effect and of aggression “instigation” ^23,32,33^. However, beyond changes to attack timing, both groups exhibit a new shared strategy consisting of two interpretable behavioral signatures: First, individuals in experience and observation groups spend more time in aggression preparatory postures and similarly transition between aggression-preparatory clusters, indicating their readiness to fight. Second, both groups spend less time in distant inattentive states where they might be vulnerable to attack. Interestingly, behaviors that involve direct social contact show greater change following direct experience than following observation suggesting that there may be some limits to what can be changed through observation alone.

While aggression “mirror” neurons have been previously been reported in a single hypothalamic population (progesterone receptor-expressing VMHvl neurons, a subset of excitatory VMHvl neurons ^11^), our data suggest that the representation of aggression during observation may be a network-wide phenomenon, activating multiple cell-types across an extended aggression-control network. Moreover, this activated network is largely distinct from cortical and subcortical networks believed to be active in observational fear learning^34^, though we do not exclude its involvement here. The current data is consistent with models of embodied social cognition, in which observation of aggression might activate neural populations associated with aggressive arousal, physiological signaling, and reward monitoring, rather than merely encoding a reflection of action or fear state.

While we observe many similarities between activation patterns of experience and observation individuals, one key difference is the unique activation patterns of signals related to reward (NAc-DA) and punishment learning (LHb) (Fig. 2). While the experience of attack robustly evokes dopamine release in the NAc (putative reward signal) without a concomitant punishment signal in the LHb, we do not observe this reward-biased pattern reflected in observers. Instead, observers exhibited greater trial-to-trial variability, but with signals largely biased towards punishment during attack observation relative to the experience animals. Intriguingly, this suggests that observers may not automatically take the point-of-view of the aggressor, and instead may learn vicariously from both the animal being attacked as well as from the aggressor. This data is consistent with recent evidence demonstrating that observation of a conspecific in pain can actually produce resilient phenotypes and that this is controlled by modulation of LHb activity^35^. Rather than solely supporting a model of appetitive reinforcement during aggression, these data may suggest a broad form of goal emulation^36^, whereby the observer may learn a strategy (avoid being attack-vulnerable) that requires a vicarious reflection of outcomes from both participants.

Across training, we reveal shared changes in network-level plasticity across the training phase in both experienced and observed animals that are not seen in the nonsocial groups. These changes suggest a network-wide “relaxation” of inhibition during critical moments around aggression and these changes persist into the new aggression context. In particular, we observe a shared plasticity motif in the core hypothalamic aggression circuit, whereby inhibition becomes less predictive of neural signals in the VMHvl and PMv. Excitatory signals that exhibit decoupling from the VMHvl (i.e. POA) have been shown to encode aversive social information, suggesting this may exert less influence following repeated exposure^37^. Notably, only one circuit relationship appeared to be strengthened as a result of experience training: the relationship between excitatory populations VMHvl and lPAG, a region known for its role as the attack effector^38^, suggesting direct experience is required to strengthen effective connectivity with motor output structures.

Updating behavior following social exposure likely results from a process that involves both evoked neural activity *and* hormonal cues. Social transmission of nonaggressive social behaviors, including parental and stress-related learning, has been shown to require hormonal modulation^39,40^, and many decades of work have shown that aggression itself robustly activates the hypothalamic-pituitary axis^41^. In addition to hormone-mediated stress responses, previous data across varied vertebrate species including both mammals, birds, and reptiles has shown that direct aggression experience can result in a testosterone surge that peaks tens of minutes after an encounter^42^. Data from both fish and humans have also shown that watching competition can result in a spike in circulating hormones, including testosterone^43,44^, suggesting that this may also be a conserved process. The core aggression network, including both the VMHvl and PMv are rich in receptors for sex hormone receptors, and also project to neurons critical for modulating pathways for hormonal release^45^, situating this network as both a driver and direct receiver of hormonal modulation. Networks expressing esr1 are robustly activated by attack^46^, and activation of these receptors can generate a host of genomic and nongenomic changes that can alter neuronal excitability and synaptic maintenance^47^. Genetic deletion of esr1 in GABAergic but not glutamatergic neurons, particularly in regions with known inhibitory inputs to the VMHvl (MeA, BNST, MPOA) has been shown to be critical for the patterning of male-specific territorial behaivors^48^, suggesting that receptor-expressing inhibitory populations in the core aggression network may be a key site of plasticity. A mechanism for shared hormonal modulation of subcortical circuits may contribute to the stabilization of neural change across long timescales. Future experiments will explore the influence of these shared physiological signals on shaping aggression-control policies.

Classic studies in the psychology of social learning have asserted that aggression learning may occur through a cognitive “disinhibitory” process^49^. Here, for the first time, we find direct circuit-level evidence for decreased influence of inhibition onto core aggression-control circuits that both emerges following repeated experience and observation, and that these changes can generalize to novel and uncertain contexts. While in ethological contexts, this plasticity may have evolved to facilitate adaptive self-defensive strategies, these data may also shed light on the underlying circuit mechanism by which extreme levels of aggression exposure in humans may lead to maladaptive behaviors.

## MATERIALS AND METHODS

### Mice

All animal procedures were approved by the Princeton University Institutional Animal Care and Use Committee and were in accordance with National Institutes of Health standards. A total of 182 experimental animals were used in this study. For multifiber photometry recording studies, we used 23 Vgat^+/−^ ^50^ backcrossed with CD1 (Charles River, Crl:CD1(ICR), 022) males for a minimum of 3 generations, 115 wild-type BALB/cj males (Taconic stock No. 1-99), and 42 wild-type CD1 males. Vgat^+/−^ and wild-type CD1 mice that were used as experimental subjects for freely moving or observational assays were sexually naïve and between the ages of 12-18 weeks. BALB/cj mice that were used as social targets for freely moving assays were sexually naïve and between the ages of 8 and 26 weeks. Aggressor animals used during hard fights were either sexually naïve or retired breeders between the ages of 16-25 weeks. Mice were housed in a 12-h light–dark cycle with experiments taking place exclusively during the dark phase. Food and water were given ad libitum. Ambient temperature was maintained at 21–26 °C and humidity at 30–70%. Prior to initiating fiber photometry experiments, Vgat^+/−^ and wild-type CD1 mice used for aggression and/or observation experiments were socially isolated for at least 1 week. All animals were bred within the Princeton University animal facilities or purchased from Taconic Biosciences or Charles River.

### Surgery

#### Viral strategy

At 12-18 weeks of age, Vgat^+/−^ mice were anesthetized (isoflurane at 3–5% for induction and 1–3% for maintenance) and placed in a stereotaxic frame before proceeding with injections and optic fiber implanting. A 1:1 300nL cocktail of Cre-out GCaMP6f^51^ (Ef1a-Cre out-GCaMP6f) and Cre-activated jRCaMP1b (pAAV.Syn.Flex.NES-jRCaMP1b.WPRE.SV40) was delivered unilaterally into each of the following targeted brain regions: PrL (AP −1.40, ML 0.35, DV −1.70), vLS (AP −2.60, ML 0.60, DV −3.25), mPOA (AP −3, ML −0.40, DV −4.75), BNST (AP −3.30, ML 0.85, DV −3.75), AH (AP −3.7, ML −0.50, DV −4.80), VMHvl (AP −4.65, ML 0.72, DV −5.45), LHb (AP −4.70, ML −0.33, DV −2.65), MeA (AP −4.70, ML −2.25, DV −4.90), PA (AP −5.50, ML −2.40, DV −4.95), PMv (AP −5.50, ML −0.58, DV −5.55), lPAG (AP −7.80, ML 0.50, DV −1.65). An additional infusion of 150nL of GRAB DA1m (hSyn-GRAB_DA1m-WPRE-hGHpA) was delivered to the NAc (AP −1.80, ML 1.15, DV −3.70). Surgery coordinates for each area were calculated relative to the skull surface in the posterior edge of the rostral rhinal vein. All regions included in the final analyses had correct viral expression and fiber tip positioning based on histology.

#### Multi-site fiber strategy

Following viral infusions, a custom-built multi-fiber implant containing twelve 200-μm-core optic fibers (220 μm cladding, 0.37 NA, THORLabs) is fixed to the skull with Metabond (Parkell). This implant is lowered into the brain to a depth of 5.55mm relative to the brain surface of the VMHvl using a custom stereotaxic attachment designed to hold multi-fiber implants. Multi-fiber implants are 3D-printed (ABS-Like Micro Resolution, Protolabs) and designed with fiber through-holes in AP and ML locations matching those used for stereotactic viral infusions. Prior to fixing the fibers in the implant, these are first cleaved with a manual fiber cleaver (THORlabs) and subsequently cut with a ruby scribe to match the specific length of each area’s depth in the brain, with added length accounting for the widths of the implant and each area’s craniotomy. Supplementary table 2 includes all the coordinates and measurements in millimeters used for viral deliveries and implant construction. Note that, prior to implantation, bulk light efficiency for every implant was measured, with implants with an average efficiency below 70% getting discarded. Following surgery completion, animals were allowed to recover for at least 3 weeks prior to the start of recording experiments.

### Multi-site fiber photometry

#### Recording

To record from up to twelve sites simultaneously, we used a custom 12-fiber bundled, branching patchcord (200-μm fiber, 220 μm cladding, 0.37 NA, Doric). One end of the patchcord terminated in an SMA connector in a honeycomb pattern, with the other end affixing all branching fibers in a custom-3D printed connector at coordinates matching the fiber placements in the multi-fiber implant. The implant fibers were aligned to the patchcord fibers in the connector using four 1 mm steel dowel pins (McMaster), and the two components affixed via 3-4 screws (Open Ephys ShuttleDrive). Recordings were made via a custom written data acquisition program in DAQAmi (Measurement Computing) and using the Neurophotometrics FP3002 system in combination with Bonsai. LEDs delivering three excitation wavelengths (470nm for optical excitation of GCaMP6f and DA1m GRAB, 560nm for optical excitation of JRCaMP1b, and 415nm for a calcium- and dopamine-independent control, light intensity ∼ 70 μW for each fiber tip of the implant) were interleaved at 40 Hz (for a total 120 Hz sampling rate) throughout recordings. Fluorescence emission was split at 552 nm into green and red channels and focused onto a CMOS sensor for detection. Photometry signals in arbitrary units were collected from twelve sites drawn separately in the green (GCaMP) and red (RCaMP) channels across the imaged honeycomb pattern of fibers.

#### Signal processing

Mean fluorescence in arbitrary units was collected as raw data for every session. To normalize this signal and correct for potential motion artifacts, we performed processing on individual photometry signals as follows: 1) baseline subtraction using an adaptive iteratively reweighted Penalized Least Squares algorithm^52^, 2) z-score standardization, and 3) signal subtraction using motion-artifact control signals from 415 nm excitation. For step 3, specifically, we fitted a multilinear regression (sklearn.linear_model.Ridge) for each individual photometry signal using as predictors all calcium-insensitive control signals from 415nm optical excitation taken from the same recorded session. The regularization strength (alpha) for each fitted model was determined individually for each session via grid search. The predicted control signal was then used to subtract the real photometry signal. The resulting corrected signals from this process were finally normalized by their mean value during a 1-minute baseline period prior to any social interaction. All analyses were completed in the resulting z-ΔF/F data.

#### Behavioral assays

Animals undergoing fiber photometry recordings were subjected to longitudinal resident-intruder experiments under varying conditions. For 8 consecutive days, animals undergoing longitudinal aggression experience (EXP) received unconstrained access to three different male BALB/cj mice across three separate 9-minute sessions, each preceded by a 1-minute baseline without social stimuli. These interactions took place in the homecage of the experimental animals (Optimice Auto-Water Cage Bottom with SE Lab Group cage, 34.3cm L x 29.2 cm W x 15.5 cm H). For longitudinal observation (OBS), a cage extender made of black acrylic and clear plexiglass was built to match the dimensions of the homecage, along with a compartment in the front of the cage extender designed to hold each observer mouse (10 cm L x 13 cm W x 17 cm H). The observer platformed compartment was separated from the cage floor via a clear plexiglass barrier with multiple horizontal cutouts allowing for visual and olfactory access of the cage bottom. Every OBS mouse was longitudinally paired with a unique CD1 male mouse, with the latter undergoing 8 consecutive days of three back-to-back resident intruder sessions in its own homecage. For each day, each observer mouse was placed in the observer compartment, and the cage extender housing the compartment was placed atop the paired animal’s homecage. Before initiating resident-intruder sessions, OBS mice and the partnered CD1 mice were allowed up to 15 minutes of habituation while separated by the plexiglass barrier. Afterwards, each resident-intruder session consisted of 9 minutes of social interaction between the paired CD1 mouse and a different male BALB/cj mouse, each preceded by a 1-minute pre-intruder baseline. Following observation, two animals from the OBS group were subsequently trained in the EXP paradigm. Behavior from these animals was excluded from behavior analysis but neural data was conserved for neural analysis. Finally, animals undergoing nonsocial experience (NON) were longitudinally exposed to an inanimate toy resembling a mouse over the same timespan as EXP and OBS mice, and across the same number of sessions per day. Note that, while throughout the training period OBS individuals did not engage in social behavior, photometry recordings were still performed in these individuals in order to mirror any bleaching-related signal decays present in EXP and OBS individuals following longitudinal imaging. On day 9, each animal from each group was subjected to three back-to-back 4-minute resident-intruder sessions in their homecage, preceded by a 1-minute solitary baseline. In each session, each animal received unconstrained social access to a unique CD1 male screened a priori for aggression against other CD1 males (“hard fight”). CD1 aggressors were reused for more than a single hard fight session provided they were never defeated in prior sessions.

#### Behavior acquisition

Social interactions were recorded using top and side cameras (40 Hz, FLIR, Spinview). These were controlled with a data acquisition device (DAQ) and associated software (Measurement Computing Corp), which further coordinated the FP3002 CMOS camera in synchrony. The top camera was centered 50 cm above the homecage of experimental mice for each recording session, while the side camera was placed at a height of 30 cm and within 3 cm of the observer compartment. Black paper-chip bedding was used for every recording and miscellaneous items such as food pellets were cleared from homecages prior to recording.

### Behavioral analysis

#### Behavioral annotation

For supervised classification of behaviors during aggression, ground truth was determined by manual annotations of videos scored with BORIS (ref). Only attack and fighting behaviors were annotated, as defined by rapid forward movements from the resident, such as chases or lunges, directed at the intruder mouse. The number of annotated frames included 29,805 attack frames out of 759,074 across 49 videos. Annotations from the hard fights were carried out blind to the group identity of the behaving mice.

#### Pose tracking

SLEAP software (v.1.2.2) was used to predict resident and intruder positions across social interactions. To do this, 6305 frames from 141 social interaction videos were labeled for tracking of resident and intruder body positions. These 6305 frames were divided into training and test sets with an 80:20 split. The following points for both resident and intruder animals were labeled and subsequently tracked for all social interaction videos: nose, head, neck, left ear, right ear, trunk, left hindpaw, right hindpaw, left forepaw, right forepaw, tailbase, bottom segment of tail, middle segment of tail, upper segment of tail, and tail tip. Training of the SLEAP model was performed over 93 iterations using a bottom-up approach with custom parameters (initialized with Unet and batch-size of 4).

#### Feature definition

To define the posture of resident and intruder mice, frame-to-frame pose data were converted to the following behavioral features:

1. Intruder body length: Euclidean distance between the head of the intruder and its tailbase.
2. Resident body length: Same as above but for the resident
3. Intruder body orientation: Angle spanning the centroid, head and nose of the intruder
4. Resident body orientation: Same as above for the resident
5. Intersection over union of resident and intruder (IoU): Overlap between resident and intruder bounding boxes by dividing the area of their intersection by the area of their union.
6. Intruder head speed: Instantaneous distance in intruder head position (every 4 frames or 100ms)
7. Resident head speed: Same as above but for the resident
8. Intruder head speed lagged at ± 3s: Sum across lags of intruder head speed, computed within a ± 3s lag window.
9. Resident head speed lagged at ± 3s: Same as above but for the resident
10. Intruder centroid speed: Instantaneous distance in intruder centroid position (every 4 frames or 100ms)
11. Resident centroid speed: Same as above but for the resident
12. Intruder centroid speed lagged at ± 3s: Sum across lags of intruder centroid speed, computed within a ± 3s lag window.
13. Resident centroid speed lagged at ± 3s: Same as above but for the resident
14. Intruder to resident head-centroid angle: Angle spanning the intruder’s neck, intruder’s head and resident’s centroid.
15. Resident to intruder head-centroid angle: Angle spanning the resident’s neck, resident’s head and intruder’s centroid
16. Intruder to resident head-to-tailbase distance: Euclidean distance between the head of the intruder and the tailbase of the resident
17. Resident to intruder head-to-tailbase distance: Euclidean distance between the head of the resident and the tailbase of the intruder
18. Between centroid distance: Euclidean distance between the centroids of the resident and intruder
19. Between head distance: Euclidean distance between the heads of the resident and intruder

Distance-related features were converted from pixels to centimeters using custom MATLAB software based on our behavior camera’s stereo parameters and pixel depth information.

#### Feature preprocessing

To standardize feature processing across all sessions, the mean and standard deviation of every feature for every recorded session were pooled, and subsequently averaged. Features for each session were processed as follows: 1) removal of outliers 3 standard deviations away from the session mean, 2) smoothing via Gaussian filter of 0.275 s, 3) linear interpolation of missing values, and 4) z-scoring using the mean and standard deviation pooled from collating features across sessions. The collated mean and standard deviation values for every feature were used for z-scoring in individual sessions to capture a wider variance in feature dynamics as opposed to unit variance which may reflect data fluctuations not scalable across samples.

#### Supervised behavior classification

To identify attack behaviors across our entire video dataset, we trained a supervised multilayer perceptron (MLP) on SLEAP-derived feature data using the Keras library (Keras and Tensorflow versions 2.11.0). A sequential model was constructed, consisting of 1) an input layer shaped after the behavior features, 2) 2 hidden layers interleaved with batch normalization layers to normalize the activations of the previous layer, and 3) an output layer with a sigmoid activation function for binary classification. The input and middle hidden layers consisted of 512 units with “ReLu” activation functions while the output layer consisted of a single unit. The target matrix was a binary indicator for hand-annotated attack labels, which were pooled from 49 videos, totaling 29,805 attack frames versus 729,269 non-attack frames. The feature matrix included the 19 features described above for each video frame, with 19 additional features adding lagged components of the existing feature set, including the instantaneous distance between resident and intruder centroids. Because non-attack frames far outnumbered attack frames, we oversampled the number of attack frames via a synthetic oversampling technique for time series^53^. This involved generating synthetic samples for both attack labels and accompanying feature matrices by randomly sampling both with replacement and then combining these synthetic samples with the original data to balance each distribution while maintaining the underlying temporal structure. The resulting data distribution totaled 699,464 attack frames versus 729,269 non-attack frames. This distribution was divided into training and test sets with a 67:33 split. Upon training the MLP, performance was evaluated via the generation of a confusion matrix from the model’s predictions on the held-out feature matrix. The following statistics summarize the model’s performance: true positives 1.0, false positives 0.00047, true negatives 1.0, false negatives 0.003. Finally, probability distributions for attacks from MLP predictions were generated for every social interaction video. The probability threshold for labeling an attack versus non-attack in these distributions was set to 0.50, with attack labels further being filtered to be at least 200ms long in consecutive frames and separated from segment to segment by at least the same amount of time. Note that supervised classification of NON behavior during the training period was not performed since individuals in this group were not socially interacting; instead, labels for individuals in this group were generated via a yoked control design whereby labels from a randomly assigned individual in the OBS or EXP groups were paired to each NON individual for each recorded day and session.

#### Unsupervised behavior embedding and clustering

To comprehensively characterize social behavior in an unbiased manner, we adapted previous unsupervised behavior classification strategies involving the reduction of high-dimensional behavior features to a low-dimensional embedding and the definition of behaviors as high-density clusters within this embedding. To achieve dense clusters, we embedded the 19 behavior features described above into a 2-dimensional embedding using UMAP. To generate this embedding, an importance sampling approach was taken. This approach increased the likelihood of generating a final version of the embedding featuring rare or nuanced behaviors. Importance sampling altogether involved two rounds of UMAP reduction. In the first round, 80,000 video frames were uniformly sampled from 504 social interaction videos. As aggression can often happen briefly and thus be underrepresented in the total pool of feature frames used for reduction, an additional 20,000 feature frames aligned to aggression labels according to our supervised attack classifier were randomly added to the frame pool. This total pool of 100,000 feature frames was next embedded into a two-component UMAP manifold (umap from umap-learn, version 0.5.5, n_neighbors=50, min_dist=0.1), which was further binned into a 200 × 200 histogram, smoothed with a 2D gaussian kernel of 3.5 standard deviations and parcellated into 21 clusters with watershed (skimage.segmentation.watershed). As UMAP is non-parametric, a separate model was implemented to learn the mapping from the original 19-dimensional feature space to the new 2D UMAP space. An MLP (sklearn.neural_network.MLPRegressor, hidden layer size of 1000 × 500 × 100 × 50 units) was used for this purpose, allowing the data from every video frame to be represented in UMAP space and thus fall into any of the 21 clusters previously defined.

To generate the final UMAP embedding, over 20,000 frames were sampled from the 21 clusters generated previously. Specifically, 2 random frames from each cluster and for each recorded session were pooled together for a total of 21,168 frames (2 frames x 21 clusters x 504 videos = 21,168). From these sampled frames, the 19-dimensional feature set was embedded into 2D UMAP space. Next, the full set of video frames were mapped into this final UMAP manifold using another MLP. The resulting embedding was turned into a 2D histogram, smoothed with a Gaussian kernel of 2 standard deviations and divided into 13 final clusters with watershed. All behavior analyses involving behavior clusters are performed using this final UMAP segmentation. Note that unsupervised classification of NON behavior during the training period was not performed since individuals in this group were not socially interacting; instead, cluster labels for individuals in this group were generated via a yoked control design whereby labels from a randomly assigned individual in the OBS or EXP groups were paired to each NON individual for each recorded day and session.

#### Behavior transitions and relative similarity of unsupervised behavior profiles

The expression of UMAP clusters was quantified via the occupancy (fraction of time in cluster relative to whole recording session) or persistence (mean number of consecutive frames comprising a segment of a behavior cluster) of each cluster, in addition to each cluster’s probability of transitioning to every other cluster on a frame-to-frame basis.

To quantify the transition structure of behavior clusters, pairwise transition probabilities were calculated as the average number of times the first cluster transitions into the second cluster. For each session, a 13×13 matrix containing pairwise transition probabilities for all 13 clusters was constructed. A global transition matrix was then generated by averaging the transition matrices of each individual from each group from both training period and hard fight sessions. A permutation test was conducted for each pairwise comparison in this global 13×13 matrix, excluding diagonal elements, to determine which transition probabilities were above chance. For each comparison, the observed value was extracted, and a permuted mean was generated by randomly sampling from the flattened matrix 10,000 times. The p-value for each comparison was computed as the proportion of permuted means greater than or equal to the observed mean. Furthermore, the global transition matrix was sorted via unsupervised hierarchical clustering (scipy.cluster.hierarchy.linkage) using the complete linkage method and Jensen-Shannon distance metric, with optimal leaf ordering to enhance the interpretability of the dendrogram. The resulting dendrogram was visualized with a color threshold of 0.8 to distinguish clusters.

To assess whether behavior occupancy, persistence or transition probabilities were predictive of group identity in the hard fight, a leave-one-session-out cross-validated approach (sklearn.model_selection.LeaveOneOut) using support vector machine classifiers (sklearn.svm.SVC) was employed. For each session, three kinds of vectors were generated: 1) a 13×1 vector wherein each element represented the average occupancy each cluster, 2) a 13×1 vector wherein each element represented the average persistence of each cluster, and 3) an 81×1 vector containing the flattened pairwise transition probabilities for 9 UMAP clusters identified as social behaviors. For each vector type above, a separate leave-one-out cross-validation procedure was implemented. Here, a new classifier was iteratively constructed in each fold consisting of 68 vectors of occupancy, persistence or transition probabilities as the training set and a single held-out vector for the classifier to predict on. The F1 score of all held-out predictions was subsequently saved. Finally, to assess statistical significance of this leave-one-out cross-validation strategy, a null distribution was created by shuffling group identity labels 1000 times and repeating the cross-validation process, recording the F1 scores for each shuffle. The p-value for each model’s performance was computed as the proportion of shuffled F1 scores greater or equal to the test held-out F1 score.

To calculate behavioral similarity between individuals in each group during hard fight sessions, the above vectors containing averaged occupancy, persistence and transition probability variables were used. For every individual in the OBS group, similarity in occupancy or behavior transitions to every individual in the EXP or NON groups was computed via the Jensen-Shannon (JS) divergence of the occupancy or transition probability vectors of each pair of individuals. JS divergence here is employed as follows:

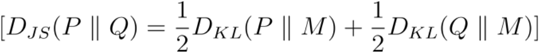

where P represents the probability distribution corresponding to either behavior occupancies or pairwise transition probabilities of an OBS individual, Q represents the same variable for an individual from either the EXP or NON groups, M is the average distribution of P and Q, and D_KL_ is the Kullback-Leibler divergence. The Kullback-Leibler divergence is defined as:

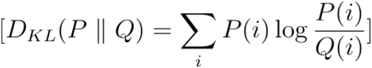

Note that JS divergence is 0 when comparing equal distributions and larger values indicate larger divergence. Therefore, as presented in Fig. 1x, values closer to 0 reflect greater relative similarity between animals. Furthermore, because behavior persistence distributions are not probabilistic, Euclidean distance was used instead as a measure of similarity between individuals in each group.

#### Feature encoding

To characterize how the behavioral features were encoded in the neural activities across the network, we fit independently for each feature and each neural signal a Linear-Gaussian Generalized Linear Model (GLM) with a smoothing regularizer that is chosen based on cross-validation. The input to the model is a one-hot vector representation of the feature value, which was binned in 20 total bins. The fitted GLM weights reflect the tuning curve for that specific region and feature, with positive weights suggesting an increase in activity and negative weights a decrease. To test group differences in the tuning curves, we leave one animal out at a time and fit group-specific tuning curves for all the other animals. Then, we compute the r squared prediction of the neural activity for the missing animal based on the tuning curves fit for its own group, as well as curves from the other groups. Lastly, we do a paired t-test pairwise between any two groups, with samples consisting of the r squared values of a particular group computed with its own tuning curves versus the r squared values of the same group computed with the tuning curve of a different group.

### Neural analysis

#### Time-warping of attack signals

To identify neural activity patterns occurring reliably on a trial-by-trial basis during attacks, a time warping approach was employed^29^. For animals with correct histology for all brain areas in the EXP group, neural activity for each population aligned to each attack (−6s to +6s) across the training period was pooled into a 3d tensor (attack x attack window size x population) and a shift-only time warping model (affinewarp.ShiftWarping) was fit to these data. Shift-only time warping can model translations in time across signals in tensors on a trial-by-trial basis without stretching or compressing said signals, thus conserving temporal structure while addressing alignment issues that could arise from jitter in behavior labeling. A maximal allowable shift of 1.2s with penalties on the magnitude of the shifts and warping templates (values of 0.1 and 1 respectively) were allowed in order to further preserve the temporal structure of neural signals. Following model fitting, a new tensor with shift-warped signals was generated from the non-warped tensor. The same shift-only time warping model was lastly applied on held-out individual tensors of animals with missing signals due to incorrect fiber placement. For animals in the OBS and NON groups, separate shift-only time warping models were applied but with maximal allowable shifts of 2.4s and with the same penalties on the magnitude of the shifts and warping templates as above.

For neural signals recorded during hard fight attacks, a 3d tensor with the same structure as above containing data from all three animal groups was generated and fit with a shift-only time warping model with a maximal allowable shift of 2.4s. A new time-warped tensor was generated from this model. This hard fight time-warped model was further used to generate time-warped tensors from other animals in the dataset with missing signals due to incorrect fiber placement.

#### Relative similarity of neural activity profiles

To compare similarity in neural signals across the behavior space, each signal from each session was binned into a 13×1 vector, wherein each element in this vector represented the mean z-scored ΔF/F of the signal for each UMAP cluster. Next, the mean activity vector of each session for every individual was compared to the mean activity vector of each session from every other individual using cosine similarity. Cosine similarity was applied as follows:

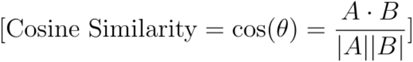

Where A and B represented mean activity vectors for different individuals. Because cosine similarity measures the orientation between activity vectors rather than magnitude differences, it is useful for revealing which populations have overlapping representations of behavior regardless of scale. Note that cosine similarity is 0 when comparing vectors with the same orientation and thus larger values indicate higher divergence. Importantly, for similarities computed during the training period, 10×1 vectors corresponding to the total of 10 social UMAP clusters (1-10) were used. This is due to the presence of observation-evoked activity during nonsocial clusters which may be driven by increased proximity between the observer and one of the socially interacting mice on the cage floor (Fig 3x, Extended Fig 4x). To avoid confounds related to behaviors in which the observer may investigate a conspecific directly during clusters 10-13, these clusters were excluded from the mean activity vectors being compared. Further, because this confound was not present in hard fight sessions since all recorded subjects were involved in social interaction, all 13 UMAP clusters were represented in the mean activity vectors derived from these sessions. Similarities for each pair of individuals across all sessions were calculated, with the averaged cosine similarity between each OBS individual and each EXP or NON individual being quantified for both the training period and hard fights separately.

#### Decoding of aggression

To map the relationship between attacks and neural activity, a frame-by-frame decoding approach was employed using *k*-nearest neighbors classifiers (sklearn.neighbors.KNeighborsClassifier). Prior to classification, predictor matrices containing multi-dimensional SBN time series for each of the three social interaction sessions within each day were concatenated in time, as were the accompanying binary attack labels generated from supervised behavior classification for the same day. Because non-attack frames outnumbered attack frames for each day, SBN matrices and labels were under sampled to accommodate the minority class (imblearn.under_sampling.RandomUnderSampler). Next, separate classifiers were trained for each day and animal on the corresponding SBN signal matrix to predict the corresponding binary attack labels. For each classifier, cross-validation via grid search (sklearn.model_selection.GridSearchCV) of the number of neighbors, weights and metric was performed in a stratified 3-fold design to preserve the percentage of samples for each class. The accuracy score, i.e. the fraction of frames accurately classified, from the highest scoring classifier obtained from this cross-validation approach was recorded for each day. Moreover, to test whether SBN activity from earlier time periods was predictive of behavior in later periods, classifiers trained from each day of training were subsequently tested on two additional held-out SBN matrices: 1) held-out SBN signals from day 8 of the training period, and 2) held-out SBN signals from hard fight sessions. Null accuracy distributions from predictions made on within-day training period or hard fight SBN matrices, in addition to day 8 or hard fight sessions from training period SBN matrices, were generated by shuffling all of the above SBN matrices and subsequently re-predicting on the newly shuffled matrices. In summary, classifiers that were generated from cross-validated grid search on data from each day of the training period were tested on held-out data from same day (train), held-out data from day 8 (d_n_->d_8_), and held-out data from the hard fight (d_n_->HF), with a final class of classifier generated from a subset of hard fight signals and tested on held-out data from the same sessions (HF).

To determine the importance of excitatory versus inhibitory circuits in attack decoding, a feature importance approach was implemented. Here, the same decoding strategy as above was used with two separate classes of SBN matrices: ones containing only excitatory signals, and ones containing only inhibitory signals. Excitatory-only or inhibitory-only classifiers were thus trained to predict binary attack labels generated from supervised behavior classification. The resulting accuracies of these excitatory-only or inhibitory-only classifiers were used to subtract the corresponding accuracies of the full classifiers from predictions generated on held-out data from the same day, from day 8, and from hard fights (both using signals from the training period and from the hard fights). The difference in performance between excitatory-only or inhibitory-only classifiers relative to the full classifier was quantified as a percentage loss in accuracy, with any potential improvements in the performance being set to 0 loss.

### Regression-based strategies for quantifying multi-site interactions

To map the statistical relationship between signals in the SBN, multiple regression-based approaches were taken.

To assess coactivity between local excitatory and inhibitory signals, separate ordinary least-squares regressions (statsmodels.regression.linear_model.OLS) were used to model excitatory activity aligned to attacks using only local inhibitory signals from the same period (6s to +6s window aligned to attack start). The regression score (R2) resulting from each fit was used as a measure of how well local excitatory and inhibitory signals from each SBN region were correlated with each other. To map whether correlations between both signals changed over time, for each animal, R2 distributions from each OLS fit during the first 30 versus last 30 overall attacks of the training period were compared. Furthermore, to map whether any correlations carried over from training to the hard fight, the same OLS strategy was employed for attacks in the hard fight.

To model connectivity across brain areas, Generalized Linear Models (GLMs) were employed in combination with a feature importance testing approach. For every attack or social UMAP cluster, a GLM (statsmodels.genmod.generalized_linear_model.GL M, family=‘Gaussian’) was trained on a feature matrix containing signals from PMv E, AH E, POA E, lPAG E, MeA E, PA E, VMHvl I, PMv I, AH I, POA I, MeA I and PA I, altogether, to predict VMHvl E activity, a population causal to aggression^9^. Following training, each predictor in the feature matrix was iteratively shuffled and the original GLM was used to predict on this newly shuffled matrix. The accuracy score (R^2^_shuf_) from this prediction was used to subtract the original model’s accuracy score (R^2^_full_), with the resulting ΔR^2^ being used as a measure of the shuffled predictor’s importance in encoding VMHvl E activity. For GLM predictions during attacks, GLMs were fit to feature matrices aligned to the first 30 versus last 30 attacks (−6s to 6s around attack start) of the training period to assess time-related changes in feature importance, in addition to all attacks from all hard fight sessions. For GLM predictions during broader social behaviors, GLMs were fit to feature matrices aligned to individual UMAP clusters 1-10 for each day of the training period and the hard fights. To determine changes in feature importance over time, ΔR^2^ scores from fits aligned to each cluster were averaged for each day of the training period and the effect of time, i.e. each day, was quantified via a linear mixed effects model. To determine feature importance during the hard fight, ΔR^2^ scores from each social cluster for each individual were pooled into separate group distributions and subsequently compared. Only individuals with correct fiber placement for all of the areas used to model VMHvl activity were used for this analysis.

#### Whole-brain histology for photometry fiber recovery and viral expression assessment

Following the completion of our photometry and behavioral studies, animals were deeply anaesthetized with an injected ketamine and xylazine cocktail, and perfused with 4% paraformaldehyde (PFA) dissolved in 1× PBS. Brains were extracted, post-fixed in 4% PFA for 12– 24 h. For a subset of animals (5/12 EXP), following PFA post-fixing, brains were cleared using a previously described iDISCO+ protocol^54^. Cleared brains were then imaged with light sheet microscopy at ×1.1 or ×1.3 magnification, and these images were registered to a common reference atlas as previously described^55^. We chose to register to the Paxinos Atlas^56^ to map fiber tip placement to anatomically defined sub-areas of each SBN location, such as the lateral division of the periaqueductal gray or the NAc core and shell structures. Fiber tip locations were manually identified in registered light sheet images and plotted together in the common Paxinos coordinate frame. For another subset of animals (7/12 EXP, 7/7 OBS and 6/6 NON), following PFA post-fixing, extracted brains were cryoprotected in 30% sucrose and embedded in optimal cutting temperature mounting medium (Fisher Healthcare Tissue-Plus OCT compound, Fisher Scientific) for freezing over dry ice. Cryosections of the frozen tissue (30 μm slices) were made and stamped directly onto glass microscope slides. Slices were washed with PBS, dried, then coverslipped with mounting medium (EMS Immuno Mount DAPI and DABSCO, Electron Microscopy Sciences, 17989-98, lot 180418) and sealed with clear nail polish around the slide edges. After at least 12 h of drying, slides were imaged with a digital robotic slide scanner (NanoZoomer S60, C13210-01, Hamamatsu).

#### Statistical analysis

Sample sizes were not predetermined ahead of time. The following statistical tests were run in Python (v3.9.12): statsmodels (v0.13.2) and scipy (v1.7.3). For determining the difference between two or more distributions, pairwise two-sample Kolmogorov-Smirnov tests (scipy.stats k2_2samp) were conducted. For comparisons between related distributions across multiple samples (e.g. early versus late distributions across multiple brain areas), all comparisons were corrected within-group using the FDR method (statsmodels.stats.multitest.multipletests, ‘fdr_tsbh’ method). For comparisons between independent distributions, corrections were performed within-sample on the group comparisons using the FDR method.For assessing the difference in means in two groups (e.g. mean z-scored ΔF/F differences between EXP and OBS groups), normality for each group was first determined via the Sharpiro-Wilks test (scipy.stats.shapiro), followed by a *t*-test (scipy.stats.ttest_ind) in normally-distributed samples or a Mann-Whitney U test (scipy.stats.mannwhitneyu) for non-normal samples, with multiple comparisons corrections as needed using the Bonferroni or FDR method (statsmodels.stats.multitest.multipletests, methods ‘bonferroni’ or ‘fdr_tsbh’ respectively). For assessing the difference in means between two related samples (e.g. feature importance tests with excitatory versus inhibitory importance), a *t*-test was performed (scipy.stats.ttest_rel). For assessing the difference in means across more than two groups (e.g. occupancy in EXP, OBS and NON groups), normality for each group was first determined via the Shapiro-Wilks test. One-way ANOVA tests (statsmodels.stats.anova_lm) were used for normally distributed data, followed by post hoc *t*-tests for pairwise comparisons which were further corrected via the Bonferroni or FDR methods as needed. For non-normal data, Kruskal-Wallis tests (scipy.stats.kruskal) were performed, followed by post hoc Mann-Whitney U tests for pairwise comparisons which were further corrected via the Bonferroni or FDR methods as needed. For assessing time effects within a group (e.g. ΔR^2^ across days in EXP or OBS groups), a mixed linear model (statsmodels.formula.api.mixedlm, fixed effect of day and random effect of mouse identity) was used. For groups with multiple time effects, multiple comparison corrections were performed using the Bonferroni method. For model and transition probability validation, a permutation test approach was employed. For validation of group identity decoders in Fig1x and Extended Fig1x, F1 score calculation was conducted 1000 times on shuffled data, with the final p-value reflecting the proportion of shuffled F1 scores greater or equal to the test F1 score. For validation of pairwise probability transitions in Fig 1k, a permuted mean was generated from randomly sampling from the flattened transition probability matrix 10,000 times, with the final p-value reflecting the proportion of permuted means greater than or equal to the observed mean. All pairwise transition probability comparisons were corrected using the Bonferroni method.

Our manuscript follows recommendations for inclusion and ethics in global research.

## Data Availability

All behavior and neural data will be made publicly available on and source data for all figures will be available on FigShare following publication of this manuscript.

## Code Availability

All code is available on GitHub repository: https://github.com/FalknerLab/AggressionObservation.

## Acknowledgments

We thank I. Witten, A. Leifer K. Rajan, T. Engel and members of the Falkner laboratories for useful discussions; D. Tsin for feedback on the manuscript, S. Oline for assistance with camera setup and syncing, P. Cherepanova for help with fiber implants, L. Sirrs and A. Le for help genotyping, staff at the PNI Viral Core Facility for reagents and the PNI Brain Registration and Histology Core Facility for assistance with histology. Funding was from DP2MH126375 (to A.L.F.), NIH R01MH126035 (to A.L.F.), NYSCF (to A.L.F.), SCGB (to A.L.F.), Klingenstein Foundation (to A.L.F.) and an Alfred P. Sloan Fellowship (to A.L.F.). A.L.F. is a New York Stem Cell Foundation Robertson Investigators.

---

**Extended Figure 1.**
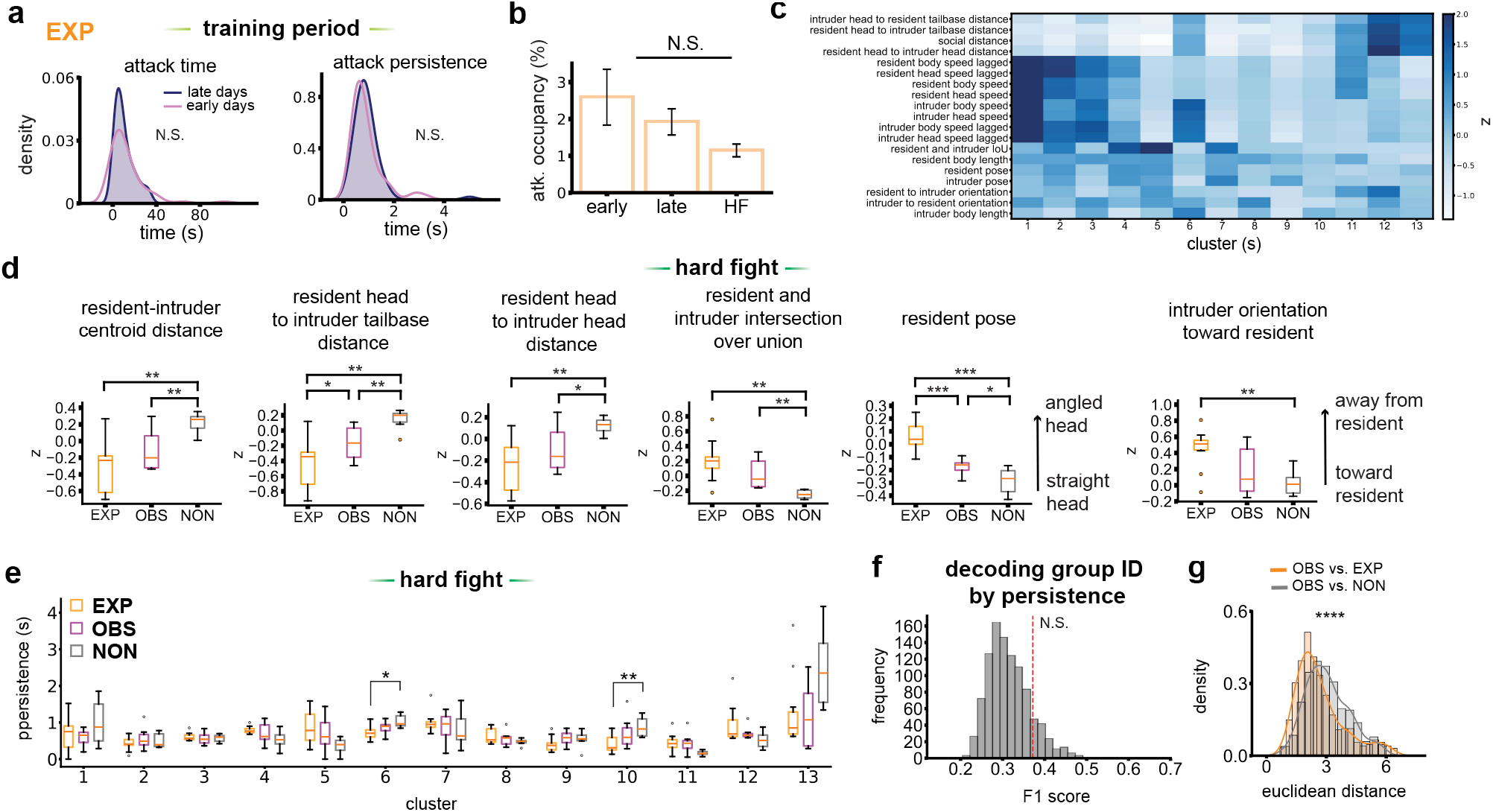
Aggression experience and observation promote increased interaction during hard fights. **a**, Distributions representing time spent attacking (s) (left) and attack persistence, i.e. average length of attack bouts (right) during early (first 2, pink) versus late (last 2, navy) days of the training period in the EXP group. (two-sample Kolmogorov Smirnov tests). **b**, Occupancy (% time spent) of attacks in early (first 2), late (last 2) and hard fight. (Data are reported as mean ± SEM. One-way ANOVA). **c**, Unsupervised UMAP clusters sorted by mean z-value of each SLEAP-derived feature utilized for clustering. **d**, Subset of averaged SLEAP-derived features for each group of animals. The features shown are 1) centroid distance between resident and intruder, 2) distance between resident head and intruder tail-base, 3) intersection over union of the resident and intruder bodies, 4) angle between resident centroid, neck and head, and 5) angle between intruder neck, head and resident centroid. (Data are reported as mean ± SEM. One-way ANOVA tests followed by post-hoc independent samples *t*-tests and FDR adjustments). **e**, Persistence (average segment length in seconds) of all behavior clusters across groups during the hard fight. Box plot lines represent the median occupancy, with the box extending from the first to the third quartiles of the data, and the whiskers 1.5x the inter-quartile range (Group comparisons done via a one-way ANOVA or Kruskal-Wallis tests pending normality, post-hoc independent samples *t*-tests or Mann-Whitney U tests, and multiple comparison corrections using the FDR method). **f**, Leave-one-session-out cross-validation F1 accuracy (0.37) of group ID (EXP, OBS, NON) predictions using occupancy of UMAP clusters. (Test F1 score is evaluated against a null distribution of F1 scores generated by randomly shuffling group labels for each session). **g**, Distribution of Euclidean distance values computed between the occupancies of UMAP clusters of every OBS individual against every EXP (orange) or NON (gray) individual (two-sample Kolmogorov-Smirnov test). For all comparisons, \**P*<0.05, \*\**P*<0.01, \*\*\**P*<0.001. For detailed statistics, see TableS1.

**Extended Figure 2.**
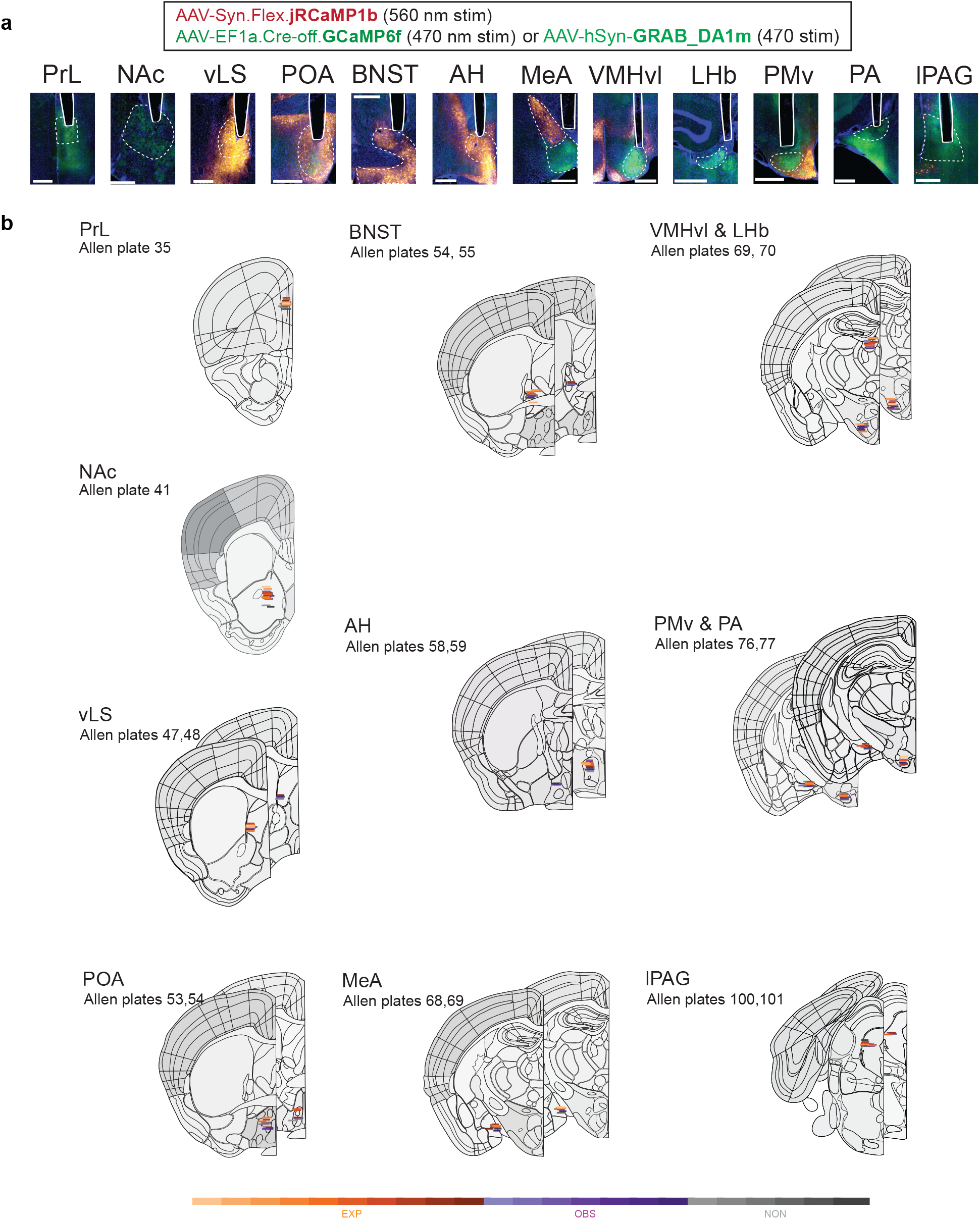
Summary of optical fiber locations and viral expression across all recorded animals. **a**, Representative GCaMP6f (green) or JRCaMP2b (red) expression across all 12 recorded brain areas. Scale bars: 500 μm. **b**, Accepted fiber placements for every experimental subject across all 12 recorded brain areas. Coronal schematics are adapted from the Allen Brain Atlas.

**Extended Figure 3.**
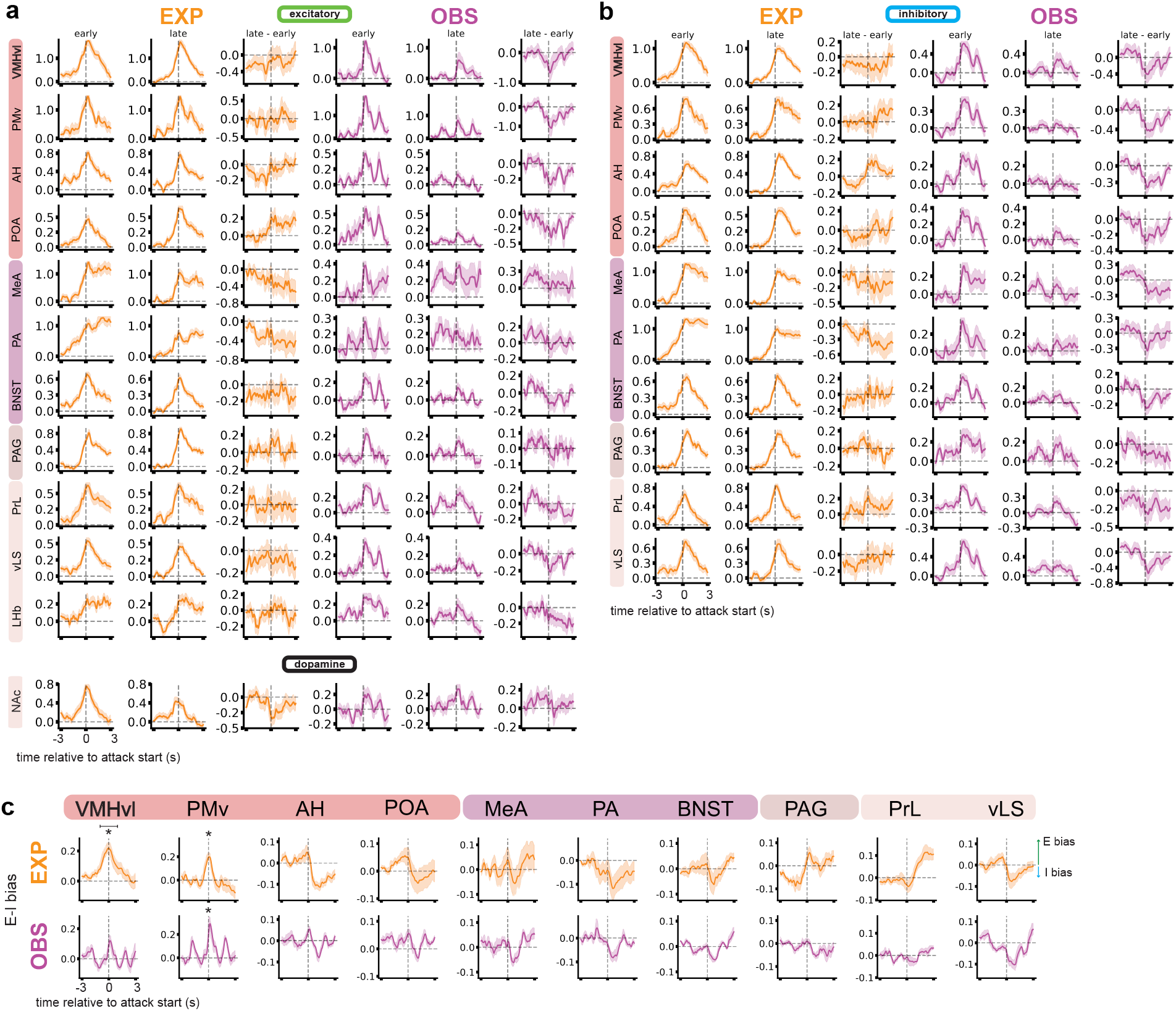
Peri-event time histograms of SBN activity during early versus late attacks. **a**, Group peri-event time histograms (PETH) of all 12 recorded brain areas during training period. *Left:* PETHs of excitatory activity for individuals in the EXP group; PETHs on the left represent first 30 attacks, PETHs in the middle represent last 30 attacks, and traces on the right represent the average difference in late versus early PETHs. *Right:* Same as left but for OBS individuals. **b**, Same as (**a**) but PETHs represent inhibitory activity instead. ç PETHs of excitatory-inhibitory (E-I) bias for all trials of the training period. E-I bias PETHs are computed as the difference between excitatory and inhibitory traces for each attack trial, normalized by the sum of the two for the same trials. (Significance in the time window of −1s to 1s is evaluated relative to 0 via independent sample *t-*tests. All comparisons in each group are further adjusted using the FDR method). For detailed statistics, see TableS1.

**Extended Figure 4.**
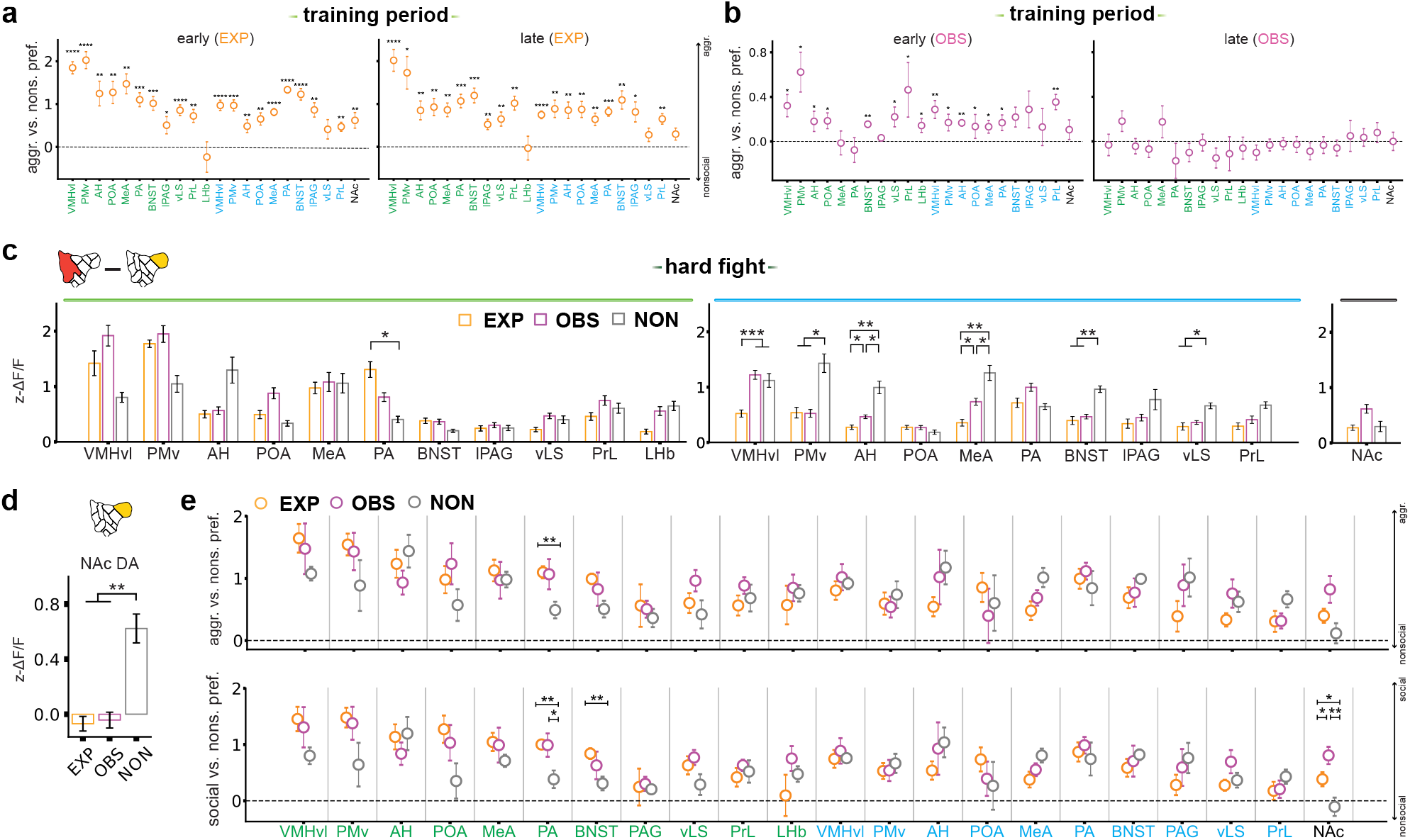
Social versus nonsocial preference in neural activity during aggression and observation. **a**, Preference in social versus nonsocial behavior for each brain area in early (left) versus late (right) days of the training period, for EXP individuals. Social versus nonsocial preference for each population is computed as the logarithm of mean z-ΔF/F during the aggression supercluster divided by mean z-ΔF/F during nonsocial clusters. Values above 0 indicate higher activity during aggression behaviors compared to nonsocial behaviors. (Each point represents mean preference ± SEM. 1-sample *t*-tests or Wilcoxon rank-sum tests relative to 0 pending normality. All comparisons are FDR-corrected). **b**, Same as (**a**) but for OBS individuals. **c**, Average z-scored ΔF/F for excitatory (left), inhibitory (middle) and dopaminergic (right) activity, taken as the difference in z-scored ΔF/F between aggression clusters and nonsocial clusters in the hard fight. (Data are presented as mean ± SEM. One-way ANOVA test followed by post hoc independent samples *t*-tests, or Kruskal-Wallis test followed by post hoc Mann-Whitney U tests. All group comparisons are FDR corrected). **d**, Average NAc dopamine z-ΔF/F aligned to nonsocial clusters in the hard fight. Activity is normalized by the mean ΔF/F of the entire behavior space. (One way ANOVA followed by post hoc independent samples *t*-tests and FDR correction). **e**, Aggression (clusters 1, 2, 3, and 7) versus nonsocial preference (top) and social (clusters 1-10) versus nonsocial preference for each population in the hard fight. (Pending normality, one-way ANOVA tests followed by post-hoc independent samples *t*-tests, or Kruskal-Wallis tests followed by post-hoc Mann-Whitney U tests. All group comparisons are FDR corrected). For detailed statistics, see TableS1.

**Extended Figure 5.**
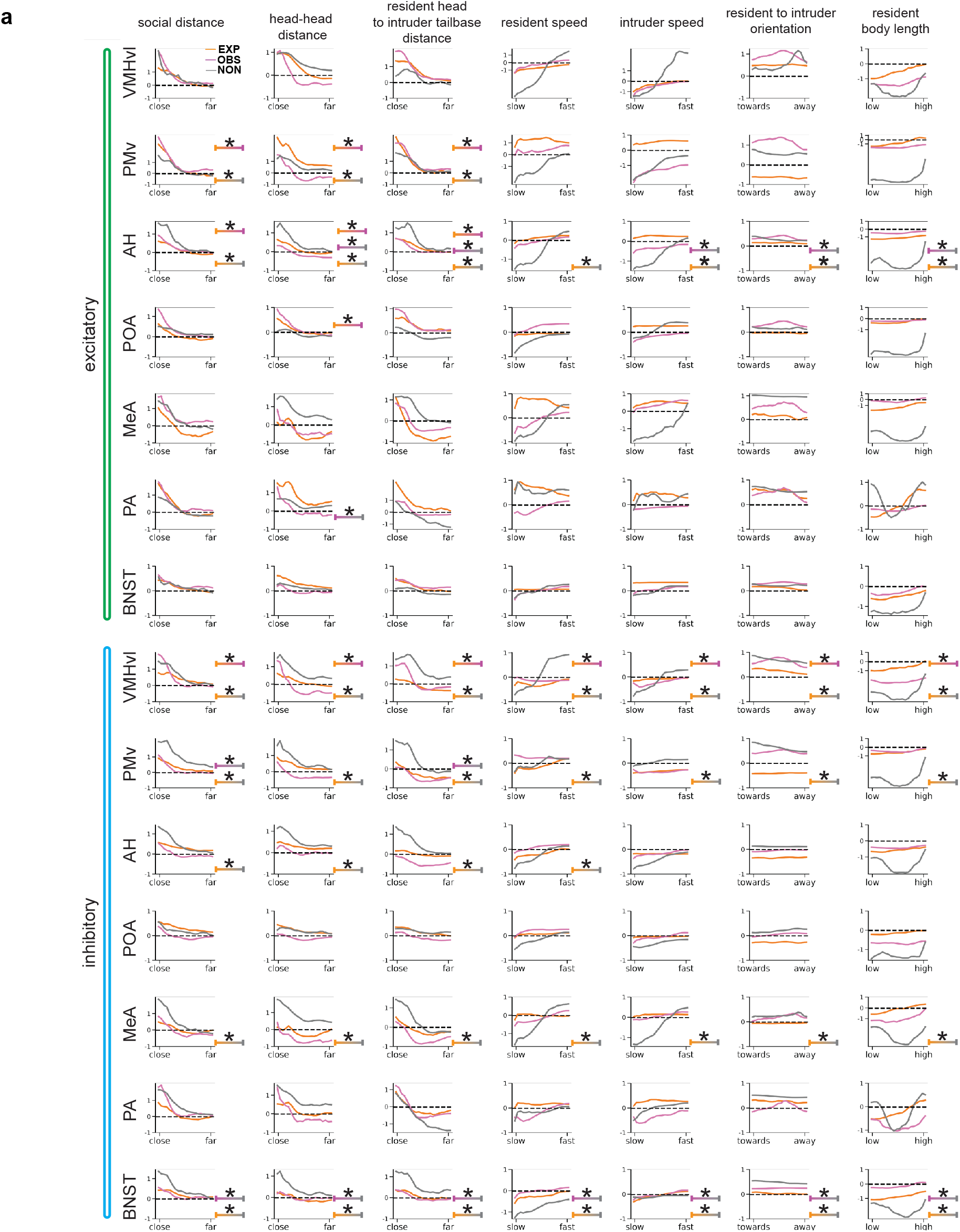
Behavior feature encoding during the hard fight. **A**, Average group-specific tuning curves showing the encoding of behavioral features in hypothalamic and amygdala regions during the hard fight. Each tuning curve represents a set of GLM weights that is fit for each feature and each region independently, while leaving out one animal at a time. A total of 7 features (columns) x 7 regions with E & I signals (rows) are shown for EXP (in orange), OBS (in pink), and NON (in gray), respectively. Pairwise group differences are computed using paired t-tests of r squared predictions on the left out animal with their own groups’ weights versus a different groups’ weight, iteratively across all animals (significance shown for p<0.05). For detailed statistics, see TableS1.

## Supplementary Table 1

Each bordered section corresponds to one or multiple panels within an individual figure using the same statistical method. Information outlined in each section contains the figure number (first column), a description of the comparisons (second column), sample sizes (third column), statistical test employed (fourth column), outcome with relevant statistics and p-values (fifth column) and post-hoc comparisons or adjustments (sixth column). Prior to any statistical test detailed in the fourth column, a Shapiro-Wilks test was used to determine normality of the relevant observations, and the type of statistical test used was adjusted to fit the normality of observations being compared. Final reported p-values are included in either the “Outcome” or “Post-hoc” columns, pending necessity for multiple comparisons corrections. Corrections to p-values reported in the “Outcome” column are detailed in the “Post-hoc” column, and the type of multiple comparisons test is indicated in the first row of the same column within each bordered section of the table. P-values in the manuscript are reported as follows: *p < 0.05, **p<0.01, ***p<0.001, ****p<0.0001, N.S. if not significant. t statistics for linear models correspond to the Wald test and the reported p-value corresponds to the fixed predictor of interest. (H statistic is from Kruskal-Wallis H-test, KS statistic is from two-sample Kolmogorov Smirnov test, F statistic is from one-way ANOVA test, t statistic is from independent samples or related samples t-tests, U statistic is from Mann-Whitney U tests, W statistic is from Wilcoxon rank-sum test).

**Supplementary Table 2.** Coordinates and measurements used for viral deliveries and fiber implantations. Anterior-posterior (AP) coordinates are determined relative to the posterior edge of the rostral rhinal vein. All coordinates are expressed in millimeters. Implants were lowered to the depth of the VMHvl and thus the length of each fiber is determined relative to this depth.

| Brain area | AP | ML | Virus DV | Fiber DV | Fiber implant length |
| --- | --- | --- | --- | --- | --- |
| PrL | -1.4 | 0.35 | -1.9 | -1.7 | 8.28 |
| NAc | -1.8 | 1.15 | -3.85 | -3.7 | 10.38 |
| vLS | -2.6 | 0.6 | -3.45 | -3.25 | 9.83 |
| POA | -3 | -0.4 | -4.9 | -4.75 | 11.28 |
| BNST | -3.3 | 0.85 | -3.95 | -3.75 | 10.08 |
| AH | -3.7 | -0.5 | -5 | -4.8 | 11.03 |
| VMHvl | -4.65 | 0.72 | -5.7 | -5.55 | 11.83 |
| LHb | -4.7 | -0.33 | -2.85 | -2.65 | 8.93 |
| MeA | -4.7 | 2.25 | -5.1 | -4.9 | 11.43 |
| PA | -5.5 | -2.4 | -5.2 | -4.95 | 11.48 |
| PMv | -5.5 | -0.58 | -5.7 | -5.5 | 11.83 |
| IPAG | -7.8 | 0.5 | -1.8 | -1.65 | 8.78 |

## Supplementary Video 1

Example behavior for aggression experience (EXP, left), observation (OBS, middle), and nonsocial control (NON, right). Observer animal in OBS is positioned on a platform to the right and above the interacting animals, and is also filmed with an “observer camera”.

## Supplementary Video 2

Example behaviors from unsupervised clusters (Fig. 1i).

## Notes

### Competing Interest Statement

The authors have declared no competing interest.

