## Supplemental Table 1 for "Aggression experience and observation promote shared behavioral and neural changes"

**Supplementary Data Table 1: Statistical significance testing**

Each bordered section corresponds to one or multiple panels within an individual figure using the same statistical method. Prior to any statistical test outlined in this table, Shapiro-Wilks tests were used to determine normality of the relevant observations, and the type of statistical test used was adjusted to fit the normality of each. Final reported p-values are included in either the “Outcome” or “Post-hoc” columns, pending necessity for multiple comparisons corrections. Corrections to p-values reported in the “Outcome” column are detailed in the “Post-hoc” column, and the type of multiple comparisons test is indicated in the first row of the same column within each bordered section of the table. P-values in the manuscript are reported as follows: *p < 0.05, **p<0.01, ***p<0.001, ****p<0.0001, N.S. if not significant. t statistics for linear models correspond to the Wald test and the reported p-value corresponds to the fixed predictor of interest. (H statistic is from Kruskal-Wallis H-test, KS statistic is from two-sample Kolmogorov Smirnov test, F statistic is from one-way ANOVA test, t statistic is from independent samples or related samples t-tests, U statistic is from Mann-Whitney U tests, W statistic is from Wilcoxon rank-sum test).

| Figure # | Fig description | Sample size | Test | Outcome | Post-hoc |
|---|---|---|---|---|---|
| Fig 1e | *% Time spent attacking for all groups* | EXP = 30 (3 for each EXP subject)<br>OBS = 21 (3 for each OBS subject)<br>NON = 18 (3 for each NON subject) | *Kruskal-Wallis H-test*<br>main effect (group) | H = 2.713572, p = 0.257487 | N/A |
| Fig 1f | *Attack latency distributions in early versus late training period* | early = 60 (6 for each EXP subject)<br>late = 60 (6 for each EXP subject) | two-sample KS test | KS statistic = 0.317, p = 0.0046 | N/A |
| Fig 1g | *Attack latency distributions in hard fight sessions* | EXP = 30 (3 for each EXP subject)<br>OBS = 21 (3 for each OBS subject)<br>NON = 18 (3 for each NON subject) | | | *Benjamini-Hochberg FDR adjustment* |
| | | | two-sample KS test (EXP, NON) | KS statistic = 0.444, p = 0.0168 | p = 0.0263 |
| | | | two sample KS test (OBS, NON) | KS statistic = 0.452, p = 0.0263 | p = 0.0263 |
| Fig 1k | *Transition probabilities in behavior space* | N = 10,000 permutations for each pairwise transition probability | *Permutation test on mean transition probability* | | *Bonferroni adjustment* |
| | | | Permutation test: (0, 1) | p = 0.000 | p = 0.000 |
| | | | Permutation test: (0, 2) | p = 0.000 | p = 0.000 |
| | | | Permutation test: (0, 3) | p = 1.000 | p = 1.000 |
| | | | Permutation test: (0, 4) | p = 1.000 | p = 1.000 |
| | | | Permutation test: (0, 5) | p = 1.000 | p = 1.000 |
| | | | Permutation test: (0, 6) | p = 1.000 | p = 1.000 |
| | | | Permutation test: (0, 7) | p = 1.000 | p = 1.000 |
| | | | Permutation test: (0, 8) | p = 1.000 | p = 1.000 |

| | | | |
|---|---|---|---|
| | Permutation test: (0, 9) | p = 1.000 | p = 1.000 |
| | Permutation test: (0, 10) | p = 1.000 | p = 1.000 |
| | Permutation test: (0, 11) | p = 1.000 | p = 1.000 |
| | Permutation test: (0, 12) | p = 1.000 | p = 1.000 |
| | Permutation test: (1, 0) | p = 0.000 | p = 0.000 |
| | Permutation test: (1, 2) | p = 0.000 | p = 0.000 |
| | Permutation test: (1, 3) | p = 0.000 | p = 0.000 |
| | Permutation test: (1, 4) | p = 1.000 | p = 1.000 |
| | Permutation test: (1, 5) | p = 1.000 | p = 1.000 |
| | Permutation test: (1, 6) | p = 1.000 | p = 1.000 |
| | Permutation test: (1, 7) | p = 1.000 | p = 1.000 |
| | Permutation test: (1, 8) | p = 1.000 | p = 1.000 |
| | Permutation test: (1, 9) | p = 1.000 | p = 1.000 |
| | Permutation test: (1, 10) | p = 1.000 | p = 1.000 |
| | Permutation test: (1, 11) | p = 1.000 | p = 1.000 |
| | Permutation test: (1, 12) | p = 1.000 | p = 1.000 |
| | Permutation test: (2, 0) | p = 0.000 | p = 0.000 |
| | Permutation test: (2, 1) | p = 0.000 | p = 0.000 |
| | Permutation test: (2, 3) | p = 0.000 | p = 0.000 |
| | Permutation test: (2, 4) | p = 1.000 | p = 1.000 |
| | Permutation test: (2, 5) | p = 0.000 | p = 0.000 |
| | Permutation test: (2, 6) | p = 1.000 | p = 1.000 |
| | Permutation test: (2, 7) | p = 0.000 | p = 0.000 |
| | Permutation test: (2, 8) | p = 1.000 | p = 1.000 |
| | Permutation test: (2, 9) | p = 1.000 | p = 1.000 |
| | Permutation test: (2, 10) | p = 1.000 | p = 1.000 |
| | Permutation test: (2, 11) | p = 1.000 | p = 1.000 |
| | Permutation test: (2, 12) | p = 1.000 | p = 1.000 |
| | Permutation test: (3, 0) | p = 1.000 | p = 1.000 |
| | Permutation test: (3, 1) | p = 0.000 | p = 0.000 |
| | Permutation test: (3, 2) | p = 0.000 | p = 0.000 |
| | Permutation test: (3, 4) | p = 1.000 | p = 1.000 |
| | Permutation test: (3, 5) | p = 1.000 | p = 1.000 |
| | Permutation test: (3, 6) | p = 0.000 | p = 0.000 |
| | Permutation test: (3, 7) | p = 0.000 | p = 0.000 |

| | | | |
|---|---|---|---|
| | Permutation test: (3, 8) | p = 1.000 | p = 1.000 |
| | Permutation test: (3, 9) | p = 1.000 | p = 1.000 |
| | Permutation test: (3, 10) | p = 1.000 | p = 1.000 |
| | Permutation test: (3, 11) | p = 1.000 | p = 1.000 |
| | Permutation test: (3, 12) | p = 1.000 | p = 1.000 |
| | Permutation test: (4, 0) | p = 1.000 | p = 1.000 |
| | Permutation test: (4, 1) | p = 1.000 | p = 1.000 |
| | Permutation test: (4, 2) | p = 1.000 | p = 1.000 |
| | Permutation test: (4, 3) | p = 0.496 | p = 1.000 |
| | Permutation test: (4, 5) | p = 1.000 | p = 1.000 |
| | Permutation test: (4, 6) | p = 0.000 | p = 0.000 |
| | Permutation test: (4, 7) | p = 1.000 | p = 1.000 |
| | Permutation test: (4, 8) | p = 1.000 | p = 1.000 |
| | Permutation test: (4, 9) | p = 1.000 | p = 1.000 |
| | Permutation test: (4, 10) | p = 1.000 | p = 1.000 |
| | Permutation test: (4, 11) | p = 1.000 | p = 1.000 |
| | Permutation test: (4, 12) | p = 1.000 | p = 1.000 |
| | Permutation test: (5, 0) | p = 1.000 | p = 1.000 |
| | Permutation test: (5, 1) | p = 1.000 | p = 1.000 |
| | Permutation test: (5, 2) | p = 0.175 | p = 1.000 |
| | Permutation test: (5, 3) | p = 1.000 | p = 1.000 |
| | Permutation test: (5, 4) | p = 1.000 | p = 1.000 |
| | Permutation test: (5, 6) | p = 1.000 | p = 1.000 |
| | Permutation test: (5, 7) | p = 1.000 | p = 1.000 |
| | Permutation test: (5, 8) | p = 1.000 | p = 1.000 |
| | Permutation test: (5, 9) | p = 0.000 | p = 0.000 |
| | Permutation test: (5, 10) | p = 0.000 | p = 0.000 |
| | Permutation test: (5, 11) | p = 1.000 | p = 1.000 |
| | Permutation test: (5, 12) | p = 1.000 | p = 1.000 |
| | Permutation test: (6, 0) | p = 1.000 | p = 1.000 |
| | Permutation test: (6, 1) | p = 1.000 | p = 1.000 |
| | Permutation test: (6, 2) | p = 1.000 | p = 1.000 |
| | Permutation test: (6, 3) | p = 0.000 | p = 0.000 |
| | Permutation test: (6, 4) | p = 0.000 | p = 0.000 |
| | Permutation test: (6, 5) | p = 1.000 | p = 1.000 |

| | | |
|---|---|---|
| Permutation test: (6, 7) | p = 0.000 | p = 0.000 |
| Permutation test: (6, 8) | p = 0.000 | p = 0.000 |
| Permutation test: (6, 9) | p = 1.000 | p = 1.000 |
| Permutation test: (6, 10) | p = 1.000 | p = 1.000 |
| Permutation test: (6, 11) | p = 1.000 | p = 1.000 |
| Permutation test: (6, 12) | p = 1.000 | p = 1.000 |
| Permutation test: (7, 0) | p = 1.000 | p = 1.000 |
| Permutation test: (7, 1) | p = 1.000 | p = 1.000 |
| Permutation test: (7, 2) | p = 0.000 | p = 0.051 |
| Permutation test: (7, 3) | p = 0.003 | p = 0.490 |
| Permutation test: (7, 4) | p = 1.000 | p = 1.000 |
| Permutation test: (7, 5) | p = 1.000 | p = 1.000 |
| Permutation test: (7, 6) | p = 0.000 | p = 0.000 |
| Permutation test: (7, 8) | p = 0.000 | p = 0.000 |
| Permutation test: (7, 9) | p = 0.000 | p = 0.000 |
| Permutation test: (7, 10) | p = 1.000 | p = 1.000 |
| Permutation test: (7, 11) | p = 1.000 | p = 1.000 |
| Permutation test: (7, 12) | p = 1.000 | p = 1.000 |
| Permutation test: (8, 0) | p = 1.000 | p = 1.000 |
| Permutation test: (8, 1) | p = 1.000 | p = 1.000 |
| Permutation test: (8, 2) | p = 1.000 | p = 1.000 |
| Permutation test: (8, 3) | p = 1.000 | p = 1.000 |
| Permutation test: (8, 4) | p = 1.000 | p = 1.000 |
| Permutation test: (8, 5) | p = 1.000 | p = 1.000 |
| Permutation test: (8, 6) | p = 0.000 | p = 0.000 |
| Permutation test: (8, 7) | p = 0.000 | p = 0.000 |
| Permutation test: (8, 9) | p = 0.000 | p = 0.000 |
| Permutation test: (8, 10) | p = 1.000 | p = 1.000 |
| Permutation test: (8, 11) | p = 1.000 | p = 1.000 |
| Permutation test: (8, 12) | p = 1.000 | p = 1.000 |
| Permutation test: (9, 0) | p = 1.000 | p = 1.000 |
| Permutation test: (9, 1) | p = 1.000 | p = 1.000 |
| Permutation test: (9, 2) | p = 1.000 | p = 1.000 |
| Permutation test: (9, 3) | p = 1.000 | p = 1.000 |
| Permutation test: (9, 4) | p = 1.000 | p = 1.000 |

| | | | |
|---|---|---|---|
| | Permutation test: (9, 5) | p = 0.000 | p = 0.000 |
| | Permutation test: (9, 6) | p = 1.000 | p = 1.000 |
| | Permutation test: (9, 7) | p = 0.000 | p = 0.000 |
| | Permutation test: (9, 8) | p = 0.000 | p = 0.000 |
| | Permutation test: (9, 10) | p = 0.000 | p = 0.000 |
| | Permutation test: (9, 11) | p = 1.000 | p = 1.000 |
| | Permutation test: (9, 12) | p = 1.000 | p = 1.000 |
| | Permutation test: (10, 0) | p = 1.000 | p = 1.000 |
| | Permutation test: (10, 1) | p = 1.000 | p = 1.000 |
| | Permutation test: (10, 2) | p = 1.000 | p = 1.000 |
| | Permutation test: (10, 3) | p = 1.000 | p = 1.000 |
| | Permutation test: (10, 4) | p = 1.000 | p = 1.000 |
| | Permutation test: (10, 5) | p = 0.030 | p = 1.000 |
| | Permutation test: (10, 6) | p = 1.000 | p = 1.000 |
| | Permutation test: (10, 7) | p = 1.000 | p = 1.000 |
| | Permutation test: (10, 8) | p = 1.000 | p = 1.000 |
| | Permutation test: (10, 9) | p = 0.000 | p = 0.000 |
| | Permutation test: (10, 11) | p = 1.000 | p = 1.000 |
| | Permutation test: (10, 12) | p = 0.000 | p = 0.000 |
| | Permutation test: (11, 0) | p = 1.000 | p = 1.000 |
| | Permutation test: (11, 1) | p = 1.000 | p = 1.000 |
| | Permutation test: (11, 2) | p = 1.000 | p = 1.000 |
| | Permutation test: (11, 3) | p = 1.000 | p = 1.000 |
| | Permutation test: (11, 4) | p = 1.000 | p = 1.000 |
| | Permutation test: (11, 5) | p = 1.000 | p = 1.000 |
| | Permutation test: (11, 6) | p = 1.000 | p = 1.000 |
| | Permutation test: (11, 7) | p = 1.000 | p = 1.000 |
| | Permutation test: (11, 8) | p = 1.000 | p = 1.000 |
| | Permutation test: (11, 9) | p = 1.000 | p = 1.000 |
| | Permutation test: (11, 10) | p = 0.426 | p = 1.000 |
| | Permutation test: (11, 12) | p = 0.000 | p = 0.000 |
| | Permutation test: (12, 0) | p = 1.000 | p = 1.000 |
| | Permutation test: (12, 1) | p = 1.000 | p = 1.000 |
| | Permutation test: (12, 2) | p = 1.000 | p = 1.000 |
| | Permutation test: (12, 3) | p = 1.000 | p = 1.000 |

| | | | | | |
|---|---|---|---|---|---|
| | | | Permutation test: (12, 4) | p = 1.000 | p = 1.000 |
| | | | Permutation test: (12, 5) | p = 1.000 | p = 1.000 |
| | | | Permutation test: (12, 6) | p = 1.000 | p = 1.000 |
| | | | Permutation test: (12, 7) | p = 1.000 | p = 1.000 |
| | | | Permutation test: (12, 8) | p = 1.000 | p = 1.000 |
| | | | Permutation test: (12, 9) | p = 1.000 | p = 1.000 |
| | | | Permutation test: (12, 10) | p = 0.000 | p = 0.000 |
| | | | Permutation test: (12, 11) | p = 0.000 | p = 0.000 |
| Fig 1m | *Group differences in* | EXP = 10 | *one-way ANOVA (F) or Kruskal-Wallis (H) test* | | *Pairwise comparison, Benjamini-Hochberg FDR adjustment* |
| | *occupancy across behaviors* | OBS = 7 | main effect cluster 1 (group) | H = 0.865424, p = 0.648747 | n.s. |
| | | NON = 6 | main effect cluster 2 (group) | F = 2.864476, p = 0.080552 | n.s. |
| | | | main effect cluster 3 (group) | F = 12.556902, p = 0.000293 | EXP vs NON; t = 4.229157, p = 0.000841<br>EXP vs OBS; t = 3.491950, p = 0.001639<br>OBS vs NON; t = 1.195960, p = 0.085619 |
| | | | main effect cluster 4 (group) | F = 11.544556, p = 0.000464 | EXP vs NON; t = 4.999954, p = 0.000584<br>EXP vs OBS; t = 2.321549, p = 0.036792<br>OBS vs NON; t = 2.375513, p = 0.036792 |
| | | | main effect cluster 5 (group) | H = 6.918144, p = 0.031459 | EXP vs NON; U = 55.000000, p = 0.009491<br>EXP vs OBS; U = 40.000000, p = 0.446044<br>OBS vs NON; U = 32.500000, p = 0.115582 |
| | | | main effect cluster 6 (group) | F = 2.864251, p = 0.080567 | n.s. |
| | | | main effect cluster 7 (group) | H = 4.251449, p = 0.119346 | n.s. |
| | | | main effect cluster 8 (group) | F = 6.029993, p = 0.008926 | EXP vs NON; t = 3.112786, p = 0.015272<br>EXP vs OBS; t = 1.982946, p = 0.065994<br>OBS vs NON; t = 1.619974, p = 0.089018 |
| | | | main effect cluster 9 (group) | F = 2.351370, p = 0.121018 | n.s. |
| | | | main effect cluster 10 (group) | F = 6.142082, p = 0.008325 | EXP vs NON; t = -3.409591, p = 0.008463<br>EXP vs OBS; t = -2.033069, p = 0.060136<br>OBS vs NON; t = -1.336031, p = 0.139014 |
| | | | main effect cluster 11 (group) | F = 2.899258, p = 0.078407 | n.s. |

| | | | | | |
|---|---|---|---|---|---|
| | | | main effect cluster 12 (group) | F = 1.488946, p = 0.249573 | n.s. |
| | | | main effect cluster 13 (group) | H = 8.221946, p = 0.016392 | EXP vs NON; U = 5.000000, p = 0.009491<br>EXP vs OBS; U = 28.000000, p = 0.357466<br>OBS vs NON; U = 6.000000, p = 0.034965 |
| Fig 1n | *Occupancy-based decoding*<br>*of group identity* | N = 1,000 permutations | *Permutation test on F1 score* | F1 = 0.68, p = 0.000000 | N/A |
| Fig 1o | *JS divergence of occupancy*<br>*vectors* | OBS vs. EXP = 630<br>OBS vs. NON = 378 | two-sample KS test (OBS vs. EXP, OBS vs. NON) | KS = 0.1153, p = 0.0034 | N/A |
| Fig 1q | *Transition probability-based*<br>*decoding of group identity* | N = 1,000 permutations | *Permutation test on F1 score* | F1 = 0.52, p = 0.000000 | |
| Fig 1r | *JS divergence of transition*<br>*probability vectors* | OBS vs. EXP = 630<br>OBS vs. NON = 378 | two-sample KS test (OBS vs. EXP, OBS vs. NON) | KS = 0.2587<br>p = 0.0000000000000228 | N/A |
| Fig 2g &<br>Fig 2h | *Neural activation of SBN populations*<br>*in EXP and OBS animals* | | *Independence t-test (t) or Mann-Whitney U test (U)* | | *Benjamini-Hochberg FDR adjustments* |
| | | EXP = 12, NON = 6 | pre-attack VMHvl E (EXP) | EXP vs NON; t = 3.546959, p = 0.003577 | p = 0.022816 |
| | | EXP = 8, NON = 6 | pre-attack PMv E (EXP) | EXP vs NON; t = 3.766657, p = 0.003118 | p = 0.022816 |
| | | EXP = 10, NON = 6 | pre-attack AH E (EXP) | EXP vs NON; t = 0.398173, p = 0.696965 | p = 0.644692 |
| | | EXP = 11, NON = 6 | pre-attack POA E (EXP) | EXP vs NON; t = 1.801888, p = 0.096721 | p = 0.197460 |
| | | EXP = 11, NON = 6 | pre-attack MeA E (EXP) | EXP vs NON; t = 4.407648, p = 0.000708 | p = 0.008727 |
| | | EXP = 9, NON = 6 | pre-attack PA E (EXP) | EXP vs NON; U = 37.000000, p = 0.022145 | p = 0.068279 |
| | | EXP = 10, NON = 6 | pre-attack BNST E (EXP) | EXP vs NON; t = 2.226208, p = 0.045923 | p = 0.121368 |
| | | EXP = 8, NON = 6 | pre-attack PAG E (EXP) | EXP vs NON; t = -0.005127, p = 0.996001 | p = 0.801131 |
| | | EXP = 12, NON = 6 | pre-attack vLS E (EXP) | EXP vs NON; t = 0.254896, p = 0.802792 | p = 0.690774 |
| | | EXP = 11, NON = 6 | pre-attack PrL E (EXP) | EXP vs NON; t = 2.081704, p = 0.057695 | p = 0.142314 |
| | | EXP = 9, NON = 6 | pre-attack LHb E (EXP) | EXP vs NON; t = -1.672664, p = 0.122563 | p = 0.222844 |
| | | EXP = 12, NON = 6 | pre-attack VMHvl I (EXP) | EXP vs NON; t = 3.183026, p = 0.007199 | p = 0.031296 |
| | | EXP = 8, NON = 6 | pre-attack PMv I (EXP) | EXP vs NON; t = 2.023977, p = 0.067951 | p = 0.157136 |
| | | EXP = 10, NON = 6 | pre-attack AH I (EXP) | EXP vs NON; t = -0.719047, p = 0.484837 | p = 0.493590 |
| | | EXP = 11, NON = 6 | pre-attack POA I (EXP) | EXP vs NON; t = 1.189749, p = 0.257153 | p = 0.339809 |

| | | | |
|---|---|---|---|
| EXP = 11, NON = 6 | pre-attack MeA I (EXP) | EXP vs NON; t = 3.448993, p = 0.004316 | p = 0.022816 |
| EXP = 9, NON = 6 | pre-attack PA I (EXP) | EXP vs NON; t = 3.642903, p = 0.003868 | p = 0.022816 |
| EXP = 10, NON = 6 | pre-attack BNST I (EXP) | EXP vs NON; t = 1.371174, p = 0.195418 | p = 0.277268 |
| EXP = 8, NON = 6 | pre-attack PAG I (EXP) | EXP vs NON; t = 1.653379, p = 0.126479 | p = 0.222844 |
| EXP = 12, NON = 6 | pre-attack vLS I (EXP) | EXP vs NON; t = 1.043401, p = 0.315778 | p = 0.376896 |
| EXP = 11, NON = 6 | pre-attack PrL I (EXP) | EXP vs NON; t = 4.814569, p = 0.000338 | p = 0.008727 |
| EXP = 11, NON = 6 | pre-attack NAc DA (EXP) | EXP vs NON; t = 4.709143, p = 0.000506 | p = 0.008727 |
| EXP = 12, NON = 6 | post-attack VMHvl E (EXP) | EXP vs NON; t = 5.653069, p = 0.000079 | p = 0.000841 |
| EXP = 8, NON = 6 | post-attack PMv E (EXP) | EXP vs NON; t = 4.704095, p = 0.000646 | p = 0.001534 |
| EXP = 10, NON = 6 | post-attack AH E (EXP) | EXP vs NON; t = 1.823951, p = 0.091223 | p = 0.049521 |
| EXP = 11, NON = 6 | post-attack POA E (EXP) | EXP vs NON; t = 4.438049, p = 0.000810 | p = 0.001709 |
| EXP = 11, NON = 6 | post-attack MeA E (EXP) | EXP vs NON; t = 5.169480, p = 0.000180 | p = 0.000857 |
| EXP = 9, NON = 6 | post-attack PA E (EXP) | EXP vs NON; U = 38.000000, p = 0.013986 | p = 0.013986 |
| EXP = 10, NON = 6 | post-attack BNST E (EXP) | EXP vs NON; t = 3.050580, p = 0.010074 | p = 0.010634 |
| EXP = 8, NON = 6 | post-attack PAG E (EXP) | EXP vs NON; t = 3.492130, p = 0.005040 | p = 0.006384 |
| EXP = 12, NON = 6 | post-attack vLS E (EXP) | EXP vs NON; t = 2.278653, p = 0.040219 | p = 0.026351 |
| EXP = 11, NON = 6 | post-attack PrL E (EXP) | EXP vs NON; t = 2.524605, p = 0.025382 | p = 0.019648 |
| EXP = 9, NON = 6 | post-attack LHb E (EXP) | EXP vs NON; t = -0.663081, p = 0.520931 | p = 0.230179 |
| EXP = 12, NON = 6 | post-attack VMHvl I (EXP) | EXP vs NON; t = 4.949127, p = 0.000266 | p = 0.001010 |
| EXP = 8, NON = 6 | post-attack PMv I (EXP) | EXP vs NON; t = 3.154741, p = 0.009164 | p = 0.010243 |
| EXP = 10, NON = 6 | post-attack AH I (EXP) | EXP vs NON; t = 4.623384, p = 0.000477 | p = 0.001295 |
| EXP = 11, NON = 6 | post-attack POA I (EXP) | EXP vs NON; t = 2.503360, p = 0.027744 | p = 0.019648 |
| EXP = 11, NON = 6 | post-attack MeA I (EXP) | EXP vs NON; t = 5.346276, p = 0.000133 | p = 0.000841 |
| EXP = 9, NON = 6 | post-attack PA I (EXP) | EXP vs NON; t = 4.090730, p = 0.001787 | p = 0.003127 |
| EXP = 10, NON = 6 | post-attack BNST I (EXP) | EXP vs NON; t = 2.695010, p = 0.019488 | p = 0.018514 |
| EXP = 8, NON = 6 | post-attack PAG I (EXP) | EXP vs NON; t = 2.628597, p = 0.023466 | p = 0.019594 |
| EXP = 12, NON = 6 | post-attack vLS I (EXP) | EXP vs NON; t = 2.573311, p = 0.023148 | p = 0.019594 |
| EXP = 11, NON = 6 | post-attack PrL I (EXP) | EXP vs NON; t = 5.356345, p = 0.000131 | p = 0.000841 |
| EXP = 11, NON = 6 | post-attack NAc DA (EXP) | EXP vs NON; t = 4.908674, p = 0.000361 | p = 0.001142 |
| OBS = 7, NON = 6 | pre-attack VMHvl E (OBS) | OBS vs NON; t = 0.708142, p = 0.493590 | p = 0.493590 |
| OBS = 7, NON = 6 | pre-attack PMv E (OBS) | OBS vs NON; t = 0.266847, p = 0.794523 | p = 0.690774 |
| OBS = 7, NON = 6 | pre-attack AH E(OBS) | OBS vs NON; t = -1.093386, p = 0.297590 | p = 0.367028 |

| | | | |
|---|---|---|---|
| OBS = 7, NON = 6 | pre-attack POA E(OBS) | OBS vs NON; t = -0.033180, p = 0.974126 | p = 0.800948 |
| OBS = 7, NON = 6 | pre-attack MeA E(OBS) | OBS vs NON; t = 1.548392, p = 0.149801 | p = 0.240984 |
| OBS = 7, NON = 6 | pre-attack PA E (OBS) | OBS vs NON; U = 29.000000, p = 0.294872 | p = 0.367028 |
| OBS = 7, NON = 6 | pre-attack BNST E (OBS) | OBS vs NON; t = 0.108212, p = 0.915776 | p = 0.770085 |
| OBS = 7, NON = 6 | pre-attack PAG E (OBS) | OBS vs NON; t = -2.666044, p = 0.021949 | p = 0.068279 |
| OBS = 7, NON = 6 | pre-attack vLS E (OBS) | OBS vs NON; t = -0.370535, p = 0.718020 | p = 0.647969 |
| OBS = 7, NON = 6 | pre-attack PrL E (OBS) | OBS vs NON; U = 9.000000, p = 0.101399 | p = 0.197460 |
| OBS = 7, NON = 6 | pre-attack LHb E (OBS) | OBS vs NON; t = 0.939157, p = 0.367815 | p = 0.425286 |
| OBS = 7, NON = 6 | pre-attack VMHvl I (OBS) | OBS vs NON; t = 3.001692, p = 0.012043 | p = 0.044560 |
| OBS = 7, NON = 6 | pre-attack PMv I (OBS) | OBS vs NON; t = -0.791674, p = 0.445287 | p = 0.484577 |
| OBS = 7, NON = 6 | pre-attack AH I (OBS) | OBS vs NON; t = 0.453638, p = 0.658908 | p = 0.625118 |
| OBS = 7, NON = 6 | pre-attack POA I (OBS) | OBS vs NON; t = 0.810347, p = 0.434922 | p = 0.484577 |
| OBS = 7, NON = 6 | pre-attack MeA I (OBS) | OBS vs NON; t = 1.419049, p = 0.183594 | p = 0.277268 |
| OBS = 7, NON = 6 | pre-attack PA I (OBS) | OBS vs NON; t = 2.264686, p = 0.044723 | p = 0.121368 |
| OBS = 7, NON = 6 | pre-attack BNST I (OBS) | OBS vs NON; t = -1.355840, p = 0.202331 | p = 0.277268 |
| OBS = 7, NON = 6 | pre-attack PAG I (OBS) | OBS vs NON; t = -0.747992, p = 0.470155 | p = 0.493590 |
| OBS = 7, NON = 6 | pre-attack vLS I (OBS) | OBS vs NON; t = 1.558344, p = 0.147439 | p = 0.240984 |
| OBS = 7, NON = 6 | pre-attack PrL I (OBS) | OBS vs NON; U = 34.000000, p = 0.073427 | p = 0.159811 |
| OBS = 7, NON = 6 | pre-attack NAc DA (OBS) | OBS vs NON; t = -0.590425, p = 0.566838 | p = 0.551921 |
| OBS = 7, NON = 6 | post-attack VMHvl E (OBS) | OBS vs NON; t = 2.554105, p = 0.026799 | p = 0.019648 |
| OBS = 7, NON = 6 | post-attack PMv E (OBS) | OBS vs NON; U = 40.000000, p = 0.004662 | p = 0.006327 |
| OBS = 7, NON = 6 | post-attack AH E (OBS) | OBS vs NON; t = 2.365633, p = 0.037440 | p = 0.025405 |
| OBS = 7, NON = 6 | post-attack POA E (OBS) | OBS vs NON; t = 1.728840, p = 0.111764 | p = 0.058986 |
| OBS = 7, NON = 6 | post-attack MeA E (OBS) | OBS vs NON; t = 2.531044, p = 0.027921 | p = 0.019648 |
| OBS = 7, NON = 6 | post-attack PA E (OBS) | OBS vs NON; U = 32.000000, p = 0.137529 | p = 0.068765 |
| OBS = 7, NON = 6 | post-attack BNST E (OBS) | OBS vs NON; t = 1.247675, p = 0.238058 | p = 0.115684 |
| OBS = 7, NON = 6 | post-attack PAG E (OBS) | OBS vs NON; t = -1.232248, p = 0.243546 | p = 0.115684 |
| OBS = 7, NON = 6 | post-attack vLS E (OBS) | OBS vs NON; t = 0.991797, p = 0.342612 | p = 0.154991 |
| OBS = 7, NON = 6 | post-attack PrL E (OBS) | OBS vs NON; U = 34.000000, p = 0.073427 | p = 0.042276 |
| OBS = 7, NON = 6 | post-attack LHb E (OBS) | OBS vs NON; t = 2.274915, p = 0.043927 | p = 0.027820 |
| OBS = 7, NON = 6 | post-attack VMHvl I (OBS) | OBS vs NON; t = 4.024096, p = 0.002002 | p = 0.003170 |
| OBS = 7, NON = 6 | post-attack PMv I (OBS) | OBS vs NON; t = 1.017831, p = 0.330623 | p = 0.153215 |

| | | | | | |
|---|---|---|---|---|---|
| | | OBS = 7, NON = 6 | post-attack AH I (OBS) | OBS vs NON; t = 1.614389, p = 0.134738 | p = 0.068765 |
| | | OBS = 7, NON = 6 | post-attack POA I (OBS) | OBS vs NON; t = 2.095193, p = 0.060098 | p = 0.036834 |
| | | OBS = 7, NON = 6 | post-attack MeA I (OBS) | OBS vs NON; t = 2.075681, p = 0.062160 | p = 0.036908 |
| | | OBS = 7, NON = 6 | post-attack PA I (OBS) | OBS vs NON; t = 2.622581, p = 0.023719 | p = 0.019594 |
| | | OBS = 7, NON = 6 | post-attack BNST I (OBS) | OBS vs NON; t = 0.262119, p = 0.798073 | p = 0.329639 |
| | | OBS = 7, NON = 6 | post-attack PAG I (OBS) | OBS vs NON; t = 0.587483, p = 0.568743 | p = 0.245594 |
| | | OBS = 7, NON = 6 | post-attack vLS I (OBS) | OBS vs NON; t = 3.284358, p = 0.007277 | p = 0.008642 |
| | | OBS = 7, NON = 6 | post-attack PrL I (OBS) | OBS vs NON; t = 3.764335, p = 0.003131 | p = 0.004576 |
| | | OBS = 7, NON = 6 | post-attack NAc DA (OBS) | OBS vs NON; t = 0.437867, p = 0.669956 | p = 0.282870 |
| Fig 2i (left) | *Mean Nac - LHb index difference* | EXP = 7, OBS = 6 | *Independence t-test* | EXP vs OBS; t = 3.1749, p = 0.007995 | N/A |
| Fig 2i (right) | *Nac - LHb index distribution difference* | EXP = 22400, OBS = 22400 | *Two-sample Kolmogorov-Smirnov test* | KS = 0.3495, p = 0.0 | N/A |
| Fig 3c (top) & Fig 3d (top) | *Neural activation of SBN populations in EXP and OBS animals* | | *Independence t-test (t) or Mann-Whitney U test (U)* | | *Benjamini-Hochberg FDR adjustments* |
| | | EXP = 12, NON = 6 | VMHvl E (EXP) | EXP vs NON; t = 5.211493, p = 0.000086 | p = 0.000373 |
| | | EXP = 8, NON = 6 | PMv E (EXP) | EXP vs NON; U = 48.000000, p = 0.000666 | p = 0.000761 |
| | | EXP = 10, NON = 6 | AH E (EXP) | EXP vs NON; t = 3.749077, p = 0.002157 | p = 0.001817 |
| | | EXP = 11, NON = 6 | POA E (EXP) | EXP vs NON; t = 4.984689, p = 0.000163 | p = 0.000373 |
| | | EXP = 11, NON = 6 | MeA E (EXP) | EXP vs NON; t = 4.827283, p = 0.000222 | p = 0.000443 |
| | | EXP = 9, NON = 6 | PA E (EXP) | EXP vs NON; t = 4.691299, p = 0.000422 | p = 0.000530 |
| | | EXP = 10, NON = 6 | BNST E (EXP) | EXP vs NON; t = 4.001504, p = 0.001312 | p = 0.001235 |
| | | EXP = 8, NON = 6 | PAG E (EXP) | EXP vs NON; t = 4.327796, p = 0.000983 | p = 0.000983 |
| | | EXP = 12, NON = 6 | vLS E (EXP) | EXP vs NON; t = 4.907418, p = 0.000158 | p = 0.000373 |
| | | EXP = 11, NON = 6 | PrL E (EXP) | EXP vs NON; U = 66.000000, p = 0.000162 | p = 0.000373 |
| | | EXP = 9, NON = 6 | LHb E (EXP) | EXP vs NON; t = 1.644688, p = 0.123985 | p = 0.052001 |
| | | EXP = 12, NON = 6 | VMHvl I (EXP) | EXP vs NON; t = 5.731250, p = 0.000031 | p = 0.000247 |
| | | EXP = 8, NON = 6 | PMv I (EXP) | EXP vs NON; t = 3.911148, p = 0.002068 | p = 0.001817 |
| | | EXP = 10, NON = 6 | AH I (EXP) | EXP vs NON; t = 7.725417, p = 0.000002 | p = 0.000033 |
| | | EXP = 11, NON = 6 | POA I (EXP) | EXP vs NON; t = 2.906609, p = 0.010848 | p = 0.007547 |
| | | EXP = 11, NON = 6 | MeA I (EXP) | EXP vs NON; t = 4.534234, p = 0.000395 | p = 0.000530 |
| | | EXP = 9, NON = 6 | PA I (EXP) | EXP vs NON; t = 4.327025, p = 0.000821 | p = 0.000876 |
| | | EXP = 10, NON = 6 | BNST I (EXP) | EXP vs NON; U = 60.000000, p = 0.000250 | p = 0.000444 |
| | | EXP = 8, NON = 6 | PAG I (EXP) | EXP vs NON; t = 3.353767, p = 0.005740 | p = 0.004373 |

| | | | | | |
|---|---|---|---|---|---|
| | | EXP = 12, NON = 6 | vLS I (EXP) | EXP vs NON; U = 70.000000, p = 0.000431 | p = 0.000530 |
| | | EXP = 11, NON = 6 | PrL I (EXP) | EXP vs NON; U = 66.000000, p = 0.000162 | p = 0.000373 |
| | | EXP = 11, NON = 6 | NAc DA (EXP) | EXP vs NON; t = 2.708287, p = 0.016187 | p = 0.010360 |
| | | OBS = 7, NON = 6 | VMHvl E (OBS) | OBS vs NON; t = 2.797082, p = 0.017365 | p = 0.010485 |
| | | OBS = 7, NON = 6 | PMv E (OBS) | OBS vs NON; U = 40.000000, p = 0.004662 | p = 0.003730 |
| | | OBS = 7, NON = 6 | AH E(OBS) | OBS vs NON; t = 0.723710, p = 0.484351 | p = 0.172214 |
| | | OBS = 7, NON = 6 | POA E(OBS) | OBS vs NON; U = 35.000000, p = 0.051282 | p = 0.025641 |
| | | OBS = 7, NON = 6 | MeA E(OBS) | OBS vs NON; t = 2.073949, p = 0.062346 | p = 0.030229 |
| | | OBS = 7, NON = 6 | PA E (OBS) | OBS vs NON; t = 1.854193, p = 0.090696 | p = 0.042680 |
| | | OBS = 7, NON = 6 | BNST E (OBS) | OBS vs NON; t = 2.651562, p = 0.022523 | p = 0.012013 |
| | | OBS = 7, NON = 6 | PAG E (OBS) | OBS vs NON; t = 1.572755, p = 0.144077 | p = 0.057631 |
| | | OBS = 7, NON = 6 | vLS E (OBS) | OBS vs NON; t = 1.685668, p = 0.119983 | p = 0.051885 |
| | | OBS = 7, NON = 6 | PrL E (OBS) | OBS vs NON; t = 1.321924, p = 0.213024 | p = 0.081152 |
| | | OBS = 7, NON = 6 | LHb E (OBS) | OBS vs NON; U = 37.000000, p = 0.022145 | p = 0.012013 |
| | | OBS = 7, NON = 6 | VMHvl I (OBS) | OBS vs NON; t = 3.028361, p = 0.011483 | p = 0.007655 |
| | | OBS = 7, NON = 6 | PMv I (OBS) | OBS vs NON; t = 1.703519, p = 0.116521 | p = 0.051787 |
| | | OBS = 7, NON = 6 | AH I (OBS) | OBS vs NON; t = 2.263709, p = 0.044800 | p = 0.023122 |
| | | OBS = 7, NON = 6 | POA I (OBS) | OBS vs NON; t = 0.890802, p = 0.392103 | p = 0.142583 |
| | | OBS = 7, NON = 6 | MeA I (OBS) | OBS vs NON; t = 1.745490, p = 0.108731 | p = 0.049706 |
| | | OBS = 7, NON = 6 | PA I (OBS) | OBS vs NON; t = 3.062695, p = 0.010800 | p = 0.007547 |
| | | OBS = 7, NON = 6 | BNST I (OBS) | OBS vs NON; t = 0.944364, p = 0.365265 | p = 0.135912 |
| | | OBS = 7, NON = 6 | PAG I (OBS) | OBS vs NON; t = 1.525225, p = 0.155426 | p = 0.060654 |
| | | OBS = 7, NON = 6 | vLS I (OBS) | OBS vs NON; t = 2.684501, p = 0.021237 | p = 0.012013 |
| | | OBS = 7, NON = 6 | PrL I (OBS) | OBS vs NON; t = 1.652049, p = 0.126753 | p = 0.052001 |
| | | OBS = 7, NON = 6 | NAc DA (OBS) | OBS vs NON; t = -0.226799, p = 0.824740 | p = 0.286866 |
| Fig 3c (bottom) & | *Cosine similarity distributions* | | *Two-sample Kolmogorov Smirnov tests* | | *Benjamini-Hochberg FDR adjustments* |
| Fig 3d (bottom) | *between EXP/OBS activity* | OBS vs. EXP = 84, OBS vs. NON = 42 | VMHvl E | KS = 0.142857, p = 0.603942 | p = 0.219615 |
| | *maps and OBS/NON activity maps* | OBS vs. EXP = 63, OBS vs. NON = 42 | PMv E | KS = 0.089286, p = 0.983038 | p = 0.341926 |
| | | OBS vs. EXP = 70, OBS vs. NON = 42 | AH E | KS = 0.614286, p = 0.000000 | p = 0.000000 |
| | | OBS vs. EXP = 84, OBS vs. NON = 42 | POA E | KS = 0.450216, p = 0.000017 | p = 0.000023 |
| | | OBS vs. EXP = 77, OBS vs. NON = 42 | MeA E | KS = 0.361472, p = 0.001133 | p = 0.000756 |

| | | | | | |
|---|---|---|---|---|---|
| | | OBS vs. EXP = 70, OBS vs. NON = 42 | PA E | KS = 0.603175, p = 0.000000 | p = 0.000000 |
| | | OBS vs. EXP = 70, OBS vs. NON = 42 | BNST E | KS = 0.423810, p = 0.000099 | p = 0.000099 |
| | | OBS vs. EXP = 63, OBS vs. NON = 42 | PAG E | KS = 0.696429, p = 0.000000 | p = 0.000000 |
| | | OBS vs. EXP = 84, OBS vs. NON = 42 | vLS E | KS = 0.333333, p = 0.003441 | p = 0.001835 |
| | | OBS vs. EXP = 77, OBS vs. NON = 42 | PrL E | KS = 0.147186, p = 0.544787 | p = 0.207538 |
| | | OBS vs. EXP = 70, OBS vs. NON = 42 | LHb E | KS = 0.444444, p = 0.000059 | p = 0.000068 |
| | | OBS vs. EXP = 84, OBS vs. NON = 42 | VMHvl I | KS = 0.333333, p = 0.003441 | p = 0.001835 |
| | | OBS vs. EXP = 63, OBS vs. NON = 42 | PMv I | KS = 0.160714, p = 0.520871 | p = 0.207538 |
| | | OBS vs. EXP = 70, OBS vs. NON = 42 | AH I | KS = 0.209524, p = 0.176207 | p = 0.078314 |
| | | OBS vs. EXP = 84, OBS vs. NON = 42 | POA I | KS = 0.367965, p = 0.000863 | p = 0.000628 |
| | | OBS vs. EXP = 77, OBS vs. NON = 42 | MeA I | KS = 0.246753, p = 0.060470 | p = 0.028456 |
| | | OBS vs. EXP = 70, OBS vs. NON = 42 | PA I | KS = 0.539683, p = 0.000000 | p = 0.000001 |
| | | OBS vs. EXP = 70, OBS vs. NON = 42 | BNST I | KS = 0.157143, p = 0.492952 | p = 0.207538 |
| | | OBS vs. EXP = 63, OBS vs. NON = 42 | PAG I | KS = 0.398810, p = 0.000665 | p = 0.000591 |
| | | OBS vs. EXP = 84, OBS vs. NON = 42 | vLS I | KS = 0.369048, p = 0.000796 | p = 0.000628 |
| | | OBS vs. EXP = 77, OBS vs. NON = 42 | PrL I | KS = 0.681818, p = 0.000000 | p = 0.000000 |
| | | OBS vs. EXP = 84, OBS vs. NON = 42 | NAc DA | KS = 0.259740, p = 0.041682 | p = 0.020841 |
| Fig 3e | *Mean similarity* | | *Independence t-test (t) or Mann-Whitney U test (U)* | | *Benjamini-Hochberg FDR adjustments* |
| | *between EXP/OBS activity* | OBS vs. EXP = 84, OBS vs. NON = 42 | VMHvl E | t = 0.412829, p = 0.680445 | p = 0.216505 |
| | *maps and OBS/NON activity maps* | OBS vs. EXP = 63, OBS vs. NON = 42 | PMv E | t = 0.173987, p = 0.862242 | p = 0.262422 |
| | | OBS vs. EXP = 70, OBS vs. NON = 42 | AH E | U = 2406.000000, p = 0.000000 | p = 0.000000 |
| | | OBS vs. EXP = 84, OBS vs. NON = 42 | POA E | U = 2561.000000, p = 0.000000 | p = 0.000000 |

| | | | | | |
|---|---|---|---|---|---|
| | | OBS vs. EXP = 77, OBS vs. NON = 42 | MeA E | U = 1027.000000, p = 0.001045 | p = 0.000732 |
| | | OBS vs. EXP = 70, OBS vs. NON = 42 | PA E | t = -6.839473, p = 0.000000 | p = 0.000000 |
| | | OBS vs. EXP = 70, OBS vs. NON = 42 | BNST E | U = 2103.000000, p = 0.000144 | p = 0.000126 |
| | | OBS vs. EXP = 63, OBS vs. NON = 42 | PAG E | U = 1996.000000, p = 0.000000 | p = 0.000000 |
| | | OBS vs. EXP = 84, OBS vs. NON = 42 | vLS E | U = 2245.000000, p = 0.012895 | p = 0.006447 |
| | | OBS vs. EXP = 77, OBS vs. NON = 42 | PrL E | t = 1.248381, p = 0.214384 | p = 0.078984 |
| | | OBS vs. EXP = 70, OBS vs. NON = 42 | LHb E | U = 2062.000000, p = 0.000001 | p = 0.000001 |
| | | OBS vs. EXP = 84, OBS vs. NON = 42 | VMHvl I | U = 1295.000000, p = 0.015327 | p = 0.007152 |
| | | OBS vs. EXP = 63, OBS vs. NON = 42 | PMv I | U = 1115.000000, p = 0.664056 | p = 0.216505 |
| | | OBS vs. EXP = 70, OBS vs. NON = 42 | AH I | t = 1.429454, p = 0.155708 | p = 0.060553 |
| | | OBS vs. EXP = 84, OBS vs. NON = 42 | POA I | U = 1130.000000, p = 0.006825 | p = 0.003865 |
| | | OBS vs. EXP = 77, OBS vs. NON = 42 | MeA I | U = 1106.000000, p = 0.004529 | p = 0.002882 |
| | | OBS vs. EXP = 70, OBS vs. NON = 42 | PA I | U = 445.000000, p = 0.000000 | p = 0.000000 |
| | | OBS vs. EXP = 70, OBS vs. NON = 42 | BNST I | U = 1636.000000, p = 0.319900 | p = 0.111965 |
| | | OBS vs. EXP = 63, OBS vs. NON = 42 | PAG I | U = 801.000000, p = 0.007178 | p = 0.003865 |
| | | OBS vs. EXP = 84, OBS vs. NON = 42 | vLS I | t = -3.644309, p = 0.000393 | p = 0.000305 |
| | | OBS vs. EXP = 77, OBS vs. NON = 42 | PrL I | t = -8.656354, p = 0.000000 | p = 0.000000 |
| | | OBS vs. EXP = 84, OBS vs. NON = 42 | NAc DA | U = 1350.000000, p = 0.138359 | p = 0.056972 |
| Fig 4b | *Neural activation of SBN populations* | EXP = 72, OBS = 80, NON = 42 | *one-way ANOVA (F) or Kruskal-Wallis (H) test* | | *Pairwise comparison, Bonferroni adjustment* |
| | *in EXP and OBS animals* | | main effect VMHvl E (group) | H = 7.157174, p = 0.027915 | EXP vs NON; U = 1261.000000, p = 0.088485<br>EXP vs OBS; U = 2358.000000, p = 0.053969<br>OBS vs NON; U = 1882.000000, p = 1.000000 |
| | | | main effect PMv E (group) | H = 4.037867, p = 0.132797 | n.s. |

| | | |
|---|---|---|
| main effect AH E (group) | H = 16.891264, p = 0.000215 | EXP vs NON; U = 931.000000, p = 0.000192<br>EXP vs OBS; U = 2594.000000, p = 0.380390<br>OBS vs NON; U = 1321.000000, p = 0.008862 |
| main effect POA E (group) | H = 4.488766, p = 0.105993 | n.s. |
| main effect MeA E (group) | H = 16.666859, p = 0.000240 | EXP vs NON; U = 949.000000, p = 0.000291<br>EXP vs OBS; U = 2319.000000, p = 0.036790<br>OBS vs NON; U = 1496.000000, p = 0.101909 |
| main effect PA E (group) | H = 1.443894, p = 0.485806 | n.s. |
| main effect BNST E (group) | H = 1.097674, p = 0.577621 | n.s. |
| main effect PAG E (group) | H = 4.249910, p = 0.119438 | n.s. |
| main effect vLS E (group) | H = 13.370737, p = 0.001249 | EXP vs NON; U = 1111.000000, p = 0.007981<br>EXP vs OBS; U = 2114.000000, p = 0.003672<br>OBS vs NON; U = 1970.000000, p = 1.000000 |
| main effect PrL E (group) | H = 5.652253, p = 0.059242 | n.s. |
| main effect LHb E (group) | H = 26.219010, p = 0.000002 | EXP vs NON; U = 636.000000, p = 0.000004<br>EXP vs OBS; U = 1532.000000, p = 0.000112<br>OBS vs NON; U = 1799.000000, p = 1.000000 |
| main effect VMHvl I (group) | H = 24.862979, p = 0.000004 | EXP vs NON; U = 1046.000000, p = 0.002312<br>EXP vs OBS; U = 1661.000000, p = 0.000004<br>OBS vs NON; U = 2012.000000, p = 1.000000 |
| main effect PMv I (group) | H = 13.206616, p = 0.001356 | EXP vs NON; U = 857.000000, p = 0.001793<br>EXP vs OBS; U = 2115.000000, p = 0.221200<br>OBS vs NON; U = 1424.000000, p = 0.040421 |
| main effect AH I (group) | H = 28.981570, p = 0.000001 | EXP vs NON; U = 729.000000, p = 0.000001<br>EXP vs OBS; U = 2030.000000, p = 0.001238<br>OBS vs NON; U = 1420.000000, p = 0.038272 |
| main effect POA I (group) | H = 2.533173, p = 0.281792 | n.s. |
| main effect MeA I (group) | H = 35.016884, p = 0.000000 | EXP vs NON; U = 595.000000, p = 0.000000<br>EXP vs OBS; U = 2233.000000, p = 0.014853<br>OBS vs NON; U = 1173.000000, p = 0.000665 |
| main effect PA I (group) | H = 2.731673, p = 0.255167 | n.s. |
| main effect BNST I (group) | H = 16.967636, p = 0.000207 | EXP vs NON; U = 895.000000, p = 0.000339<br>EXP vs OBS; U = 2398.000000, p = 0.270065 |

| | | | | | |
|---|---|---|---|---|---|
| | | | | | OBS vs NON; U = 1302.000000, p = 0.006530 |
| | | | main effect PAG I (group) | H = 15.023507, p = 0.000547 | EXP vs NON; U = 859.000000, p = 0.001878<br>EXP vs OBS; U = 1734.000000, p = 0.002757<br>OBS vs NON; U = 1860.000000, p = 1.000000 |
| | | | main effect vLS I (group) | H = 9.283485, p = 0.009641 | EXP vs NON; U = 1127.000000, p = 0.010630<br>EXP vs OBS; U = 2517.000000, p = 0.215303<br>OBS vs NON; U = 1590.000000, p = 0.289087 |
| | | | main effect PrL I (group) | H = 3.404978, p = 0.182229 | n.s. |
| | | | main effect Nac DA (group) | H = 8.942867, p = 0.011431 | EXP vs NON; U = 1257.000000, p = 0.465333<br>EXP vs OBS; U = 1949.000000, p = 0.008435<br>OBS vs NON; U = 2176.000000, p = 0.707350 |
| Fig 4c (bottom) & | *Cosine similarity distributions* | | *Two-sample Kolmogorov Smirnov tests* | | *Benjamini-Hochberg FDR adjustments* |
| Fig 4d (bottom) | *between EXP/OBS activity* | OBS vs. EXP = 84, OBS vs. NON = 42 | VMHvl E | KS = 0.285714, p = 0.018921 | p = 0.017344 |
| | *maps and OBS/NON activity maps* | OBS vs. EXP = 63, OBS vs. NON = 42 | PMv E | KS = 0.460317, p = 0.000027 | p = 0.000136 |
| | | OBS vs. EXP = 70, OBS vs. NON = 42 | AH E | KS = 0.333333, p = 0.004495 | p = 0.004945 |
| | | OBS vs. EXP = 84, OBS vs. NON = 42 | POA E | KS = 0.166667, p = 0.406155 | p = 0.230593 |
| | | OBS vs. EXP = 77, OBS vs. NON = 42 | MeA E | KS = 0.406926, p = 0.000150 | p = 0.000286 |
| | | OBS vs. EXP = 70, OBS vs. NON = 42 | PA E | KS = 0.400000, p = 0.000298 | p = 0.000469 |
| | | OBS vs. EXP = 70, OBS vs. NON = 42 | BNST E | KS = 0.138095, p = 0.653965 | p = 0.312766 |
| | | OBS vs. EXP = 63, OBS vs. NON = 42 | PAG E | KS = 0.253968, p = 0.068414 | p = 0.057889 |
| | | OBS vs. EXP = 84, OBS vs. NON = 42 | vLS E | KS = 0.214286, p = 0.146003 | p = 0.100377 |
| | | OBS vs. EXP = 77, OBS vs. NON = 42 | PrL E | KS = 0.145022, p = 0.563322 | p = 0.281661 |
| | | OBS vs. EXP = 70, OBS vs. NON = 42 | LHb E | KS = 0.347619, p = 0.002629 | p = 0.003213 |
| | | OBS vs. EXP = 84, OBS vs. NON = 42 | VMHvl I | KS = 0.404762, p = 0.000156 | p = 0.000286 |
| | | OBS vs. EXP = 63, OBS vs. NON = 42 | PMv I | KS = 0.452381, p = 0.000040 | p = 0.000136 |
| | | OBS vs. EXP = 70, OBS vs. NON = 42 | AH I | KS = 0.171429, p = 0.385103 | p = 0.230593 |

| | | | | | |
|---|---|---|---|---|---|
| | | OBS vs. EXP = 84, OBS vs. NON = 42 | POA I | KS = 0.238095, p = 0.079028 | p = 0.062093 |
| | | OBS vs. EXP = 77, OBS vs. NON = 42 | MeA I | KS = 0.445887, p = 0.000021 | p = 0.000136 |
| | | OBS vs. EXP = 70, OBS vs. NON = 42 | PA I | KS = 0.438095, p = 0.000049 | p = 0.000136 |
| | | OBS vs. EXP = 70, OBS vs. NON = 42 | BNST I | KS = 0.166667, p = 0.419261 | p = 0.230593 |
| | | OBS vs. EXP = 63, OBS vs. NON = 42 | PAG I | KS = 0.198413, p = 0.251457 | p = 0.162708 |
| | | OBS vs. EXP = 84, OBS vs. NON = 42 | vLS I | KS = 0.357143, p = 0.001320 | p = 0.001815 |
| | | OBS vs. EXP = 77, OBS vs. NON = 42 | PrL I | KS = 0.151515, p = 0.507567 | p = 0.265868 |
| | | OBS vs. EXP = 84, OBS vs. NON = 42 | NAc DA | KS = 0.297619, p = 0.012683 | p = 0.012683 |
| Fig 4e | *Mean similarity* | | *Independence t-test (t) or Mann-Whitney U test (U)* | | *Benjamini-Hochberg FDR adjustments* |
| | *between EXP/OBS activity maps and OBS/NON activity maps* | OBS vs. EXP = 84, OBS vs. NON = 42 | VMHvl E | U = 1136.000000, p = 0.001165 | p = 0.001289 |
| | | OBS vs. EXP = 63, OBS vs. NON = 42 | PMv E | U = 623.000000, p = 0.000005 | p = 0.000012 |
| | | OBS vs. EXP = 70, OBS vs. NON = 42 | AH E | U = 934.000000, p = 0.001289 | p = 0.001289 |
| | | OBS vs. EXP = 84, OBS vs. NON = 42 | POA E | U = 1621.000000, p = 0.460804 | p = 0.242528 |
| | | OBS vs. EXP = 77, OBS vs. NON = 42 | MeA E | t = -5.013996, p = 0.000002 | p = 0.000012 |
| | | OBS vs. EXP = 70, OBS vs. NON = 42 | PA E | t = -4.569104, p = 0.000013 | p = 0.000026 |
| | | OBS vs. EXP = 70, OBS vs. NON = 42 | BNST E | U = 1407.000000, p = 0.707194 | p = 0.353597 |
| | | OBS vs. EXP = 63, OBS vs. NON = 42 | PAG E | U = 1591.000000, p = 0.080124 | p = 0.053416 |
| | | OBS vs. EXP = 84, OBS vs. NON = 42 | vLS E | U = 1522.000000, p = 0.211371 | p = 0.132107 |
| | | OBS vs. EXP = 77, OBS vs. NON = 42 | PrL E | U = 1800.000000, p = 0.310187 | p = 0.182463 |
| | | OBS vs. EXP = 70, OBS vs. NON = 42 | LHb E | U = 1963.000000, p = 0.003066 | p = 0.002555 |
| | | OBS vs. EXP = 84, OBS vs. NON = 42 | VMHvl I | U = 972.000000, p = 0.000042 | p = 0.000058 |
| | | OBS vs. EXP = 63, OBS vs. NON = 42 | PMv I | U = 624.000000, p = 0.000005 | p = 0.000012 |

| | | | | | |
|---|---|---|---|---|---|
| | | OBS vs. EXP = 70, OBS vs. NON = 42 | AH I | U = 1425.000000, p = 0.789125 | p = 0.358693 |
| | | OBS vs. EXP = 84, OBS vs. NON = 42 | POA I | U = 1219.000000, p = 0.004812 | p = 0.003702 |
| | | OBS vs. EXP = 77, OBS vs. NON = 42 | MeA I | U = 883.000000, p = 0.000045 | p = 0.000058 |
| | | OBS vs. EXP = 70, OBS vs. NON = 42 | PA I | U = 700.000000, p = 0.000004 | p = 0.000012 |
| | | OBS vs. EXP = 70, OBS vs. NON = 42 | BNST I | U = 1326.000000, p = 0.387877 | p = 0.215487 |
| | | OBS vs. EXP = 63, OBS vs. NON = 42 | PAG I | U = 1319.000000, p = 0.981731 | p = 0.426840 |
| | | OBS vs. EXP = 84, OBS vs. NON = 42 | vLS I | t = -4.222145, p = 0.000046 | p = 0.000058 |
| | | OBS vs. EXP = 77, OBS vs. NON = 42 | PrL I | U = 1562.000000, p = 0.761845 | p = 0.358693 |
| | | OBS vs. EXP = 84, OBS vs. NON = 42 | NAc DA | U = 1187.000000, p = 0.002848 | p = 0.002555 |
| Fig 4f | *Cosine similarity distributions between aggregated EXP/OBS activity maps and aggregated OBS/NON activity maps* | | *Two-sample Kolmogorov Smirnov tests* | | |
| | | OBS vs. EXP = 812, OBS vs. NON = 462 | Excitatory distributions | KS = 0.1296, p = 8.841e-05 | N/A |
| | | OBS vs. EXP = 812, OBS vs. NON = 462 | Inhibitory distributions | KS = 0.2106, p = 6.3076e-12 | N/A |
| Fig 5a | *Decoding results* | EXP = 12, OBS = 7, NON = 6 | *Independence t-tests (t)* | | *Benjamini-Hochberg FDR adjustments* |
| | | | EXP: Cross-validation within-day | EXP vs NON; t = 19.416186, p = 0.000000 | p = 0.000000 |
| | | | EXP: Predicting on held-out signals from late training | EXP vs NON; t = 13.405998, p = 0.000000 | p = 0.000000 |
| | | | EXP: Predicting on held-out signals from hard fight | EXP vs NON; t = 10.732263, p = 0.000000 | p = 0.000000 |
| | | | OBS: Cross-validation within-day | OBS vs NON; t = 0.731368, p = 0.476618 | p = 0.612795 |
| | | | OBS: Predicting on held-out signals from late training | OBS vs NON; t = 0.071565, p = 0.944127 | p = 0.944127 |
| | | | OBS: Predicting on held-out signals from hard fight | OBS vs NON; t = 0.396371, p = 0.697807 | p = 0.785033 |
| | | | EXP: Decoding validation | EXP vs shuffle; t = 5.758901, p = 0.000009 | p = 0.000013 |

| | | | | | |
|---|---|---|---|---|---|
| | | | during hard fight | | |
| | | | OBS: Decoding validation during hard fight | OBS vs shuffle; t = 8.528477, p = 0.000002 | p = 0.000004 |
| | | | NON: Decoding validation during hard fight | NON vs shuffle; t = 8.718138, p = 0.000006 | p = 0.000010 |
| Fig 5b | *Time effect on predicting HF signals* | EXP = 12 | *Linear mixed effects model* | | |
| | | | EXP main effect (day) | t = 2.3449, p = 0.01903 | N/.A |
| Fig 5c | *Importance testing of* | EXP = 12, OBS = 7, NON = 6 | *Related samples t-tests (t)* | | *Benjamini-Hochberg FDR adjustments* |
| | *excitatory and inhibitory circuits* | | Training Period EXP: E vs. I | t = 4.620738, p = 0.000739 | |
| | | | Hard fight EXP: E vs I | t = 4.852835, p = 0.000508 | p = 0.001525 |
| | | | Hard fight OBS: E vs I | t = 0.201474, p = 0.846984 | p = 0.846985 |
| | | | Hard fight NON: E vs I | t = -1.003730, p = 0.361581 | p = 0.542373 |
| Fig 5d | *Time effect on E vs. I importance* | EXP = 12 | *Linear mixed effects model* | | *Bonferroni adjustments* |
| | | | EXP E main effect (day) | t = 2.462728, p = 0.013788 | p = 0.0275769 |
| | | | EXP I main effect (day) | t = 1.121107, p = 0.262242 | p = 0.52448426 |
| Fig 5f (training period, left) | *Excitatory-inhibitory decoupling* | | *Two-sample Kolmogorov Smirnov tests* | | *Benjamini-Hochberg FDR adjustments* |
| | *of neural activity* | EXP early N = 270, EXP late N = 270 | EXP training period VMHvl | KS = 0.162963, p = 0.001510 | p = 0.003019 |
| | *(training period)* | EXP early N = 210, EXP late N = 210 | EXP training period PMv | KS = 0.171429, p = 0.004111 | p = 0.006167 |
| | | EXP early N = 270, EXP late N = 270 | EXP training period AH | KS = 0.125926, p = 0.027550 | p = 0.027550 |
| | | EXP early N = 240, EXP late N = 240 | EXP training period POA | KS = 0.066667, p = 0.661473 | p = 0.396884 |
| | | EXP early N = 270, EXP late N = 270 | EXP training period MeA | KS = 0.181481, p = 0.000266 | p = 0.000798 |
| | | EXP early N = 210, EXP late N = 210 | EXP training period PA | KS = 0.147619, p = 0.020470 | p = 0.024564 |
| | | EXP early N = 240, EXP late N = 240 | EXP training period BNST | KS = 0.237500, p = 0.000002 | p = 0.000014 |
| | | OBS early N = 210, OBS late N = 210 | OBS training period VMHvl | KS = 0.166667, p = 0.005780 | p = 0.010114 |
| | | OBS early N = 210, OBS late N = 210 | OBS training period PMv | KS = 0.214286, p = 0.000123 | p = 0.000863 |
| | | OBS early N = 210, OBS late N = 210 | OBS training period AH | KS = 0.176190, p = 0.002896 | p = 0.010114 |

| | | | | | |
|---|---|---|---|---|---|
| | | OBS early N = 210, OBS late N = 210 | OBS training period POA | KS = 0.114286, p = 0.128812 | p = 0.100187 |
| | | OBS early N = 210, OBS late N = 210 | OBS training period MeA | KS = 0.133333, p = 0.047721 | p = 0.062062 |
| | | OBS early N = 210, OBS late N = 210 | OBS training period PA | KS = 0.071429, p = 0.658879 | p = 0.461215 |
| | | OBS early N = 210, OBS late N = 210 | OBS training period BNST | KS = 0.052381, p = 0.936400 | p = 0.595891 |
| | | NON early N = 180, NON late N = 180 | NON training period VMHvl | KS = 0.100000, p = 0.329810 | p = 0.293164 |
| | | NON early N = 179, NON late N = 180 | NON training period PMv | KS = 0.109559, p = 0.203870 | p = 0.203870 |
| | | NON early N = 180, NON late N = 180 | NON training period AH | KS = 0.138889, p = 0.061999 | p = 0.120974 |
| | | NON early N = 180, NON late N = 180 | NON training period POA | KS = 0.127778, p = 0.105852 | p = 0.120974 |
| | | NON early N = 179, NON late N = 180 | NON training period MeA | KS = 0.055525, p = 0.920161 | p = 0.669208 |
| | | NON early N = 179, NON late N = 180 | NON training period PA | KS = 0.210646, p = 0.000499 | p = 0.001994 |
| | | NON early N = 179, NON late N = 180 | NON training period BNST | KS = 0.126816, p = 0.094130 | p = 0.120974 |
| Fig 5f (hard fight, right) | *Excitatory-inhibitory decoupling* | (first 3 attacks) | *Two-sample Kolmogorov Smirnov tests* | *Pairwise comparisons* | *Benjamini-Hochberg FDR adjustments* |
| | *of neural activity* | AGG = 26, OBS = 21, NON = 17 | hard fight VMHvl | AGG vs NON; KS = 0.219457, p = 0.621114<br>OBS vs AGG; KS = 0.232601, p = 0.472864<br>OBS vs NON; KS = 0.204482, p = 0.738195 | AGG vs NON; p = 0.738195<br>OBS vs AGG; p = 0.738195<br>OBS vs NON; p = 0.738195 |
| | *(hard fight)* | AGG = 20, OBS = 21, NON = 17 | hard fight PMv | AGG vs NON; KS = 0.564706, p = 0.003123<br>OBS vs AGG; KS = 0.316667, p = 0.201894<br>OBS vs NON; KS = 0.445378, p = 0.031644 | AGG vs NON; p = 0.003123<br>OBS vs AGG; p = 0.067298<br>OBS vs NON; p = 0.015822 |
| | | AGG = 26, OBS = 21, NON = 17 | hard fight AH | AGG vs NON; KS = 0.608597, p = 0.000466<br>OBS vs AGG; KS = 0.452381, p = 0.011283<br>OBS vs NON; KS = 0.327731, p = 0.201160 | AGG vs NON; p = 0.000466<br>OBS vs AGG; p = 0.005641<br>OBS vs NON; p = 0.067053 |
| | | AGG = 22, OBS = 21, NON = 17 | hard fight POA | AGG vs NON; KS = 0.307487, p = 0.255978<br>OBS vs AGG; KS = 0.354978, p = 0.091421<br>OBS vs NON; KS = 0.243697, p = 0.532222 | AGG vs NON; p = 0.383968<br>OBS vs AGG; p = 0.274262<br>OBS vs NON; p = 0.532222 |
| | | AGG = 26, OBS = 21, NON = 17 | hard fight MeA | AGG vs NON; KS = 0.395928, p = 0.060183<br>OBS vs AGG; KS = 0.300366, p = 0.196117<br>OBS vs NON; KS = 0.291317, p = 0.321330 | AGG vs NON; p = 0.180550<br>OBS vs AGG; p = 0.294176<br>OBS vs NON; p = 0.321330 |

| | | | | | |
|---|---|---|---|---|---|
| | | AGG = 20, OBS = 21, NON = 17 | hard fight PA | AGG vs NON; KS = 0.447059, p = 0.034040 OBS vs AGG; KS = 0.464286, p = 0.015905 OBS vs NON; KS = 0.254902, p = 0.480718 | AGG vs NON; p = 0.034040 OBS vs AGG; p = 0.031810 OBS vs NON; p = 0.320478 |
| | | AGG = 24, OBS = 21, NON = 17 | hard fight BNST | AGG vs NON; KS = 0.549020, p = 0.002682 OBS vs AGG; KS = 0.458333, p = 0.011596 OBS vs NON; KS = 0.190476, p = 0.803795 | AGG vs NON; p = 0.002682 OBS vs AGG; p = 0.005798 OBS vs NON; p = 0.267932 |
| Fig 5h (training period, left) | *Mapping changes in importance* | AGG early N = 210, AGG late N = 210 | *Two-sample Kolmogorov Smirnov tests* | | *Benjamini-Hochberg FDR adjustments* |
| | *(delta r-squared) in predictors* | OBS early N = 210, OBS late N = 210 | EXP training period PMv E | KS = 0.162963, p = 0.971729 | p = 0.566842 |
| | *encoding VMHvl E during attacks* | NON early N = 179, NON late N = 180 | EXP training period AH E | KS = 0.171429, p = 0.015141 | p = 0.021197 |
| | *(training period)* | | EXP training period POA E | KS = 0.125926, p = 0.424194 | p = 0.403854 |
| | | | EXP training period lPAG E | KS = 0.066667, p = 0.015141 | p = 0.021197 |
| | | | EXP training period MeA E | KS = 0.181481, p = 0.297256 | p = 0.346798 |
| | | | EXP training period PA E | KS = 0.147619, p = 0.576934 | p = 0.403854 |
| | | | EXP training period VMHvl I | KS = 0.237500, p = 0.002896 | p = 0.006758 |
| | | | EXP training period PMv I | KS = 0.061905, p = 0.002896 | p = 0.006758 |
| | | | EXP training period AH I | KS = 0.077778, p = 0.576934 | p = 0.403854 |
| | | | EXP training period POA I | KS = 0.118519, p = 0.002020 | p = 0.006758 |
| | | | EXP training period MeA I | KS = 0.085714, p = 0.497937 | p = 0.403854 |
| | | | EXP training period PA I | KS = 0.047619, p = 0.740398 | p = 0.471162 |
| | | | OBS training period PMv E | KS = 0.166667, p = 0.244848 | p = 0.349782 |
| | | | OBS training period AH E | KS = 0.214286, p = 0.002896 | p = 0.028898 |
| | | | OBS training period POA E | KS = 0.176190, p = 0.036340 | p = 0.072680 |
| | | | OBS training period lPAG E | KS = 0.114286, p = 0.297256 | p = 0.371570 |
| | | | OBS training period MeA E | KS = 0.133333, p = 0.936400 | p = 0.780333 |
| | | | OBS training period PA E | KS = 0.071429, p = 0.817093 | p = 0.742812 |
| | | | OBS training period VMHvl I | KS = 0.052381, p = 0.036340 | p = 0.072680 |
| | | | OBS training period PMv I | KS = 0.114286, p = 0.005780 | p = 0.028898 |
| | | | OBS training period AH I | KS = 0.166667, p = 0.576934 | p = 0.641038 |
| | | | OBS training period POA I | KS = 0.128571, p = 0.036340 | p = 0.072680 |
| | | | OBS training period MeA I | KS = 0.128571, p = 0.740398 | p = 0.740398 |
| | | | OBS training period PA I | KS = 0.100000, p = 0.199630 | p = 0.332717 |
| | | | NON training period PMv E | KS = 0.100000, p = 0.163024 | p = 0.298877 |

| | | | | | |
|---|---|---|---|---|---|
| | | | NON training period AH E | KS = 0.109559, p = 0.220208 | p = 0.339877 |
| | | | NON training period POA E | KS = 0.138889, p = 0.262482 | p = 0.339877 |
| | | | NON training period lPAG E | KS = 0.127778, p = 0.003193 | p = 0.035128 |
| | | | NON training period MeA E | KS = 0.055525, p = 0.142550 | p = 0.298877 |
| | | | NON training period PA E | KS = 0.210646, p = 0.651917 | p = 0.597590 |
| | | | NON training period VMHvl I | KS = 0.126816, p = 0.029471 | p = 0.108062 |
| | | | NON training period PMv I | KS = 0.133333, p = 0.025792 | p = 0.108062 |
| | | | NON training period AH I | KS = 0.255556, p = 0.278082 | p = 0.339877 |
| | | | NON training period POA I | KS = 0.196120, p = 0.118141 | p = 0.298877 |
| | | | NON training period MeA I | KS = 0.083333, p = 0.540250 | p = 0.540250 |
| | | | NON training period PA I | KS = 0.114991, p = 0.415839 | p = 0.457423 |
| Fig 5h (hard fight, right) | *Comparing importance* | AGG = 61, OBS = 84, NON = 46 | *Two-sample Kolmogorov Smirnov tests* | | *Benjamini-Hochberg FDR adjustments* |
| | *of VMHvl predictors* | | hard fight PMv E | AGG vs NON KS = 0.093015, p = 0.955290; OBS vs NON KS = 0.091615, p = 0.938923 | AGG p = 0.955290, OBS p = 0.955290 |
| | *between groups* | | hard fight AH E | AGG vs NON KS = 0.140413, p = 0.611161; OBS vs NON KS = 0.187888, p = 0.210096 | AGG p = 0.611161, OBS p = 0.420192 |
| | *during the hard fight* | | hard fight POA E | AGG vs NON KS = 0.100855, p = 0.918293; OBS vs NON KS = 0.124224, p = 0.690587 | AGG p = 0.918293, OBS p = 0.918293 |
| | *(attack)* | | hard fight lPAG E | AGG vs NON KS = 0.368496, p = 0.001012; OBS vs NON KS = 0.138716, p = 0.557299 | AGG p = 0.001012, OBS p = 0.278649 |
| | | | hard fight MeA E | AGG vs NON KS = 0.113685, p = 0.834318; OBS vs NON KS = 0.112319, p = 0.796390 | AGG p = 0.834318, OBS p = 0.834318 |
| | | | hard fight PA E | AGG vs NON KS = 0.132929, p = 0.680790; OBS vs NON KS = 0.217909, p = 0.099181 | AGG p = 0.680790, OBS p = 0.198361 |
| | | | hard fight VMHvl I | AGG vs NON KS = 0.187812, p = 0.269649; OBS vs NON KS = 0.107143, p = 0.838777 | AGG p = 0.539298, OBS p = 0.838777 |
| | | | hard fight PMv I | AGG vs NON KS = 0.308268, p = 0.009996; OBS vs NON KS = 0.266046, p = 0.023365 | AGG p = 0.019992, OBS p = 0.023365 |
| | | | hard fight AH I | AGG vs NON KS = 0.161083, p = 0.443871; OBS vs NON KS = 0.211180, p = 0.118469 | AGG p = 0.443871, OBS p = 0.236939 |
| | | | hard fight POA I | AGG vs NON KS = 0.191732, p = 0.251294; OBS vs NON KS = 0.095756, p = 0.916935 | AGG p = 0.502587, OBS p = 0.916935 |
| | | | hard fight MeA I | AGG vs NON KS = 0.301853, p = 0.012990; | AGG p = 0.012990, OBS p = 0.035126 |

| | | | | | |
|---|---|---|---|---|---|
| | | | hard fight PA I | OBS vs NON KS = 0.230331, p = 0.070252 AGG vs NON KS = 0.166073, p = 0.408144; OBS vs NON KS = 0.213768, p = 0.110678 | AGG p = 0.408144, OBS p = 0.221356 |
| Fig 5j | *Time effect on importance of predictors on VMHvl activity during social behavior* | AGG = 7, OBS = 7, NON = 6 | *Linear mixed effects model* | | *Bonferroni adjustments* |
| | | | EXP main effect (day) PMv E | t = -0.858494, p = 0.390620 | p = 1.000000 |
| | | | EXP main effect (day) AH E | t = 1.315282, p = 0.730652 | p = 1.000000 |
| | | | EXP main effect (day) POA E | t = 0.344259, p = 0.971851 | p = 1.000000 |
| | | | EXP main effect (day) lPAG E | t = -2.601253, p = 0.138874 | p = 1.000000 |
| | | | EXP main effect (day) MeA E | t = 0.035287, p = 0.039782 | p = 0.477389 |
| | | | EXP main effect (day) PA E | t = -2.963760, p = 0.047519 | p = 0.570231 |
| | | | EXP main effect (day) VMHvl I | t = 1.479999, p = 0.000023 | p = 0.000279 |
| | | | EXP main effect (day) PMv I | t = 0.910542, p = 0.004081 | p = 0.048970 |
| | | | EXP main effect (day) AH I | t = 2.056001, p = 0.071167 | p = 0.854006 |
| | | | EXP main effect (day) POA I | t = -0.292787, p = 0.442662 | p = 1.000000 |
| | | | EXP main effect (day) MeA I | t = 1.981643, p = 0.020657 | p = 0.247887 |
| | | | EXP main effect (day) PA I | t = -0.747312, p = 0.862241 | p = 1.000000 |
| | | | OBS main effect (day) PMv E | t = -0.858494, p = 0.188415 | p = 1.000000 |
| | | | OBS main effect (day) AH E | t = 1.315282, p = 0.009288 | p = 0.111461 |
| | | | OBS main effect (day) POA E | t = 0.344259, p = 0.003039 | p = 0.036469 |
| | | | OBS main effect (day) lPAG E | t = -2.601253, p = 0.362537 | p = 1.000000 |
| | | | OBS main effect (day) MeA E | t = 0.035287, p = 0.769685 | p = 1.000000 |
| | | | OBS main effect (day) PA E | t = -2.963760, p = 0.454875 | p = 1.000000 |
| | | | OBS main effect (day) VMHvl I | t = 1.479999, p = 0.339835 | p = 1.000000 |
| | | | OBS main effect (day) PMv I | t = 0.910542, p = 0.001924 | p = 0.023083 |

| | | | | | |
|---|---|---|---|---|---|
| | | | OBS main effect (day) AH I | t = 2.056001, p = 0.471384 | p = 1.000000 |
| | | | OBS main effect (day) POA I | t = -0.292787, p = 0.234506 | p = 1.000000 |
| | | | OBS main effect (day) MeA I | t = 1.981643, p = 0.223566 | p = 1.000000 |
| | | | OBS main effect (day) PA I | t = -0.747312, p = 0.905915 | p = 1.000000 |
| Fig 5k | *Comparing importance* | AGG = 143, OBS = 137, NON = 98 | *Two-sample Kolmogorov Smirnov tests* | | *Benjamini-Hochberg FDR adjustments* |
| | *of VMHvl predictors* | | hard fight PMv E | AGG vs NON KS = 0.127301, p = 0.272346; OBS vs NON KS = 0.111872, p = 0.430359 | AGG vs NON p = 0.430359; OBS vs NON p = 0.430359 |
| | *between groups* | | hard fight AH E | AGG vs NON KS = 0.187384, p = 0.028535; OBS vs NON KS = 0.218159, p = 0.007028 | AGG vs NON p = 0.028535; OBS vs NON p = 0.014056 |
| | *during the hard fight* | | hard fight POA E | AGG vs NON KS = 0.207649, p = 0.010903; OBS vs NON KS = 0.183599, p = 0.035701 | AGG vs NON p = 0.021805; OBS vs NON p = 0.035701 |
| | *(attack)* | | hard fight lPAG E | AGG vs NON KS = 0.224989, p = 0.004469; OBS vs NON KS = 0.092731, p = 0.664484 | AGG vs NON p = 0.895632; OBS vs NON p = 0.090755 |
| | | | hard fight MeA E | AGG vs NON KS = 0.071928, p = 0.895632; OBS vs NON KS = 0.178460, p = 0.045377 | AGG vs NON p = 0.704936; OBS vs NON p = 0.704936 |
| | | | hard fight PA E | AGG vs NON KS = 0.113244, p = 0.406283; OBS vs NON KS = 0.089602, p = 0.704936 | AGG vs NON p = 0.008939; OBS vs NON p = 0.664484 |
| | | | hard fight VMHvl I | AGG vs NON KS = 0.204438, p = 0.012772; OBS vs NON KS = 0.119246, p = 0.353048 | AGG vs NON p = 0.025544; OBS vs NON p = 0.353048 |
| | | | hard fight PMv I | AGG vs NON KS = 0.258813, p = 0.000628; OBS vs NON KS = 0.302771, p = 0.000038 | AGG vs NON p = 0.000628; OBS vs NON p = 0.000076 |
| | | | hard fight AH I | AGG vs NON KS = 0.197945, p = 0.017536; OBS vs NON KS = 0.245643, p = 0.001601 | AGG vs NON p = 0.017536; OBS vs NON p = 0.003203 |
| | | | hard fight POA I | AGG vs NON KS = 0.111817, p = 0.421587; OBS vs NON KS = 0.154178, p = 0.115014 | AGG vs NON p = 0.421587; OBS vs NON p = 0.230028 |
| | | | hard fight MeA I | AGG vs NON KS = 0.259241, p = 0.000606; OBS vs NON KS = 0.216893, p = 0.007405 | AGG vs NON p = 0.001213; OBS vs NON p = 0.007405 |
| | | | hard fight PA I | AGG vs NON KS = 0.177894, p = 0.043144; OBS vs NON KS = 0.114107, p = 0.408993 | AGG vs NON p = 0.086288; OBS vs NON p = 0.408993 |
| Fig E1a | *Comparing attack time and persistence distributions in early vs late* | early = N = 60 | *two-sample Kolmogorov Smirnov test* | | |
| | | late N = 60 | attack time | KS = 0.116667, p = 0.813281 | N/A |

| | | | | | |
|---|---|---|---|---|---|
| | | | attack persistence | KS= 0.200000, p = 0.182057 | |
| Fig E1b | *Comparing attack occupancy in early, late and HF* | N = 30 | Kruskal-Wallis H-test | H = 2.112988, p = 0.347673 | n.s. |
| Fig E1d | *Comparing behavior features during hard fight* | EXP = 10 | *One-way ANOVA tests* | | post hoc independent samples t-test |
| | | OBS = 7 | resident-intruder centroid distance | F = 8.165357, p = 0.002556 | AGG vs NON; t = -4.009824, p = 0.001291<br>AGG vs OBS; t = -1.497838, p = 0.051641<br>OBS vs NON; t = -2.838624, p = 0.008061 |
| | | NON = 6 | resident head-to-intruder tailbase distance | F = 9.909086, p = 0.001022 | AGG vs NON; t = -4.262371, p = 0.000789<br>AGG vs OBS; t = -1.983195, p = 0.021988<br>OBS vs NON; t = -2.812913, p = 0.008440 |
| | | | head-head distance | F = 6.504821, p = 0.006666 | AGG vs NON; t = -3.729609, p = 0.004484<br>AGG vs OBS; t = -1.425564, p = 0.116314<br>OBS vs NON; t = -2.282613, p = 0.043337 |
| | | | Intersection over union | F = 8.304569, p = 0.002368 | AGG vs NON; t = 3.983126, p = 0.001360<br>AGG vs OBS; t = 1.433286, p = 0.057430<br>OBS vs NON; t = 3.225148, p = 0.004042 |
| | | | resident pose | F = 23.994276, p = 0.000005 | AGG vs NON; t = 5.836082, p = 0.000130<br>AGG vs OBS; t = 4.771813, p = 0.000371<br>OBS vs NON; t = 2.265556, p = 0.044655 |
| | | | intruder orientation toward resident | F = 5.560739, p = 0.012014 | AGG vs NON; t = 3.619067, p = 0.005582<br>AGG vs OBS; t = 1.960943, p = 0.068725<br>OBS vs NON; t = 1.054645, p = 0.209468 |
| Fig E1e | *Group differences in cluster persistence* | | *one-way ANOVA (F) or Kruskal-Wallis (H) test* | | post hoc independent samples t-tests or Mann Whitney U tests |
| | | | main effect cluster 1 (group) | F = 1.428307, p = 0.263136 | |
| | | | main effect cluster 2 (group) | H = 0.754384, p = 0.685784 | |
| | | | main effect cluster 3 (group) | F = 0.319547, p = 0.730119 | |
| | | | main effect cluster 4 (group) | F = 2.769416, p = 0.086754 | |
| | | | main effect cluster 5 (group) | F = 2.745553, p = 0.088392 | |
| | | | main effect cluster 6 (group) | F = 5.112662, p = 0.016091 | AGG vs NON; t = -3.162652, p = 0.013831<br>AGG vs OBS; t = -1.372171, p = 0.126789<br>OBS vs NON; t = -1.772116, p = 0.104036 |

| | | | | | |
|---|---|---|---|---|---|
| | | | main effect cluster 7 (group) | H = 1.195031, p = 0.550177 | H = 1.619451, p = 0.444980 |
| | | | main effect cluster 8 (group) | H = 1.619451, p = 0.444980 | |
| | | | main effect cluster 9 (group) | F = 1.444163, p = 0.259513 | |
| | | | main effect cluster 10 (group) | F = 1.444163, p = 0.259513 | AGG vs NON; t = -3.554188, p = 0.006349<br>AGG vs OBS; t = -1.847129, p = 0.084548<br>OBS vs NON; t = -0.768821, p = 0.305460 |
| | | | main effect cluster 11 (group) | F = 3.038728, p = 0.070412 | AGG vs NON; t = 2.565218, p = 0.066510<br>AGG vs OBS; t = -0.019827, p = 0.984443<br>OBS vs NON; t = 2.269579, p = 0.066510 |
| | | | main effect cluster 12 (group) | H = 4.142443, p = 0.126032 | |
| | | | main effect cluster 13 (group) | H = 5.751139, p = 0.056384 | |
| Fig E1f | Group ID decoding via persistence | N = 1,000 permutations<br>EXP = 10<br>OBS = 7<br>NON = 6 | *Permutation test on F1 score* | F1 = 0.37, p = 0.123 | N/A |
| Fig E1g | Similarities in persistence | OBS/EXP = 630<br>OBS/NON = 378 | *two-sample Kolmogorov Smirnov test* | KS = 0.219, p = 2.11566e-10 | N/A |
| Fig E3c | *E-I bias in the SBN* | | *Independence t-test (t) or Mann-Whitney U test (U)* | | *Benjamini-Hochberg FDR adjustments* |
| | | EXP = 12, NON = 6 | VMHvl (EXP) | AGG vs NON; t = 3.592358, p = 0.003279 | p = 0.029515 |
| | | EXP = 8, NON = 6 | PMv (EXP) | AGG vs NON; t = 3.250701, p = 0.007726 | p = 0.034765 |
| | | EXP = 10, NON = 6 | AH (EXP) | AGG vs NON; U = 36.000000, p = 0.327672 | p = 0.445910 |
| | | EXP = 11, NON = 6 | POA (EXP) | AGG vs NON; t = 1.023375, p = 0.326319 | p = 0.445910 |
| | | EXP = 11, NON = 6 | MeA (EXP) | AGG vs NON; t = -0.246774, p = 0.808937 | p = 0.766361 |
| | | EXP = 9, NON = 6 | PA (EXP) | AGG vs NON; t = -0.493170, p = 0.631590 | p = 0.668678 |
| | | EXP = 10, NON = 6 | BNST (EXP) | AGG vs NON; t = 0.739659, p = 0.473729 | p = 0.532945 |
| | | EXP = 8, NON = 6 | PAG (EXP) | AGG vs NON; t = 1.626096, p = 0.132209 | p = 0.297471 |
| | | EXP = 12, NON = 6 | vLS (EXP) | AGG vs NON; t = -0.925453, p = 0.371591 | p = 0.445910 |
| | | EXP = 11, NON = 6 | PrL (EXP) | AGG vs NON; t = 0.437856, p = 0.668678 | p = 0.668678 |
| | | EXP = 9, NON = 6 | LHb (EXP) | AGG vs NON; t = -5.718022, p = 0.000135 | p = 0.002422 |
| | | OBS = 7, NON = 6 | VMHvl (OBS) | OBS vs NON; t = 2.363476, p = 0.037582 | p = 0.112747 |

| | | | | | |
|---|---|---|---|---|---|
| | | OBS = 7, NON = 6 | PMv (OBS) | OBS vs NON; t = 3.469271, p = 0.005247 | p = 0.031480 |
| | | OBS = 7, NON = 6 | AH E(OBS) | OBS vs NON; U = 34.000000, p = 0.073427 | p = 0.188811 |
| | | OBS = 7, NON = 6 | POA E(OBS) | OBS vs NON; t = 2.629289, p = 0.023437 | p = 0.084373 |
| | | OBS = 7, NON = 6 | MeA E(OBS) | OBS vs NON; U = 13.000000, p = 0.294872 | p = 0.445910 |
| | | OBS = 7, NON = 6 | PA (OBS) | OBS vs NON; t = -0.074439, p = 0.941998 | p = 0.810190 |
| | | OBS = 7, NON = 6 | BNST (OBS) | OBS vs NON; t = 0.961871, p = 0.356782 | p = 0.445910 |
| | | OBS = 7, NON = 6 | PAG (OBS) | OBS vs NON; t = 1.115688, p = 0.288336 | p = 0.445910 |
| | | OBS = 7, NON = 6 | vLS (OBS) | OBS vs NON; U = 20.000000, p = 0.945221 | p = 0.810190 |
| | | OBS = 7, NON = 6 | PrL (OBS) | OBS vs NON; t = -0.974411, p = 0.350794 | p = 0.445910 |
| | | OBS = 7, NON = 6 | LHb (OBS) | OBS vs NON; t = 0.008509, p = 0.993363 | p = 0.812752 |
| Fig E4a-b | *Nonsocial versus social or aggression* | (late) | 1 sample t-test or Wilcoxon rank sum test | | *Benjamini-Hochberg FDR adjustments* |
| | *preferences* | EXP = 12 | VMHvl E (EXP) | t = 7.927278, p = 0.000007 | p = 0.000093 |
| | | EXP = 8 | PMv E (EXP) | W = 0.000000, p = 0.007812 | p = 0.010691 |
| | | EXP = 10 | AH E (EXP) | t = 3.878179, p = 0.003742 | p = 0.005405 |
| | | EXP = 11 | POA E (EXP) | t = 4.638440, p = 0.000924 | p = 0.002308 |
| | | EXP = 11 | MeA E (EXP) | W = 0.000000, p = 0.000977 | p = 0.002308 |
| | | EXP = 9 | PA E (EXP) | t = 6.782254, p = 0.000140 | p = 0.000730 |
| | | EXP = 10 | BNST E (EXP) | t = 6.977945, p = 0.000065 | p = 0.000421 |
| | | EXP = 8 | PAG E (EXP) | t = 4.345127, p = 0.003375 | p = 0.005178 |
| | | EXP = 12 | vLS E (EXP) | t = 3.943870, p = 0.002297 | p = 0.004265 |
| | | EXP = 11 | PrL E (EXP) | W = 0.000000, p = 0.000977 | p = 0.002308 |
| | | EXP = 9 | LHb E (EXP) | W = 17.000000, p = 0.570312 | p = 0.390214 |
| | | EXP = 12 | VMHvl I (EXP) | t = 8.452053, p = 0.000004 | p = 0.000093 |
| | | EXP = 8 | PMv I (EXP) | t = 4.384073, p = 0.003218 | p = 0.005178 |
| | | EXP = 10 | AH I (EXP) | t = 4.472883, p = 0.001548 | p = 0.003096 |
| | | EXP = 11 | POA I (EXP) | t = 4.650005, p = 0.000908 | p = 0.002308 |
| | | EXP = 11 | MeA I (EXP) | t = 4.405545, p = 0.001324 | p = 0.002868 |
| | | EXP = 9 | PA I (EXP) | t = 7.834095, p = 0.000051 | p = 0.000421 |
| | | EXP = 10 | BNST I (EXP) | t = 5.154219, p = 0.000600 | p = 0.002308 |
| | | EXP = 8 | PAG I (EXP) | t = 3.491094, p = 0.010114 | p = 0.013148 |
| | | EXP = 12 | vLS I (EXP) | t = 1.803262, p = 0.098777 | p = 0.111661 |
| | | EXP = 11 | PrL I (EXP) | W = 0.000000, p = 0.000977 | p = 0.002308 |
| | | EXP = 11 | NAc DA (EXP) | t = 2.105155, p = 0.061542 | p = 0.076194 |
| | | OBS = 7 | VMHvl E (OBS) | t = -0.329917, p = 0.752677 | p = 0.444763 |

| | | | |
|---|---|---|---|
| OBS = 7 | PMv E (OBS) | t = 1.966700, p = 0.096792 | p = 0.111661 |
| OBS = 7 | AH E(OBS) | t = -0.632939, p = 0.550120 | p = 0.390214 |
| OBS = 7 | POA E(OBS) | t = -0.927415, p = 0.389494 | p = 0.326673 |
| OBS = 7 | MeA E(OBS) | t = 1.219811, p = 0.268304 | p = 0.257292 |
| OBS = 7 | PA E (OBS) | W = 7.000000, p = 0.296875 | p = 0.257292 |
| OBS = 7 | BNST E (OBS) | t = -1.203054, p = 0.274268 | p = 0.257292 |
| OBS = 7 | PAG E (OBS) | t = -0.139932, p = 0.893293 | p = 0.516125 |
| OBS = 7 | vLS E (OBS) | t = -1.670810, p = 0.145794 | p = 0.157944 |
| OBS = 7 | PrL E (OBS) | t = -0.731103, p = 0.492268 | p = 0.365685 |
| OBS = 7 | LHb E (OBS) | W = 7.000000, p = 0.296875 | p = 0.257292 |
| OBS = 7 | VMHvl I (OBS) | t = -1.481800, p = 0.188904 | p = 0.196460 |
| OBS = 7 | PMv I (OBS) | t = -0.618988, p = 0.558678 | p = 0.390214 |
| OBS = 7 | AH I (OBS) | t = -0.387328, p = 0.711880 | p = 0.440688 |
| OBS = 7 | POA I (OBS) | t = -0.399194, p = 0.703571 | p = 0.440688 |
| OBS = 7 | MeA I (OBS) | t = -1.166525, p = 0.287666 | p = 0.257292 |
| OBS = 7 | PA I (OBS) | t = -0.449693, p = 0.668721 | p = 0.436213 |
| OBS = 7 | BNST I (OBS) | t = -0.837012, p = 0.434651 | p = 0.341802 |
| OBS = 7 | PAG I (OBS) | t = 0.353195, p = 0.736020 | p = 0.444763 |
| OBS = 7 | vLS I (OBS) | t = 0.446211, p = 0.671097 | p = 0.436213 |
| OBS = 7 | PrL I (OBS) | t = 0.883253, p = 0.411089 | p = 0.334010 |
| OBS = 7 | NAc DA (OBS) | t = -0.010459, p = 0.991994 | p = 0.560692 |
| (early) | | | |
| EXP = 12 | VMHvl E (EXP) | t = 13.067654, p = 0.000000 | p = 0.000001 |
| EXP = 8 | PMv E (EXP) | t = 10.265738, p = 0.000018 | p = 0.000060 |
| EXP = 10 | AH E (EXP) | t = 4.294263, p = 0.002007 | p = 0.001912 |
| EXP = 11 | POA E (EXP) | t = 4.916999, p = 0.000608 | p = 0.001013 |
| EXP = 11 | MeA E (EXP) | W = 0.000000, p = 0.000977 | p = 0.001289 |
| EXP = 9 | PA E (EXP) | t = 6.809680, p = 0.000136 | p = 0.000268 |
| EXP = 10 | BNST E (EXP) | t = 6.407252, p = 0.000124 | p = 0.000268 |
| EXP = 8 | PAG E (EXP) | t = 2.513813, p = 0.040172 | p = 0.028926 |
| EXP = 12 | vLS E (EXP) | t = 6.813888, p = 0.000029 | p = 0.000083 |
| EXP = 11 | PrL E (EXP) | t = 4.786701, p = 0.000738 | p = 0.001136 |
| EXP = 9 | LHb E (EXP) | t = -0.668134, p = 0.522853 | p = 0.237661 |
| EXP = 12 | VMHvl I (EXP) | t = 8.140774, p = 0.000006 | p = 0.000028 |
| EXP = 8 | PMv I (EXP) | t = 7.416744, p = 0.000147 | p = 0.000268 |

| | | | | | |
|---|---|---|---|---|---|
| | | EXP = 10 | AH I (EXP) | t = 3.169159, p = 0.011381 | p = 0.009503 |
| | | EXP = 11 | POA I (EXP) | t = 4.597469, p = 0.000984 | p = 0.001289 |
| | | EXP = 11 | MeA I (EXP) | t = 9.781107, p = 0.000002 | p = 0.000013 |
| | | EXP = 9 | PA I (EXP) | t = 17.330587, p = 0.000000 | p = 0.000001 |
| | | EXP = 10 | BNST I (EXP) | t = 9.137244, p = 0.000008 | p = 0.000030 |
| | | EXP = 8 | PAG I (EXP) | t = 5.178473, p = 0.001283 | p = 0.001426 |
| | | EXP = 12 | vLS I (EXP) | t = 1.822954, p = 0.095577 | p = 0.050304 |
| | | EXP = 11 | PrL I (EXP) | t = 4.527661, p = 0.001095 | p = 0.001289 |
| | | EXP = 11 | NAc DA (EXP) | t = 3.393856, p = 0.006841 | p = 0.006219 |
| | | OBS = 7 | VMHvl E (OBS) | t = 3.219139, p = 0.018157 | p = 0.013967 |
| | | OBS = 7 | PMv E (OBS) | t = 3.485585, p = 0.013053 | p = 0.010443 |
| | | OBS = 7 | AH E(OBS) | W = 2.000000, p = 0.046875 | p = 0.030778 |
| | | OBS = 7 | POA E(OBS) | t = 2.603056, p = 0.040496 | p = 0.028926 |
| | | OBS = 7 | MeA E(OBS) | t = -0.127036, p = 0.903063 | p = 0.401361 |
| | | OBS = 7 | PA E (OBS) | W = 13.000000, p = 0.937500 | p = 0.407609 |
| | | OBS = 7 | BNST E (OBS) | t = 5.283408, p = 0.001859 | p = 0.001912 |
| | | OBS = 7 | PAG E (OBS) | t = 1.232328, p = 0.263923 | p = 0.128743 |
| | | OBS = 7 | vLS E (OBS) | t = 2.481546, p = 0.047705 | p = 0.030778 |
| | | OBS = 7 | PrL E (OBS) | W = 2.000000, p = 0.046875 | p = 0.030778 |
| | | OBS = 7 | LHb E (OBS) | t = 2.279342, p = 0.062857 | p = 0.034591 |
| | | OBS = 7 | VMHvl I (OBS) | t = 3.597311, p = 0.011403 | p = 0.009503 |
| | | OBS = 7 | PMv I (OBS) | t = 2.266288, p = 0.063993 | p = 0.034591 |
| | | OBS = 7 | AH I (OBS) | t = 5.903424, p = 0.001050 | p = 0.001289 |
| | | OBS = 7 | POA I (OBS) | t = 1.252127, p = 0.257120 | p = 0.128560 |
| | | OBS = 7 | MeA I (OBS) | t = 2.326075, p = 0.058957 | p = 0.033689 |
| | | OBS = 7 | PA I (OBS) | t = 2.404318, p = 0.052982 | p = 0.032110 |
| | | OBS = 7 | BNST I (OBS) | t = 2.456151, p = 0.049377 | p = 0.030860 |
| | | OBS = 7 | PAG I (OBS) | t = 1.743578, p = 0.131856 | p = 0.067619 |
| | | OBS = 7 | vLS I (OBS) | W = 7.000000, p = 0.296875 | p = 0.138081 |
| | | OBS = 7 | PrL I (OBS) | t = 5.206490, p = 0.002002 | p = 0.001912 |
| | | OBS = 7 | NAc DA (OBS) | t = 1.156377, p = 0.291486 | p = 0.138081 |
| Fig E4c | z-scored DF/F during hard fight aggro | | One-way ANOVA tests or Kruskal Wallis H tests | | Post hoc independent samples t tests or Mann-Whitney U tests with Benjamini-Hochberg correction |
| | cluster | NON = 6, OBS = 7, EXP = 12 | VMHvl E | H = 5.484396, p = 0.064429 | n.s. |

| | | | | |
|---|---|---|---|---|
| | NON = 6, OBS = 7, EXP = 9 | PMv E | F = 3.438832, p = 0.054346 | n.s. |
| | NON = 6, OBS = 7, EXP = 10 | AH E | F = 2.829230, p = 0.082793 | n.s. |
| | NON = 6, OBS = 7, EXP = 11 | POA E | F = 3.375398, p = 0.053569 | n.s. |
| | NON = 6, OBS = 7, EXP = 11 | MeA E | H = 0.141818, p = 0.931547 | n.s. |
| | NON = 6, OBS = 7, EXP = 9 | PA E | F = 5.070919, p = 0.017189 | AGG vs NON; t = 2.861078, p = 0.026740<br>AGG vs OBS; t = 1.628087, p = 0.083863<br>OBS vs NON; t = 1.971041, p = 0.074399 |
| | NON = 6, OBS = 7, EXP = 10 | BNST E | F = 1.378370, p = 0.274916 | n.s. |
| | NON = 6, OBS = 7, EXP = 9 | PAG E | F = 0.120509, p = 0.887179 | n.s. |
| | NON = 6, OBS = 7, EXP = 12 | vLS E | F = 2.285984, p = 0.125311 | n.s. |
| | NON = 6, OBS = 7, EXP = 11 | PrL E | F = 1.077960, p = 0.358386 | n.s. |
| | NON = 6, OBS = 7, EXP = 9 | LHb E | H = 7.053757, p = 0.029397 | AGG vs NON; U = 7.000000, p = 0.061202<br>AGG vs OBS; U = 12.000000, p = 0.064128<br>OBS vs NON; U = 20.000000, p = 0.945221 |
| | NON = 6, OBS = 7, EXP = 12 | VMHvl I | F = 6.848225, p = 0.004873 | AGG vs NON; t = -2.637763, p = 0.008955<br>AGG vs OBS; t = -4.112621, p = 0.000727<br>OBS vs NON; t = 0.335294, p = 0.247904 |
| | NON = 6, OBS = 7, EXP = 9 | PMv I | F = 4.871800, p = 0.020370 | AGG vs NON; t = -2.420797, p = 0.016135<br>AGG vs OBS; t = 0.057428, p = 0.318359<br>OBS vs NON; t = -2.487839, p = 0.016135 |
| | NON = 6, OBS = 7, EXP = 10 | AH I | F = 8.539374, p = 0.002085 | AGG vs NON; t = -3.585808, p = 0.005963<br>AGG vs OBS; t = -2.009646, p = 0.041875<br>OBS vs NON; t = -2.258552, p = 0.041875 |
| | NON = 6, OBS = 7, EXP = 11 | POA I | F = 0.511588, p = 0.606825 | n.s. |
| | NON = 6, OBS = 7, EXP = 11 | MeA I | F = 8.122066, p = 0.002439 | AGG vs NON; t = -3.698569, p = 0.002145<br>AGG vs OBS; t = -2.517351, p = 0.011431<br>OBS vs NON; t = -1.741580, p = 0.036479 |
| | NON = 6, OBS = 7, EXP = 9 | PA I | H = 2.541878, p = 0.280568 | n.s. |
| | NON = 6, OBS = 7, EXP = 10 | BNST I | H = 9.561077, p = 0.008391 | AGG vs NON; U = 5.000000, p = 0.004079<br>AGG vs OBS; U = 25.000000, p = 0.121281<br>OBS vs NON; U = 3.000000, p = 0.004079 |
| | NON = 6, OBS = 7, EXP = 9 | PAG I | H = 1.808905, p = 0.404763 | n.s. |
| | NON = 6, OBS = 7, EXP = 12 | vLS I | H = 7.522857, p = 0.023251 | AGG vs NON; U = 10.000000, p = 0.026934<br>AGG vs OBS; U = 29.000000, p = 0.199413<br>OBS vs NON; U = 6.000000, p = 0.034965 |

| | | | | | |
|---|---|---|---|---|---|
| | | NON = 6, OBS = 7, EXP = 11 | PrL I | F = 2.946660, p = 0.074479 | n.s. |
| | | NON = 6, OBS = 7, EXP = 11 | NAc DA | H = 3.883238, p = 0.143471 | n.s. |
| Fig E4d | DA during nonsocial behavior | NON = 6, OBS = 7, EXP = 11 | One-way ANOVA tests or Kruskal Wallis H tests | | Post hoc independent samples t tests or Mann-Whitney U tests with Benjamini-Hochberg correction |
| | | | NAc DA | F = 7.979767, p = 0.002643 | AGG vs NON; t = -3.541284, p = 0.002962<br>AGG vs OBS; t = -0.197122, p = 0.282071<br>OBS vs NON; t = -2.776569, p = 0.009007 |
| Fig E4e | Preference in hard fight | | One-way ANOVA tests or Kruskal Wallis H tests | | |
| | Nonsocial versus aggression | NON = 6, OBS = 7, EXP = 12 | VMHvl E | F = 0.999528, p = 0.384161 | n.s. |
| | | NON = 6, OBS = 7, EXP = 9 | PMv E | F = 1.386185, p = 0.275472 | n.s. |
| | | NON = 6, OBS = 7, EXP = 10 | AH E | H = 1.840994, p = 0.398321 | n.s. |
| | | NON = 6, OBS = 7, EXP = 11 | POA E | F = 1.279489, p = 0.298992 | n.s. |
| | | NON = 6, OBS = 7, EXP = 11 | MeA E | F = 0.194388, p = 0.824803 | n.s. |
| | | NON = 6, OBS = 7, EXP = 9 | PA E | F = 3.948172, p = 0.036819 | AGG vs NON; t = 3.755219, p = 0.004807<br>AGG vs OBS; t = 0.125027, p = 0.601520<br>OBS vs NON; t = 1.979652, p = 0.073313 |
| | | NON = 6, OBS = 7, EXP = 10 | BNST E | F = 2.115318, p = 0.146775 | n.s. |
| | | NON = 6, OBS = 7, EXP = 9 | PAG E | F = 0.160675, p = 0.852777 | n.s. |
| | | NON = 6, OBS = 7, EXP = 12 | vLS E | F = 1.841521, p = 0.182196 | n.s. |
| | | NON = 6, OBS = 7, EXP = 11 | PrL E | F = 0.915547, p = 0.415692 | n.s. |
| | | NON = 6, OBS = 7, EXP = 9 | LHb E | H = 0.102195, p = 0.950186 | n.s. |
| | | NON = 6, OBS = 7, EXP = 12 | VMHvl I | H = 1.006813, p = 0.604468 | n.s. |
| | | NON = 6, OBS = 7, EXP = 9 | PMv I | F = 0.281985, p = 0.757556 | n.s. |
| | | NON = 6, OBS = 7, EXP = 10 | AH I | H = 2.556522, p = 0.278521 | n.s. |
| | | NON = 6, OBS = 7, EXP = 11 | POA I | H = 2.596364, p = 0.273028 | n.s. |
| | | NON = 6, OBS = 7, EXP = 11 | MeA I | F = 3.029559, p = 0.069825 | n.s. |
| | | NON = 6, OBS = 7, EXP = 9 | PA I | F = 0.468736, p = 0.632839 | n.s. |
| | | NON = 6, OBS = 7, EXP = 10 | BNST I | H = 3.921429, p = 0.140758 | n.s. |
| | | NON = 6, OBS = 7, EXP = 9 | PAG I | F = 1.308811, p = 0.294649 | n.s. |
| | | NON = 6, OBS = 7, EXP = 12 | vLS I | F = 2.128823, p = 0.142835 | n.s. |
| | | NON = 6, OBS = 7, EXP = 11 | PrL I | F = 1.380661, p = 0.273314 | n.s. |
| | | NON = 6, OBS = 7, EXP = 11 | NAc DA | F = 4.273251, p = 0.027732 | AGG vs NON; t = 1.466639, p = 0.163125<br>AGG vs OBS; t = -1.928050, p = 0.107684 |

| | | | | | |
|---|---|---|---|---|---|
| | | | | | OBS vs NON; t = 2.546220, p = 0.081532 |
| | Nonsocial versus aggression | NON = 6, OBS = 7, EXP = 12 | VMHvl E | F = 1.560931, p = 0.232316 | n.s. |
| | | NON = 6, OBS = 7, EXP = 9 | PMv E | F = 2.482633, p = 0.111637 | n.s. |
| | | NON = 6, OBS = 7, EXP = 10 | AH E | F = 0.588706, p = 0.564379 | n.s. |
| | | NON = 6, OBS = 7, EXP = 11 | POA E | F = 2.552614, p = 0.101780 | n.s. |
| | | NON = 6, OBS = 7, EXP = 11 | MeA E | F = 0.657703, p = 0.528387 | n.s. |
| | | NON = 6, OBS = 7, EXP = 9 | PA E | F = 5.390785, p = 0.013985 | AGG vs NON; t = 4.047139, p = 0.002768<br>AGG vs OBS; t = 0.057070, p = 0.636864<br>OBS vs NON; t = 2.276767, p = 0.043784 |
| | | NON = 6, OBS = 7, EXP = 10 | BNST E | H = 7.019772, p = 0.029900 | AGG vs NON; U = 54.000000, p = 0.014985<br>AGG vs OBS; U = 52.000000, p = 0.108803<br>OBS vs NON; U = 25.000000, p = 0.418803 |
| | | NON = 6, OBS = 7, EXP = 9 | PAG E | F = 0.039873, p = 0.960997 | n.s. |
| | | NON = 6, OBS = 7, EXP = 12 | vLS E | F = 1.669709, p = 0.211294 | n.s. |
| | | NON = 6, OBS = 7, EXP = 11 | PrL E | F = 0.445131, p = 0.646647 | n.s. |
| | | NON = 6, OBS = 7, EXP = 9 | LHb E | F = 1.338232, p = 0.285936 | n.s. |
| | | NON = 6, OBS = 7, EXP = 12 | VMHvl I | H = 0.593407, p = 0.743265 | n.s. |
| | | NON = 6, OBS = 7, EXP = 9 | PMv I | F = 0.195203, p = 0.824386 | n.s. |
| | | NON = 6, OBS = 7, EXP = 10 | AH I | H = 2.827640, p = 0.243212 | n.s. |
| | | NON = 6, OBS = 7, EXP = 11 | POA I | H = 4.291948, p = 0.116954 | n.s. |
| | | NON = 6, OBS = 7, EXP = 11 | MeA I | F = 2.381120, p = 0.116943 | n.s. |
| | | NON = 6, OBS = 7, EXP = 9 | PA I | F = 0.332226, p = 0.721409 | n.s. |
| | | NON = 6, OBS = 7, EXP = 10 | BNST I | F = 0.368251, p = 0.696538 | n.s. |
| | | NON = 6, OBS = 7, EXP = 9 | PAG I | F = 0.886272, p = 0.429427 | n.s. |
| | | NON = 6, OBS = 7, EXP = 12 | vLS I | F = 2.466633, p = 0.108012 | n.s. |
| | | NON = 6, OBS = 7, EXP = 11 | PrL I | F = 0.653968, p = 0.530248 | n.s. |
| | | NON = 6, OBS = 7, EXP = 11 | NAc DA | F = 8.155229, p = 0.002394 | AGG vs NON; t = 2.370374, p = 0.047400<br>AGG vs OBS; t = -2.144043, p = 0.047736<br>OBS vs NON; t = 4.078646, p = 0.005473 |
| Fig E5a | Within-group average tuning curves for each region and feature | | *Related samples t-tests (t) 'greater'* | | |

| | | |
|---|---|---|
| EXP vs OBS: 12 EXP r2 with EXP weights vs OBS weights | VMHvl E | res-intr inter-centroid dist: t = -1.715, p = 0.943<br>res-intr inter-head dist: t = -1.852, p = 0.955<br>res-intr head-tailbase dist: t = -1.728, p = 0.944<br>res centroid speed: t = -2.304, p = 0.979<br>intr centroid speed: t = -2.304, p = 0.979<br>res-intr head-head angle: t = -3.382, p = 0.997<br>res body length: t = -3.187, p = 0.996 |
| OBS vs NON: 7 OBS with OBS weights vs NON weights | VMHvl E | res-intr inter-centroid dist: t = -0.066, p = 0.525<br>res-intr inter-head dist: t = -0.075, p = 0.529<br>res-intr head-tailbase dist: t = -0.208, p = 0.579<br>res centroid speed: t = 0.886, p = 0.205<br>intr centroid speed: t = -0.373, p = 0.639<br>res-intr head-head angle: t = -2.239, p = 0.967<br>res body length: t = -2.068, p = 0.958 |
| EXP vs NON: 12 EXP r2 with EXP weights vs NON weights | VMHvl E | res-intr inter-centroid dist: t = -1.756, p = 0.947<br>res-intr inter-head dist: t = -1.88, p = 0.957<br>res-intr head-tailbase dist: t = -1.772, p = 0.948<br>res centroid speed: t = -2.281, p = 0.978<br>intr centroid speed: t = -2.284, p = 0.978<br>res-intr head-head angle: t = -2.991, p = 0.994<br>res body length: t = -2.811, p = 0.992 |
| EXP vs OBS: 12 EXP r2 with EXP weights vs OBS weights | PMv E | res-intr inter-centroid dist: t = 5.975, p = 0.0<br>res-intr inter-head dist: t = 4.605, p = 0.001<br>res-intr head-tailbase dist: t = 2.922, p = 0.011<br>res centroid speed: t = -0.771, p = 0.767<br>intr centroid speed: t = -0.69, p = 0.744<br>res-intr head-head angle: t = -0.827, p = 0.782<br>res body length: t = -0.625, p = 0.724 |
| OBS vs NON: 7 OBS with OBS weights vs NON weights | PMv E | res-intr inter-centroid dist: t = 0.516, p = 0.312<br>res-intr inter-head dist: t = 1.204, p = 0.137<br>res-intr head-tailbase dist: t = -0.201, p = 0.576<br>res centroid speed: t = -1.298, p = 0.879<br>intr centroid speed: t = -0.934, p = 0.807<br>res-intr head-head angle: t = -1.564, p = 0.916<br>res body length: t = -1.647, p = 0.925 |

| | | |
|---|---|---|
| EXP vs NON: 12 EXP r2 with EXP weights vs NON weights | PMv E | res-intr inter-centroid dist: t = 6.384, p = 0.0<br>res-intr inter-head dist: t = 4.07, p = 0.002<br>res-intr head-tailbase dist: t = 3.344, p = 0.006<br>res centroid speed: t = 1.299, p = 0.118<br>intr centroid speed: t = 0.673, p = 0.261<br>res-intr head-head angle: t = 0.591, p = 0.286<br>res body length: t = 0.59, p = 0.287 |
| EXP vs OBS: 12 EXP r2 with EXP weights vs OBS weights | AH E | res-intr inter-centroid dist: t = 2.01, p = 0.038<br>res-intr inter-head dist: t = 1.902, p = 0.045<br>res-intr head-tailbase dist: t = 2.016, p = 0.037<br>res centroid speed: t = 1.286, p = 0.115<br>intr centroid speed: t = 1.227, p = 0.125<br>res-intr head-head angle: t = 1.231, p = 0.125<br>res body length: t = 1.3, p = 0.113 |
| OBS vs NON: 7 OBS with OBS weights vs NON weights | AH E | res-intr inter-centroid dist: t = 1.713, p = 0.069<br>res-intr inter-head dist: t = 2.201, p = 0.035<br>res-intr head-tailbase dist: t = 2.442, p = 0.025<br>res centroid speed: t = 1.914, p = 0.052<br>intr centroid speed: t = 2.004, p = 0.046<br>res-intr head-head angle: t = 2.011, p = 0.046<br>res body length: t = 2.004, p = 0.046 |
| EXP vs NON: 12 EXP r2 with EXP weights vs NON weights | AH E | res-intr inter-centroid dist: t = 2.392, p = 0.02<br>res-intr inter-head dist: t = 2.341, p = 0.022<br>res-intr head-tailbase dist: t = 2.401, p = 0.02<br>res centroid speed: t = 2.128, p = 0.031<br>intr centroid speed: t = 2.099, p = 0.033<br>res-intr head-head angle: t = 2.103, p = 0.032<br>res body length: t = 2.133, p = 0.031 |
| EXP vs OBS: 12 EXP r2 with EXP weights vs OBS weights | POA E | res-intr inter-centroid dist: t = 0.753, p = 0.234<br>res-intr inter-head dist: t = 1.854, p = 0.047<br>res-intr head-tailbase dist: t = 0.989, p = 0.173<br>res centroid speed: t = 1.66, p = 0.064<br>intr centroid speed: t = 1.715, p = 0.059<br>res-intr head-head angle: t = 1.716, p = 0.058<br>res body length: t = 1.713, p = 0.059 |

| | | |
|---|---|---|
| OBS vs NON: 7 OBS with OBS weights vs NON weights | POA E | res-intr inter-centroid dist: t = -0.122, p = 0.547<br>res-intr inter-head dist: t = -0.023, p = 0.509<br>res-intr head-tailbase dist: t = -0.437, p = 0.661<br>res centroid speed: t = -0.761, p = 0.762<br>intr centroid speed: t = -0.898, p = 0.798<br>res-intr head-head angle: t = -0.642, p = 0.728<br>res body length: t = -0.8, p = 0.773 |
| EXP vs NON: 12 EXP r2 with EXP weights vs NON weights | POA E | res-intr inter-centroid dist: t = -0.408, p = 0.654<br>res-intr inter-head dist: t = -0.27, p = 0.604<br>res-intr head-tailbase dist: t = -0.543, p = 0.7<br>res centroid speed: t = -0.999, p = 0.829<br>intr centroid speed: t = 0.721, p = 0.244<br>res-intr head-head angle: t = 0.674, p = 0.258<br>res body length: t = 0.574, p = 0.289 |
| EXP vs OBS: 12 EXP r2 with EXP weights vs OBS weights | MeA E | res-intr inter-centroid dist: t = -0.226, p = 0.587<br>res-intr inter-head dist: t = -0.256, p = 0.598<br>res-intr head-tailbase dist: t = 0.158, p = 0.439<br>res centroid speed: t = -0.978, p = 0.824<br>intr centroid speed: t = -0.973, p = 0.823<br>res-intr head-head angle: t = -1.08, p = 0.847<br>res body length: t = -1.028, p = 0.836 |
| OBS vs NON: 7 OBS with OBS weights vs NON weights | MeA E | res-intr inter-centroid dist: t = -0.691, p = 0.742<br>res-intr inter-head dist: t = -0.99, p = 0.82<br>res-intr head-tailbase dist: t = -0.295, p = 0.611<br>res centroid speed: t = -1.397, p = 0.894<br>intr centroid speed: t = -1.433, p = 0.899<br>res-intr head-head angle: t = -1.766, p = 0.936<br>res body length: t = -1.583, p = 0.918 |
| EXP vs NON: 12 EXP r2 with EXP weights vs NON weights | MeA E | res-intr inter-centroid dist: t = -0.075, p = 0.529<br>res-intr inter-head dist: t = -0.027, p = 0.51<br>res-intr head-tailbase dist: t = 0.263, p = 0.399<br>res centroid speed: t = -0.699, p = 0.75<br>intr centroid speed: t = -0.597, p = 0.718<br>res-intr head-head angle: t = -0.682, p = 0.745<br>res body length: t = -0.626, p = 0.727 |

| | | |
|---|---|---|
| EXP vs OBS: 12 EXP r2 with EXP weights vs OBS weights | PA E | res-intr inter-centroid dist: t = 1.172, p = 0.137<br>res-intr inter-head dist: t = 0.876, p = 0.203<br>res-intr head-tailbase dist: t = 0.85, p = 0.21<br>res centroid speed: t = -0.232, p = 0.589<br>intr centroid speed: t = -0.244, p = 0.593<br>res-intr head-head angle: t = -0.271, p = 0.603<br>res body length: t = -0.154, p = 0.559 |
| OBS vs NON: 7 OBS with OBS weights vs NON weights | PA E | res-intr inter-centroid dist: t = 1.94, p = 0.05<br>res-intr inter-head dist: t = 2.081, p = 0.041<br>res-intr head-tailbase dist: t = -0.291, p = 0.609<br>res centroid speed: t = 1.367, p = 0.11<br>intr centroid speed: t = 1.376, p = 0.109<br>res-intr head-head angle: t = 1.38, p = 0.108<br>res body length: t = 1.341, p = 0.114 |
| EXP vs NON: 12 EXP r2 with EXP weights vs NON weights | PA E | res-intr inter-centroid dist: t = 1.497, p = 0.086<br>res-intr inter-head dist: t = 1.398, p = 0.1<br>res-intr head-tailbase dist: t = 1.032, p = 0.166<br>res centroid speed: t = -0.581, p = 0.711<br>intr centroid speed: t = -0.585, p = 0.713<br>res-intr head-head angle: t = -0.62, p = 0.724<br>res body length: t = -0.382, p = 0.644 |
| EXP vs OBS: 12 EXP r2 with EXP weights vs OBS weights | BNST E | res-intr inter-centroid dist: t = -1.752, p = 0.943<br>res-intr inter-head dist: t = -1.842, p = 0.951<br>res-intr head-tailbase dist: t = -1.61, p = 0.929<br>res centroid speed: t = -2.341, p = 0.978<br>intr centroid speed: t = -2.363, p = 0.979<br>res-intr head-head angle: t = -2.392, p = 0.98<br>res body length: t = -2.204, p = 0.973 |
| OBS vs NON: 7 OBS with OBS weights vs NON weights | BNST E | res-intr inter-centroid dist: t = -1.044, p = 0.832<br>res-intr inter-head dist: t = -1.19, p = 0.861<br>res-intr head-tailbase dist: t = -1.066, p = 0.836<br>res centroid speed: t = -0.027, p = 0.51<br>intr centroid speed: t = -2.461, p = 0.975<br>res-intr head-head angle: t = -2.444, p = 0.975<br>res body length: t = -2.547, p = 0.978 |

| | | | |
|---|---|---|---|
| | EXP vs NON: 12 EXP r2 with EXP weights vs NON weights | BNST E | res-intr inter-centroid dist: t = -1.729, p = 0.941<br>res-intr inter-head dist: t = -1.818, p = 0.949<br>res-intr head-tailbase dist: t = -1.582, p = 0.926<br>res centroid speed: t = -2.375, p = 0.979<br>intr centroid speed: t = -2.402, p = 0.98<br>res-intr head-head angle: t = -2.431, p = 0.981<br>res body length: t = -2.217, p = 0.973 |
| | EXP vs OBS: 12 EXP r2 with EXP weights vs OBS weights | PAG E | res-intr inter-centroid dist: t = -0.475, p = 0.675<br>res-intr inter-head dist: t = -0.22, p = 0.584<br>res-intr head-tailbase dist: t = -0.296, p = 0.612<br>res centroid speed: t = -0.611, p = 0.72<br>intr centroid speed: t = 1.167, p = 0.141<br>res-intr head-head angle: t = -0.5, p = 0.684<br>res body length: t = -0.703, p = 0.748 |
| | OBS vs NON: 7 OBS with OBS weights vs NON weights | PAG E | res-intr inter-centroid dist: t = 1.04, p = 0.169<br>res-intr inter-head dist: t = 1.534, p = 0.088<br>res-intr head-tailbase dist: t = 1.33, p = 0.116<br>res centroid speed: t = 1.633, p = 0.077<br>intr centroid speed: t = 1.604, p = 0.08<br>res-intr head-head angle: t = 1.577, p = 0.083<br>res body length: t = 1.695, p = 0.071 |
| | EXP vs NON: 12 EXP r2 with EXP weights vs NON weights | PAG E | res-intr inter-centroid dist: t = 2.11, p = 0.036<br>res-intr inter-head dist: t = 2.01, p = 0.042<br>res-intr head-tailbase dist: t = 1.741, p = 0.063<br>res centroid speed: t = 1.526, p = 0.085<br>intr centroid speed: t = 2.271, p = 0.029<br>res-intr head-head angle: t = 1.939, p = 0.047<br>res body length: t = 1.843, p = 0.054 |
| | EXP vs OBS: 12 EXP r2 with EXP weights vs OBS weights | vLS E | res-intr inter-centroid dist: t = 3.585, p = 0.002<br>res-intr inter-head dist: t = 3.581, p = 0.002<br>res-intr head-tailbase dist: t = 3.538, p = 0.002<br>res centroid speed: t = 3.012, p = 0.006<br>intr centroid speed: t = 2.978, p = 0.006<br>res-intr head-head angle: t = 3.047, p = 0.006<br>res body length: t = 3.116, p = 0.005 |

| | | |
|---|---|---|
| OBS vs NON: 7 OBS with OBS weights vs NON weights | vLS E | res-intr inter-centroid dist: t = 1.694, p = 0.071<br>res-intr inter-head dist: t = 1.874, p = 0.055<br>res-intr head-tailbase dist: t = 1.435, p = 0.101<br>res centroid speed: t = 1.065, p = 0.164<br>intr centroid speed: t = 1.156, p = 0.146<br>res-intr head-head angle: t = 1.251, p = 0.129<br>res body length: t = 1.243, p = 0.13 |
| EXP vs NON: 12 EXP r2 with EXP weights vs NON weights | vLS E | res-intr inter-centroid dist: t = 4.568, p = 0.0<br>res-intr inter-head dist: t = 4.57, p = 0.0<br>res-intr head-tailbase dist: t = 4.548, p = 0.0<br>res centroid speed: t = 4.38, p = 0.001<br>intr centroid speed: t = 4.382, p = 0.001<br>res-intr head-head angle: t = 4.406, p = 0.001<br>res body length: t = 4.457, p = 0.0 |
| EXP vs OBS: 12 EXP r2 with EXP weights vs OBS weights | PrL E | res-intr inter-centroid dist: t = -0.701, p = 0.75<br>res-intr inter-head dist: t = -1.004, p = 0.83<br>res-intr head-tailbase dist: t = -0.568, p = 0.709<br>res centroid speed: t = -0.195, p = 0.575<br>intr centroid speed: t = -1.264, p = 0.882<br>res-intr head-head angle: t = -1.676, p = 0.938<br>res body length: t = -1.961, p = 0.961 |
| OBS vs NON: 7 OBS with OBS weights vs NON weights | PrL E | res-intr inter-centroid dist: t = 2.868, p = 0.014<br>res-intr inter-head dist: t = 2.835, p = 0.015<br>res-intr head-tailbase dist: t = 2.498, p = 0.023<br>res centroid speed: t = 2.243, p = 0.033<br>intr centroid speed: t = 2.303, p = 0.03<br>res-intr head-head angle: t = 1.751, p = 0.065<br>res body length: t = 1.684, p = 0.072 |
| EXP vs NON: 12 EXP r2 with EXP weights vs NON weights | PrL E | res-intr inter-centroid dist: t = 1.996, p = 0.037<br>res-intr inter-head dist: t = 1.941, p = 0.04<br>res-intr head-tailbase dist: t = 1.984, p = 0.038<br>res centroid speed: t = 1.835, p = 0.048<br>intr centroid speed: t = 1.845, p = 0.047<br>res-intr head-head angle: t = 1.805, p = 0.051<br>res body length: t = 1.677, p = 0.062 |

| | | |
|---|---|---|
| EXP vs OBS: 12 EXP r2 with EXP weights vs OBS weights | LHb E | res-intr inter-centroid dist: t = 1.907, p = 0.046<br>res-intr inter-head dist: t = 1.915, p = 0.046<br>res-intr head-tailbase dist: t = 1.861, p = 0.05<br>res centroid speed: t = 1.724, p = 0.062<br>intr centroid speed: t = 2.042, p = 0.038<br>res-intr head-head angle: t = 2.045, p = 0.038<br>res body length: t = 1.928, p = 0.045 |
| OBS vs NON: 7 OBS with OBS weights vs NON weights | LHb E | res-intr inter-centroid dist: t = -0.033, p = 0.513<br>res-intr inter-head dist: t = 0.073, p = 0.472<br>res-intr head-tailbase dist: t = -0.241, p = 0.591<br>res centroid speed: t = -0.692, p = 0.743<br>intr centroid speed: t = -1.589, p = 0.918<br>res-intr head-head angle: t = -2.016, p = 0.955<br>res body length: t = -2.188, p = 0.964 |
| EXP vs NON: 12 EXP r2 with EXP weights vs NON weights | LHb E | res-intr inter-centroid dist: t = 1.65, p = 0.069<br>res-intr inter-head dist: t = 1.638, p = 0.07<br>res-intr head-tailbase dist: t = 1.616, p = 0.072<br>res centroid speed: t = 1.242, p = 0.125<br>intr centroid speed: t = 1.757, p = 0.059<br>res-intr head-head angle: t = 1.754, p = 0.059<br>res body length: t = 1.619, p = 0.072 |
| EXP vs OBS: 12 EXP r2 with EXP weights vs OBS weights | VMHvl I | res-intr inter-centroid dist: t = 2.989, p = 0.006<br>res-intr inter-head dist: t = 2.905, p = 0.007<br>res-intr head-tailbase dist: t = 3.093, p = 0.005<br>res centroid speed: t = 2.82, p = 0.008<br>intr centroid speed: t = 2.766, p = 0.009<br>res-intr head-head angle: t = 2.739, p = 0.01<br>res body length: t = 2.687, p = 0.011 |
| OBS vs NON: 7 OBS with OBS weights vs NON weights | VMHvl I | res-intr inter-centroid dist: t = -0.849, p = 0.786<br>res-intr inter-head dist: t = -0.86, p = 0.789<br>res-intr head-tailbase dist: t = -0.914, p = 0.802<br>res centroid speed: t = -1.122, p = 0.848<br>intr centroid speed: t = -1.108, p = 0.845<br>res-intr head-head angle: t = -1.096, p = 0.842<br>res body length: t = -1.117, p = 0.847 |

| | | |
|---|---|---|
| EXP vs NON: 12 EXP r2 with EXP weights vs NON weights | VMHvl I | res-intr inter-centroid dist: t = 2.884, p = 0.007<br>res-intr inter-head dist: t = 2.719, p = 0.01<br>res-intr head-tailbase dist: t = 3.124, p = 0.005<br>res centroid speed: t = 2.533, p = 0.014<br>intr centroid speed: t = 2.445, p = 0.016<br>res-intr head-head angle: t = 2.407, p = 0.017<br>res body length: t = 2.324, p = 0.02 |
| EXP vs OBS: 12 EXP r2 with EXP weights vs OBS weights | PMv I | res-intr inter-centroid dist: t = 0.384, p = 0.356<br>res-intr inter-head dist: t = 0.906, p = 0.198<br>res-intr head-tailbase dist: t = 0.518, p = 0.31<br>res centroid speed: t = -1.114, p = 0.849<br>intr centroid speed: t = -2.65, p = 0.984<br>res-intr head-head angle: t = -2.138, p = 0.965<br>res body length: t = -2.296, p = 0.972 |
| OBS vs NON: 7 OBS with OBS weights vs NON weights | PMv I | res-intr inter-centroid dist: t = 2.164, p = 0.037<br>res-intr inter-head dist: t = 1.558, p = 0.085<br>res-intr head-tailbase dist: t = 2.426, p = 0.026<br>res centroid speed: t = 1.222, p = 0.134<br>intr centroid speed: t = 1.229, p = 0.133<br>res-intr head-head angle: t = 1.258, p = 0.128<br>res body length: t = 1.237, p = 0.131 |
| EXP vs NON: 12 EXP r2 with EXP weights vs NON weights | PMv I | res-intr inter-centroid dist: t = 3.797, p = 0.003<br>res-intr inter-head dist: t = 2.939, p = 0.011<br>res-intr head-tailbase dist: t = 3.442, p = 0.005<br>res centroid speed: t = 2.83, p = 0.013<br>intr centroid speed: t = 2.4, p = 0.024<br>res-intr head-head angle: t = 2.505, p = 0.02<br>res body length: t = 2.498, p = 0.021 |
| EXP vs OBS: 12 EXP r2 with EXP weights vs OBS weights | AH I | res-intr inter-centroid dist: t = -0.235, p = 0.59<br>res-intr inter-head dist: t = -0.339, p = 0.629<br>res-intr head-tailbase dist: t = -0.445, p = 0.667<br>res centroid speed: t = -0.654, p = 0.735<br>intr centroid speed: t = -1.028, p = 0.835<br>res-intr head-head angle: t = -1.032, p = 0.835<br>res body length: t = -1.068, p = 0.843 |

| | | |
|---|---|---|
| OBS vs NON: 7 OBS with OBS weights vs NON weights | AH I | res-intr inter-centroid dist: t = 0.458, p = 0.331<br>res-intr inter-head dist: t = 0.723, p = 0.249<br>res-intr head-tailbase dist: t = 1.431, p = 0.101<br>res centroid speed: t = 0.585, p = 0.29<br>intr centroid speed: t = 0.727, p = 0.247<br>res-intr head-head angle: t = 0.617, p = 0.28<br>res body length: t = 0.594, p = 0.287 |
| EXP vs NON: 12 EXP r2 with EXP weights vs NON weights | AH I | res-intr inter-centroid dist: t = 2.306, p = 0.023<br>res-intr inter-head dist: t = 2.264, p = 0.025<br>res-intr head-tailbase dist: t = 2.123, p = 0.031<br>res centroid speed: t = 1.904, p = 0.045<br>intr centroid speed: t = 1.661, p = 0.066<br>res-intr head-head angle: t = 1.703, p = 0.061<br>res body length: t = 1.685, p = 0.063 |
| EXP vs OBS: 12 EXP r2 with EXP weights vs OBS weights | POA I | res-intr inter-centroid dist: t = -0.79, p = 0.776<br>res-intr inter-head dist: t = -0.878, p = 0.8<br>res-intr head-tailbase dist: t = -0.75, p = 0.765<br>res centroid speed: t = -1.159, p = 0.863<br>intr centroid speed: t = -2.008, p = 0.964<br>res-intr head-head angle: t = -2.164, p = 0.972<br>res body length: t = -2.065, p = 0.967 |
| OBS vs NON: 7 OBS with OBS weights vs NON weights | POA I | res-intr inter-centroid dist: t = -1.058, p = 0.835<br>res-intr inter-head dist: t = -1.57, p = 0.916<br>res-intr head-tailbase dist: t = -1.119, p = 0.847<br>res centroid speed: t = -1.235, p = 0.869<br>intr centroid speed: t = -1.346, p = 0.887<br>res-intr head-head angle: t = -1.874, p = 0.945<br>res body length: t = -1.948, p = 0.95 |
| EXP vs NON: 12 EXP r2 with EXP weights vs NON weights | POA I | res-intr inter-centroid dist: t = -0.792, p = 0.777<br>res-intr inter-head dist: t = -0.87, p = 0.798<br>res-intr head-tailbase dist: t = -0.756, p = 0.766<br>res centroid speed: t = -1.117, p = 0.855<br>intr centroid speed: t = -1.707, p = 0.941<br>res-intr head-head angle: t = -1.835, p = 0.952<br>res body length: t = -1.778, p = 0.947 |

| | | |
|---|---|---|
| EXP vs OBS: 12 EXP r2 with EXP weights vs OBS weights | MeA I | res-intr inter-centroid dist: t = 0.151, p = 0.442<br>res-intr inter-head dist: t = -0.881, p = 0.801<br>res-intr head-tailbase dist: t = 0.106, p = 0.459<br>res centroid speed: t = -1.52, p = 0.92<br>intr centroid speed: t = -2.274, p = 0.977<br>res-intr head-head angle: t = -2.968, p = 0.993<br>res body length: t = -2.464, p = 0.983 |
| OBS vs NON: 7 OBS with OBS weights vs NON weights | MeA I | res-intr inter-centroid dist: t = 1.838, p = 0.058<br>res-intr inter-head dist: t = 1.578, p = 0.083<br>res-intr head-tailbase dist: t = 1.054, p = 0.166<br>res centroid speed: t = 0.987, p = 0.181<br>intr centroid speed: t = 1.131, p = 0.151<br>res-intr head-head angle: t = 1.177, p = 0.142<br>res body length: t = 1.22, p = 0.134 |
| EXP vs NON: 12 EXP r2 with EXP weights vs NON weights | MeA I | res-intr inter-centroid dist: t = 1.828, p = 0.049<br>res-intr inter-head dist: t = 1.833, p = 0.048<br>res-intr head-tailbase dist: t = 1.949, p = 0.04<br>res centroid speed: t = 1.874, p = 0.045<br>intr centroid speed: t = 1.912, p = 0.042<br>res-intr head-head angle: t = 1.836, p = 0.048<br>res body length: t = 1.903, p = 0.043 |
| EXP vs OBS: 12 EXP r2 with EXP weights vs OBS weights | PA I | res-intr inter-centroid dist: t = 0.653, p = 0.266<br>res-intr inter-head dist: t = -0.161, p = 0.562<br>res-intr head-tailbase dist: t = 0.516, p = 0.31<br>res centroid speed: t = -0.852, p = 0.791<br>intr centroid speed: t = -1.306, p = 0.886<br>res-intr head-head angle: t = -1.073, p = 0.843<br>res body length: t = -0.795, p = 0.775 |
| OBS vs NON: 7 OBS with OBS weights vs NON weights | PA I | res-intr inter-centroid dist: t = -0.191, p = 0.573<br>res-intr inter-head dist: t = -0.045, p = 0.517<br>res-intr head-tailbase dist: t = -0.577, p = 0.708<br>res centroid speed: t = -2.353, p = 0.972<br>intr centroid speed: t = -3.012, p = 0.988<br>res-intr head-head angle: t = -1.535, p = 0.912<br>res body length: t = -2.134, p = 0.962 |

| | | |
|---|---|---|
| EXP vs NON: 12 EXP r2 with EXP weights vs NON weights | PA I | res-intr inter-centroid dist: t = 1.397, p = 0.1<br>res-intr inter-head dist: t = 0.999, p = 0.174<br>res-intr head-tailbase dist: t = 1.254, p = 0.123<br>res centroid speed: t = 1.063, p = 0.159<br>intr centroid speed: t = 0.889, p = 0.2<br>res-intr head-head angle: t = 0.976, p = 0.179<br>res body length: t = 1.033, p = 0.166 |
| EXP vs OBS: 12 EXP r2 with EXP weights vs OBS weights | BNST I | res-intr inter-centroid dist: t = -1.007, p = 0.83<br>res-intr inter-head dist: t = -1.054, p = 0.84<br>res-intr head-tailbase dist: t = -0.903, p = 0.805<br>res centroid speed: t = -1.338, p = 0.893<br>intr centroid speed: t = -1.346, p = 0.894<br>res-intr head-head angle: t = -1.546, p = 0.922<br>res body length: t = -1.547, p = 0.922 |
| OBS vs NON: 7 OBS with OBS weights vs NON weights | BNST I | res-intr inter-centroid dist: t = 2.553, p = 0.022<br>res-intr inter-head dist: t = 2.352, p = 0.028<br>res-intr head-tailbase dist: t = 2.414, p = 0.026<br>res centroid speed: t = 2.177, p = 0.036<br>intr centroid speed: t = 2.16, p = 0.037<br>res-intr head-head angle: t = 2.152, p = 0.037<br>res body length: t = 2.152, p = 0.037 |
| EXP vs NON: 12 EXP r2 with EXP weights vs NON weights | BNST I | res-intr inter-centroid dist: t = 2.283, p = 0.024<br>res-intr inter-head dist: t = 2.265, p = 0.025<br>res-intr head-tailbase dist: t = 2.333, p = 0.022<br>res centroid speed: t = 1.951, p = 0.041<br>intr centroid speed: t = 2.146, p = 0.03<br>res-intr head-head angle: t = 2.091, p = 0.033<br>res body length: t = 2.094, p = 0.033 |
| EXP vs OBS: 12 EXP r2 with EXP weights vs OBS weights | PAG I | res-intr inter-centroid dist: t = -1.174, p = 0.861<br>res-intr inter-head dist: t = -1.17, p = 0.86<br>res-intr head-tailbase dist: t = -1.211, p = 0.867<br>res centroid speed: t = -1.242, p = 0.873<br>intr centroid speed: t = -1.108, p = 0.848<br>res-intr head-head angle: t = -1.225, p = 0.87<br>res body length: t = -1.182, p = 0.862 |

| | | |
|---|---|---|
| OBS vs NON: 7 OBS with OBS weights vs NON weights | PAG I | res-intr inter-centroid dist: t = 1.847, p = 0.057<br>res-intr inter-head dist: t = 1.79, p = 0.062<br>res-intr head-tailbase dist: t = 1.58, p = 0.083<br>res centroid speed: t = 1.683, p = 0.072<br>intr centroid speed: t = 1.592, p = 0.081<br>res-intr head-head angle: t = 1.649, p = 0.075<br>res body length: t = 1.533, p = 0.088 |
| EXP vs NON: 12 EXP r2 with EXP weights vs NON weights | PAG I | res-intr inter-centroid dist: t = -1.513, p = 0.913<br>res-intr inter-head dist: t = -1.547, p = 0.917<br>res-intr head-tailbase dist: t = -1.618, p = 0.925<br>res centroid speed: t = -1.535, p = 0.916<br>intr centroid speed: t = -1.538, p = 0.916<br>res-intr head-head angle: t = -1.641, p = 0.928<br>res body length: t = -1.619, p = 0.925 |
| EXP vs OBS: 12 EXP r2 with EXP weights vs OBS weights | vLS I | res-intr inter-centroid dist: t = -1.637, p = 0.935<br>res-intr inter-head dist: t = -1.482, p = 0.917<br>res-intr head-tailbase dist: t = -1.68, p = 0.939<br>res centroid speed: t = -1.548, p = 0.925<br>intr centroid speed: t = -1.565, p = 0.927<br>res-intr head-head angle: t = -1.498, p = 0.919<br>res body length: t = -1.544, p = 0.925 |
| OBS vs NON: 7 OBS with OBS weights vs NON weights | vLS I | res-intr inter-centroid dist: t = 1.869, p = 0.055<br>res-intr inter-head dist: t = 2.005, p = 0.046<br>res-intr head-tailbase dist: t = 1.492, p = 0.093<br>res centroid speed: t = 1.579, p = 0.083<br>intr centroid speed: t = 1.633, p = 0.077<br>res-intr head-head angle: t = 1.651, p = 0.075<br>res body length: t = 1.659, p = 0.074 |
| EXP vs NON: 12 EXP r2 with EXP weights vs NON weights | vLS I | res-intr inter-centroid dist: t = -2.108, p = 0.971<br>res-intr inter-head dist: t = -1.617, p = 0.933<br>res-intr head-tailbase dist: t = -1.91, p = 0.959<br>res centroid speed: t = -1.569, p = 0.928<br>intr centroid speed: t = -1.666, p = 0.938<br>res-intr head-head angle: t = -1.483, p = 0.917<br>res body length: t = -1.712, p = 0.943 |

| | | |
|---|---|---|
| EXP vs OBS: 12 EXP r2 with EXP weights vs OBS weights | PrL I | res-intr inter-centroid dist: t = -2.205, p = 0.974<br>res-intr inter-head dist: t = -2.529, p = 0.985<br>res-intr head-tailbase dist: t = -2.54, p = 0.985<br>res centroid speed: t = 0.107, p = 0.459<br>intr centroid speed: t = -2.925, p = 0.992<br>res-intr head-head angle: t = -3.054, p = 0.994<br>res body length: t = -3.047, p = 0.994 |
| OBS vs NON: 7 OBS with OBS weights vs NON weights | PrL I | res-intr inter-centroid dist: t = -0.526, p = 0.691<br>res-intr inter-head dist: t = -0.569, p = 0.705<br>res-intr head-tailbase dist: t = -0.393, p = 0.646<br>res centroid speed: t = 1.35, p = 0.113<br>intr centroid speed: t = -0.59, p = 0.712<br>res-intr head-head angle: t = -1.544, p = 0.913<br>res body length: t = -1.69, p = 0.929 |
| EXP vs NON: 12 EXP r2 with EXP weights vs NON weights | PrL I | res-intr inter-centroid dist: t = -2.005, p = 0.964<br>res-intr inter-head dist: t = -2.185, p = 0.973<br>res-intr head-tailbase dist: t = -2.342, p = 0.979<br>res centroid speed: t = -0.112, p = 0.543<br>intr centroid speed: t = -2.432, p = 0.982<br>res-intr head-head angle: t = -2.46, p = 0.983<br>res body length: t = -2.555, p = 0.986 |
| EXP vs OBS: 12 EXP r2 with EXP weights vs OBS weights | LHb I | res-intr inter-centroid dist: t = 2.863, p = 0.011<br>res-intr inter-head dist: t = 2.816, p = 0.011<br>res-intr head-tailbase dist: t = 2.761, p = 0.012<br>res centroid speed: t = 2.788, p = 0.012<br>intr centroid speed: t = 2.74, p = 0.013<br>res-intr head-head angle: t = 2.746, p = 0.013<br>res body length: t = 2.715, p = 0.013 |
| OBS vs NON: 7 OBS with OBS weights vs NON weights | LHb I | res-intr inter-centroid dist: t = 0.381, p = 0.358<br>res-intr inter-head dist: t = 0.242, p = 0.408<br>res-intr head-tailbase dist: t = 0.004, p = 0.498<br>res centroid speed: t = -0.954, p = 0.812<br>intr centroid speed: t = -1.545, p = 0.913<br>res-intr head-head angle: t = -1.29, p = 0.878<br>res body length: t = -1.324, p = 0.883 |

| | | | |
|---|---|---|---|
| | EXP vs NON: 12 EXP r2 with EXP weights vs NON weights | LHb I | res-intr inter-centroid dist: t = 2.869, p = 0.01<br>res-intr inter-head dist: t = 2.772, p = 0.012<br>res-intr head-tailbase dist: t = 2.685, p = 0.014<br>res centroid speed: t = 2.337, p = 0.024<br>intr centroid speed: t = 2.616, p = 0.015<br>res-intr head-head angle: t = 2.598, p = 0.016<br>res body length: t = 2.569, p = 0.017 |
| | EXP vs OBS: 12 EXP r2 with EXP weights vs OBS weights | Nac DA | res-intr inter-centroid dist: t = 0.851, p = 0.207<br>res-intr inter-head dist: t = 1.079, p = 0.153<br>res-intr head-tailbase dist: t = 0.811, p = 0.218<br>res centroid speed: t = 0.853, p = 0.207<br>intr centroid speed: t = 0.763, p = 0.231<br>res-intr head-head angle: t = 0.743, p = 0.237<br>res body length: t = 0.761, p = 0.232 |
| | OBS vs NON: 7 OBS with OBS weights vs NON weights | Nac DA | res-intr inter-centroid dist: t = 0.43, p = 0.341<br>res-intr inter-head dist: t = 0.904, p = 0.2<br>res-intr head-tailbase dist: t = 1.252, p = 0.129<br>res centroid speed: t = 1.21, p = 0.136<br>intr centroid speed: t = 1.163, p = 0.144<br>res-intr head-head angle: t = 1.232, p = 0.132<br>res body length: t = 1.287, p = 0.123 |
| | EXP vs NON: 12 EXP r2 with EXP weights vs NON weights | Nac DA | res-intr inter-centroid dist: t = 1.69, p = 0.061<br>res-intr inter-head dist: t = 1.822, p = 0.049<br>res-intr head-tailbase dist: t = 1.699, p = 0.06<br>res centroid speed: t = 1.691, p = 0.061<br>intr centroid speed: t = 1.674, p = 0.063<br>res-intr head-head angle: t = 1.667, p = 0.063<br>res body length: t = 1.663, p = 0.064 |
